# Protein landscape of the chromatin states in the malaria parasite *Plasmodium falciparum*

**DOI:** 10.1101/2025.09.23.678001

**Authors:** Gala Ramón-Zamorano, Sheila Mainye, Jessica Kimmel, Inge de Krijger, Abhishek Kanyal, Vendula Horáčková, Jonas Gockel, Yangyang Guo, Michaela Petter, Max Graser, Andrés Guillén-Samander, Michiel Vermeulen, Tobias Spielmann, Richárd Bártfai

**Affiliations:** Bernhard Nocht Institute for Tropical Medicine, Hamburg, Germany; Department of Molecular Biology, Radboud Institute for Molecular Life Sciences, Radboud University, Nijmegen, the Netherlands; Department of Molecular Biology, Radboud Institute for Molecular Life Sciences, Oncode Institute, Radboud University Nijmegen, Nijmegen, the Netherlands; Mikrobiologisches Institut – Klinische Mikrobiologie, Immunologie und Hygiene, Universitätsklinikum Erlangen, Friedrich-Alexander-Universität (FAU) Erlangen-Nürnberg, Erlangen, Germany; Division of Molecular Genetics, Netherlands Cancer Institute, Amsterdam, Netherlands

**Keywords:** malaria, epigenetics, chromatin, proxiome, mass-spectrometry

## Abstract

Epigenetic regulation is essential for development and adaptation across eukaryotes. However, a comprehensive overview of the molecular framework of chromatin-mediated regulation, particularly in non-model organisms, is lacking. Here, we present a systematic proteomic characterization of the chromatin states in *P. falciparum*, an ancient human pathogen with unique genome composition and epigenetic blueprint. We adapted and systematically employed three proximity-labelling approaches to provide a high-confidence and comprehensive proteome of heterochromatin, euchromatin and (peri)centromeric chromatin comprised of 214 proteins, including both expected and new chromatin components. Characterization of 20 proteins validated our approach and i) uncovered a protein influencing parasite transmission, ii) defined complexes relevant for histone variant exchange and chromatin-RNA interactions and iii) provided evidence for the so far believed to be absent spindle assembly checkpoint and the corresponding Bub1-like kinase. This study hence offers a reference proteome of the chromatin states and a resource to uncover novel chromatin biology.

## INTRODUCTION

Epigenetic regulation increases the functional complexity of DNA by providing mechanism to maintain an ON or OFF state of genes through cell division or even generations, yet plastic enough to enable changes of these states. It is central to numerous biological processes throughout the kingdom of life: from mating type determination in yeast^1^, through controlling cell fate decisions during animal development^2^, to ensuring timely flowering in plants^3^. The gene expression status of individual genes or large chromosomal regions is defined by changes in the chromatin structure and orchestrated by a complex interplay between histone variants/modifications, reader, writer and eraser proteins as well as DNA and non-coding RNAs^4–8^. Based on the specific constellation of these features, chromatin can be categorized into structurally and functionally distinct chromatin states, such as heterochromatin or euchromatin that are central to epigenetic regulation^6,9^. Despite a continuing progress in unveiling chromatin biology, we are far from understanding the full granularity of chromatin states and defining functional interactions between their components. Because most epigenetic studies are focusing on a specific protein, locus or regulatory process, the molecular composition of various chromatin states remains largely unexplored in a truly systematic way. Furthermore, while epigenetic regulatory mechanisms are generally well-conserved, key differences exist between different taxa or even species^10–13^, requiring investigation of these processes at various branches of life.

*Plasmodium falciparum*, the parasite causing malaria, provides an important case to understand the composition and function of chromatin states as epigenetic regulation is central to the pathogen’s success^14–16^ and is a tangible target of drug-based intervention strategies^17^. *P. falciparum* caused 600,000 deaths in 2024 alone^18^ and is the medically most important representative of the Apicomplexa, an ancient eukaryotic phylum with numerous pathogens. The disease is caused by the asexual replication of the parasite inside the red blood cells of infected humans. It is transmitted through the bite of female *Anopheles* mosquitoes wherein sexual reproduction occurs. The remarkable cellular transformation and adaptability of the parasite during the life cycle requires stage-specific gene expression programs^26,27^ and with that, a tight regulation of gene expression and cell fate decisions at developmental forking points such as commitment to sexual development and transmission. Stage-specific gene expression and control of developmental switches are achieved by the combination of transcriptional, epigenetic and posttranscriptional regulatory mechanisms^21^.

The genome of *P. falciparum* contains about 5,500 genes and with its overall AT composition of ∼82%, is rated as the most AT-rich genome known to date, a factor profoundly influencing gene regulation and chromatin organization^22^. This haploid genome consists of 14 individual chromosomes, each featuring the three main chromatin states: heterochromatin, euchromatin and centromeric chromatin. Chromatin conformation, nuclear localization, histone modifications, GC-content and transcriptional activity, are some of the features that define and distinguish each chromatin domain^23^. Although numerous histone marks have been identified in *Plasmodium* parasites^24–26^, functional characterization of the post-translational modifications and their associated enzymatic factors is still ongoing.

Transcriptionally silent heterochromatin covers about 10% of the genome in *P. falciparum* parasites, including chromosomal ends and a few chromosome-internal islands^27^. It is maintained in the nuclear periphery as 3-5 heterochromatic clusters^28,29^. Heterochromatin formation is mediated by deacetylation and trimethylation of lysine 9 on histone 3 (H3K9me3) and consequent binding of heterochromatin protein 1 (HP1)^14,15,30–32^. Furthermore, SETvs/SET2 mediated trimethylation of lysine 36 on histone 3 (H3K36me3) has been implicated in heterochromatin formation^33^. Heterochromatic regions receive extensive attention because they harbour genes (*var*), coding for the major virulence factor of the parasite that is antigenically variant and for which epigenetically controlled, mutually exclusive expression is central to parasite virulence, immune invasion and persistence in the host^34^. Heterochromatic regions also contain other surface antigens and multigene families and are involved in key cell fate decision events during parasite development, such as commitment to gametocytogenesis or formation of dormant liver stages^35,36^. Consequently, rearrangement in heterochromatin organization is a major mediator of phenotypic variability and adaptation of the parasites^20,32,37^. Despite these central functions of heterochromatin in parasite biology, the proteins occupying this domain have not been systematically identified, leaving many fundamental questions about formation and maintenance of these compact chromatin structures unanswered.

Outside of heterochromatic regions, most of the genome is in a transcriptionally permissive, euchromatic state, characterized by the lack of H3K9me3 and different degree of acetylation on various histone tails^38,39^. While the genome of *P. falciparum* is densely populated by genes, still clear distinction can be made between coding and intergenic (regulatory) regions in chromatin organization. A parasite-specific feature in euchromatic intergenic regions is the presence of double-variant nucleosomes containing the histone variants H2A.Z and H2B.Z^40,41^. These very same regions are marked by various histone modifications including, H3K4me3, H3K9ac, H3K18ac, H3K27ac as well as acetylation of H4, H2A.Z and H2B.Z^38,39,42,43^. The level of several of these modifications positively correlate with the transcriptional activity of the downstream gene^38,44^, yet the degree of change at the acetylation level cannot on its own explain stage-specific gene regulation. It is more likely that these histone modifications in conjunction with stage-specific transcription factors from the ApiAP2 family recruit the transcriptional machinery to specific sets of genes^44–47^. However, the intricate interactions that collectively orchestrate gene expression in euchromatic regions are not fully understood.

The smallest, but not any less important, chromatin “domain” is the centromere, which is present once on every chromosome. It is demarcated by a CENH3-occupied 4-4.5 kb long region, encompassing a 2–2.5 kb extremely AT-rich (∼97%) core^48,49^. This AT-rich sequence is essential for the centromeres role in chromatid segregation during mitosis and is a conserved feature of centromeres in the *Plasmodium* lineage^50–52^. Members of the Structural Maintenance of Chromosomes (SMC) protein family typically occupy the pericentromeric regions in eukaryotic cells including *Plasmodium* and higher eukaryotes^53–55^. However, although pericentromeric regions are gene-free and HP1-bound in many other organisms, *Plasmodium*’s pericentromeric regions are not heterochromatinised or gene-free^49^. The parasite’s centromeres are displayed in a single cluster at the nuclear periphery close to the centrosome equivalent, but distinct from heterochromatin clusters^49,53^.

In this study, we adapted and employed various proximity labelling methods to provide a comprehensive catalogue of heterochromatin-, euchromatin- and (peri)centromere-specific proteins in malaria parasites. This permitted us to assign a large set of proteins to specific chromatin regions, including both conserved and parasite-specific proteins. Validation of 20 proteins with unknown function not only confirmed accurate assignment of these proteins but also revealed previously hidden regulatory processes essential for proper gene regulation and cell division of this deadly pathogen.

## RESULTS

### Approaches to identify chromatin-domain-associated proteins

We adapted and compared three different proximity labelling approaches (Figure 1A) by using the well-known heterochromatic factor HP1^14,31^ to identify proteins associated with heterochromatin. The first method, DiQ-BioID^57^, conditionally recruits the biotinylation enzyme BirA*^58^ (fused to FRB) to a FKBP-tagged bait using rapalog-dependent heterodimerization. This enables biotinylation of proteins in proximity to the bait in living parasites and allows distinction between background biotinylation (control, cytoplasmic distribution of biotin ligase) and bait-proximal proteins (rapalog, biotin ligase at target protein). The second approach was a modified version of DiQ-BioID using miniTurbo^59^ instead of BirA*, a more active proximity-biotinylation enzyme, permitting considerably shorter labelling times. This approach is referred to as mT-DiQ-BioID (Figure 1A). The biotinylation enzymes – either N- or C-terminal FRB fusions of BirA* (BirA-N^L^ and BirA-C^L^)^60^, or a C-terminal FRB fusion of miniTurbo (mCherry-FRB-miniTurbo)^61^ – were expressed episomally in a previously established and validated parasite line expressing C-terminally 2xFKBP-GFP-2xFKBP-tagged HP1 from its endogenous genomic locus^60^. As a third method, the recently established ProtA-TurboID methodology^62^ was adapted to *P. falciparum*. In this approach, an antibody directs the biotinylation enzyme to the bait (Figure 1A). We applied this method using a specific PfHP1 antibody^14^ with purified nuclei from wild-type parasites.

**Figure 1.**
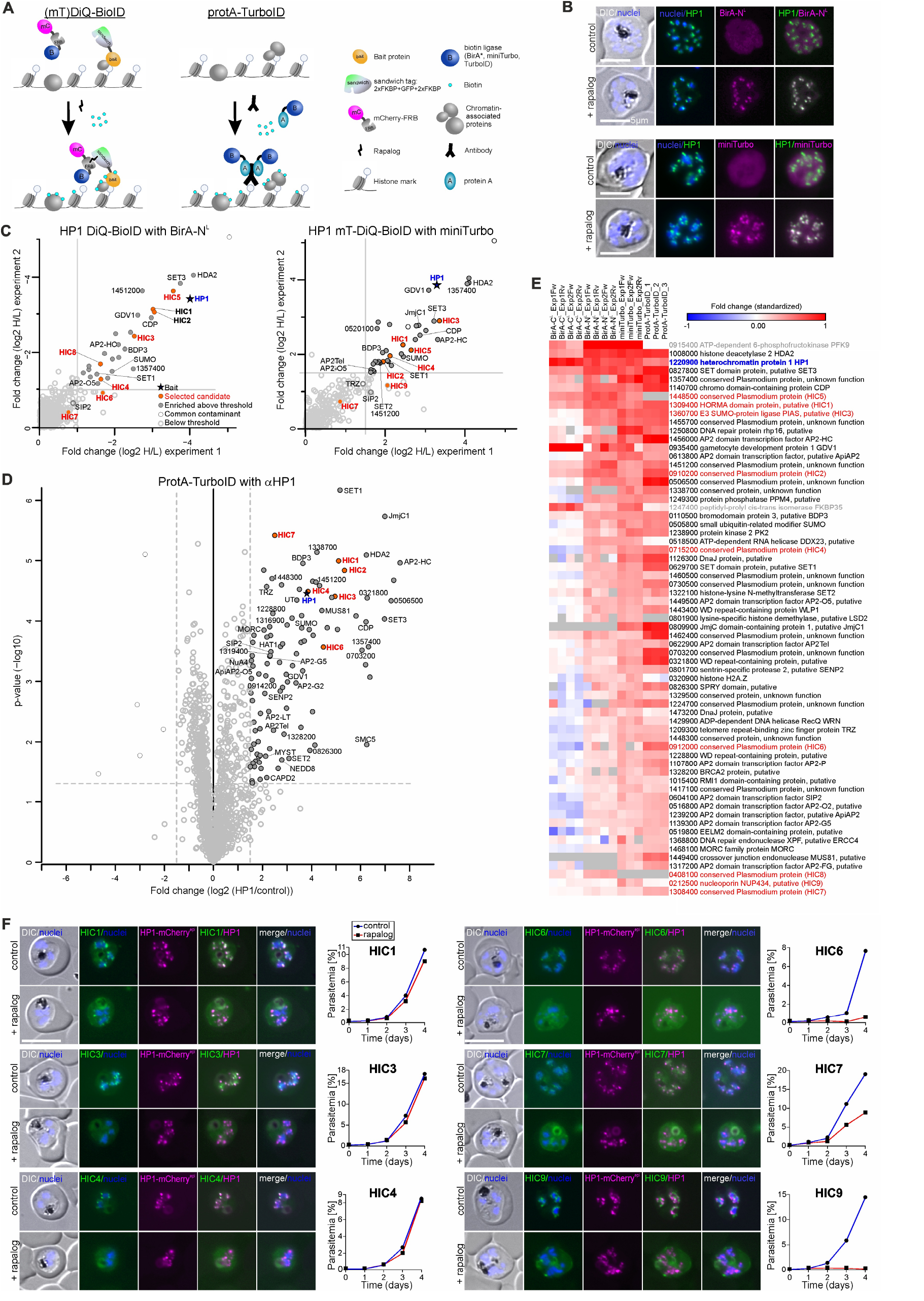
The protein landscape of heterochromatin. **(A)** Schematics of proximity-labelling approaches to target the chromatin-associated proteome. **(B)** Live fluorescence microscopy images showing recruitment of the biotin ligase BirA-NL or mCherry-FRB-miniTurbo (miniTurbo) to HP1 upon rapalog induction (+ rapalog) compared to control (cytoplasmic distribution). Representative images of 2 biological replicates with an average of 20 parasites analyzed per condition and session. See Figure S1A for HP1 BirA-CL images. **(C)** Representative HP1 (mT)-DiQ-BioID experiments using BirA* (left) or miniTurbo (right). Top-right quadrant of scatter plots show log2 fold enrichment of rapalog over control from 2 biological replicates (threshold > 1 (left) or 1.5 (right)). Relevant heterochromatin-associated proteins are marked with their abbreviation (if available) or gene ID (excluding “PF3D7_”). BirA-NL: n = 2 biological replicates including 2 technical replicates each; mCherry-FRB-miniTurbo: n = 2 biological replicates including 1 and 2 technical replicates, respectively. See Figure S1B-D for full plots of all experiments. **(D)** Volcano plot of the HP1-targeting ProtA-TurboID experiment (n = 3 biological replicates). Threshold (dashed lines) log2 enrichment > 1.5 of anti-HP1 over control IgG and p-value < 0.05. Relevant heterochromatin-associated proteins are marked with their abbreviation (if available) or gene ID (excluding “PF3D7_”). Hits coded as indicated in (C). **(E)** Heatmap of top hits detected across all HP1 proximity labelling experiments. Plotted is enrichment (fold change (log2)) standardized to < 1 and > -1 within each replicate. Each HP1 ProtA-TurboID replicate’s values are calculated over the mean of 3 IgG control replicates. Bait (HP1), blue font; HICs, red font; common contaminants, grey font; grey boxes, no value. **(F)** Live fluorescence microscopy images of parasites expressing endogenously tagged HICs and episomally co-expressing HP1-mCherry_Lyn-FRB, showing localization compared to HP1 in parasites grown in absence (control) and presence of rapalog (induced HICs-KS). Representative images of at least 2 independent experiments including microscopy sessions every 24 hours for 2-4 days after KS induction, with an average of 20 parasites analyzed for each condition in each session. The effect of KS of each HIC on asexual parasite growth is shown in line graph next to image panel (one representative out of 3 independent experiments, see Figure S2E for all replicates). Scale bars, 5 μm; DIC, differential interference contrast; nuclei were stained with Hoechst 33342.

The biotinylation enzymes were efficiently recruited to HP1 following the addition of rapalog in all (mT)-DiQ-BioID experiments (Figure 1B), except for the BirA-C^L^, recruitment of which disrupted HP1 localization in most cells (∼70-90%, Figure S1A). Quantitative mass-spectrometric analysis of biotinylated proteins across all approaches identified HP1 as highly enriched compared to control (Figure 1C-D, S1B-D and Table S1), further demonstrating successful recruitment of the biotinylation enzymes to the bait. Moreover, all hit lists included a large number of known and plausible heterochromatic proteins (discussed in the next paragraph), indicating selective biotinylation of HP1-proximal proteins (Figure 1C-D and Table S1). DiQ-BioID with BirA-N^L^ and mT-DiQ-BioID yielded highly similar sets of proteins despite the latter using a considerably shorter rapalog and biotin incubation (30 min versus 24 hours). In contrast, BirA-C^L^, due to interference with HP1 localization, identified only few proteins, likely favoring direct HP1 interactors (such as GDV1, SET3, etc., Figure S1B). It is important to note that this is a bait-specific effect rather than a general limitation of the BirA-C^L^ DiQ-BioID, as it has been successfully applied in previous studies^57,63,64^ and in two other lines in this study (see below). The ProtA-TurboID approach yielded the largest number of (putative) heterochromatin-associated proteins (Figure 1D and Table S1), most of which were also detected by the (mT)-DiQ-BioID methods (Figure 1E).

Collectively these experiments demonstrate that proximity biotinylation is an effective strategy to identify (hetero)-chromatin-associated proteins in *P. falciparum*. Moreover, we highlight the utility of two advanced approaches that offer significant advantages: i) mT-DiQ-BioID enables substantially shorter labeling times than DiQ-BioID, facilitating the detection of, for example, stage-specific interactors; and ii) ProtA-TurboID allows characterization of the proximal proteome (proxiome) for non-tagged baits, such as those in wild-type field isolates or of protein modifications, such as chromatin marks – provided that specific antibodies are available.

### The protein landscape of heterochromatin

To establish a high confidence list of heterochromatin-associated proteins we combined the data from the above experiments using enrichment and significance cut-offs (see Methods for details). Sixty-one proteins were significantly enriched above the set thresholds in more than one type of experiment, including several known heterochromatin-associated factors, such as HDA2, SET3, CDP and GDV1^35,45,65,66^. Somewhat surprisingly, this list revealed a clear overrepresentation of transcription factors belonging to the ApiAP2 family, with 9 out of 26 members detected (Figure 1E). Among these, AP2Tel and SIP2 have previously been shown to bind (sub)telomeric regions, i.e. regions close to heterochromatin^67,68^, while others, such as AP2HC^69^, -O2, -O5 and -G2^70^ have been reported to broadly associate with heterochromatin in ChIP-seq experiments, supporting our finding that these proteins can be found in heterochromatic regions. Certain proteins, including the histone demethylase JmjC1 and the crossover junction endonuclease MUS81, were enriched in both mT-DiQ-BioID experiments and ProtA-TurboID experiments, but not detected using BirA* (Figure 1E and Table S1). This may reflect the more efficient labelling capacity of miniTurbo and ProtA-TurboID^59^.

Among the top 61 proteins, 17 lack functional annotation according to PlasmoDB (https://plasmodb.org)^71^. These include PF3D7_1357400 and PF3D7_1451200, which have also been reported in other studies as HP1-associated^35,45^. From the 61 high-confidence hits, we selected six candidates for validation and further analysis (termed Heterochromatin-Interaction Candidates HIC1 to HIC6). Among these, HIC1 and HIC3 contain annotated putative domains: a HORMA domain^72^ and E3 SUMO-protein ligase domain (PIAS)^73^, respectively. HIC2, HIC4, HIC5 and HIC6 are annotated as unknown. However, structural predictions through HHPred^74^ identified chromatin-related, high probability domain matches for HIC2 (matching to a nucleosome assembly protein) and HIC5 (matching a methyltransferase) (Table S2). To explore lower confidence hits and to find proteins potentially located at the fringes of heterochromatic regions or bridging other chromatin states, we additionally selected proteins significantly enriched in only one of the applied methods for additional validation (HIC7 to HIC9). HIC7 was significantly enriched in ProtA-TurboID and while enriched in the other approaches, it fell below our strict thresholds. HIC8 was identified in BirA*-based DiQ-BioIDs, while HIC9 was detected in mT-DiQ-BioIDs. Structural predictions did not identify functional domains in HIC7, only found coiled-coils in HIC8, and a low confidence match between HIC9 and the human nucleoporin NUP434 (Table S2). In the course of this study, HIC9’s orthologs in *P. berghei* was reported as a nucleoporin^75^ and annotated as NUP434. While nuclear pores and heterochromatin occupy rather distinct regions at the nuclear periphery in *P. falciparum*^76^ (see also Figure 1F, HIC9 panel), interaction of NUPs with heterochromatin has been reported in yeast^77^.

All HICs were endogenously tagged at the C-terminus with the multipurpose 2xFKBP-GFP-2xFKBP (“sandwich”) tag^60^ in *P. falciparum* 3D7 parasites (Figure S2A-B and File S1), with the exception of HIC2, for which we were unable to generate knock-in parasites despite multiple attempts. Seven out of the eight analyzed candidates displayed a nuclear localization, with six showing full or partial co-localization with an episomal HP1-mCherry fusion protein (Figure 1F). While HIC5 was nuclear, it was detectable in only a very low proportion of cells and its uniform distribution in the nucleus indicates it is not concentrated in heterochromatic regions (Figure S2C). HIC8 co-localized with the Golgi apparatus, indicating it is not a nuclear protein (Figure S2D).

To evaluate the functional relevance of the nuclear HICs (apart from HIC5) during parasite blood-stage development, we employed the knock-sideway (KS) system^78^, a strategy previously shown to be effective to achieve conditional functional inactivation for various *P. falciparum* proteins, including HP1 and other nuclear proteins^60,64^. This approach successfully depleted all nuclear HICs from the nucleus, except for HIC1, which remained nuclear despite KS induction (Figure 1F). These experiments showed that depletion of HIC3 and HIC4 from the nucleus did not impact parasite growth, while HIC6, HIC7 and HIC9 were important for asexual growth of the parasite (Figure 1F and S2E). HP1’s localization was not obviously affected after KS of the HICs, indicating that these effects were not due to a gross negative effect on heterochromatin (Figure 1F).

In summary, we present a high-confidence and comprehensive list of heterochromatin-interacting proteins (Table S1), including numerous novel heterochromatic candidates, including unique proteins that may have parasite-specific functions.

### HIC7 influences the commitment to sexual development

Amongst the novel heterochromatin-associated proteins important for asexual growth, HIC7 drew particular interest. Despite extensive comparative searches of its sequence and predicted structure in various databases, no similarities to known proteins were identified. This suggests that HIC7 may have a parasite-specific role in chromatin biology. An in-depth analysis of the relationship of the HIC7-HP1 localization using 3D reconstructed cells revealed that in agreement with its mild enrichment in the HP1-proximity labelling experiments, HIC7 only partially overlapped with HP1 and was located also in regions generally overlapping with the DNA stain (Figure 2A). Despite this wider distribution, focal or elongated accumulations of HIC7 frequently overlapped either fully (Figure 2A, white arrows) or partially (Figure 2A, yellow arrows) with HP1 foci. We occasional also observed HP1 foci without associated or overlapping HIC7 (Figure 2A, cyan arrows). Typically, the partially overlapping HIC7 foci were located on the side of the HP1 foci facing the nuclear lumen (Figure 2A). However, KS of HIC7 had no obvious effect on HP1 localisation (Figure 2A, rapalog).

**Figure 2.**
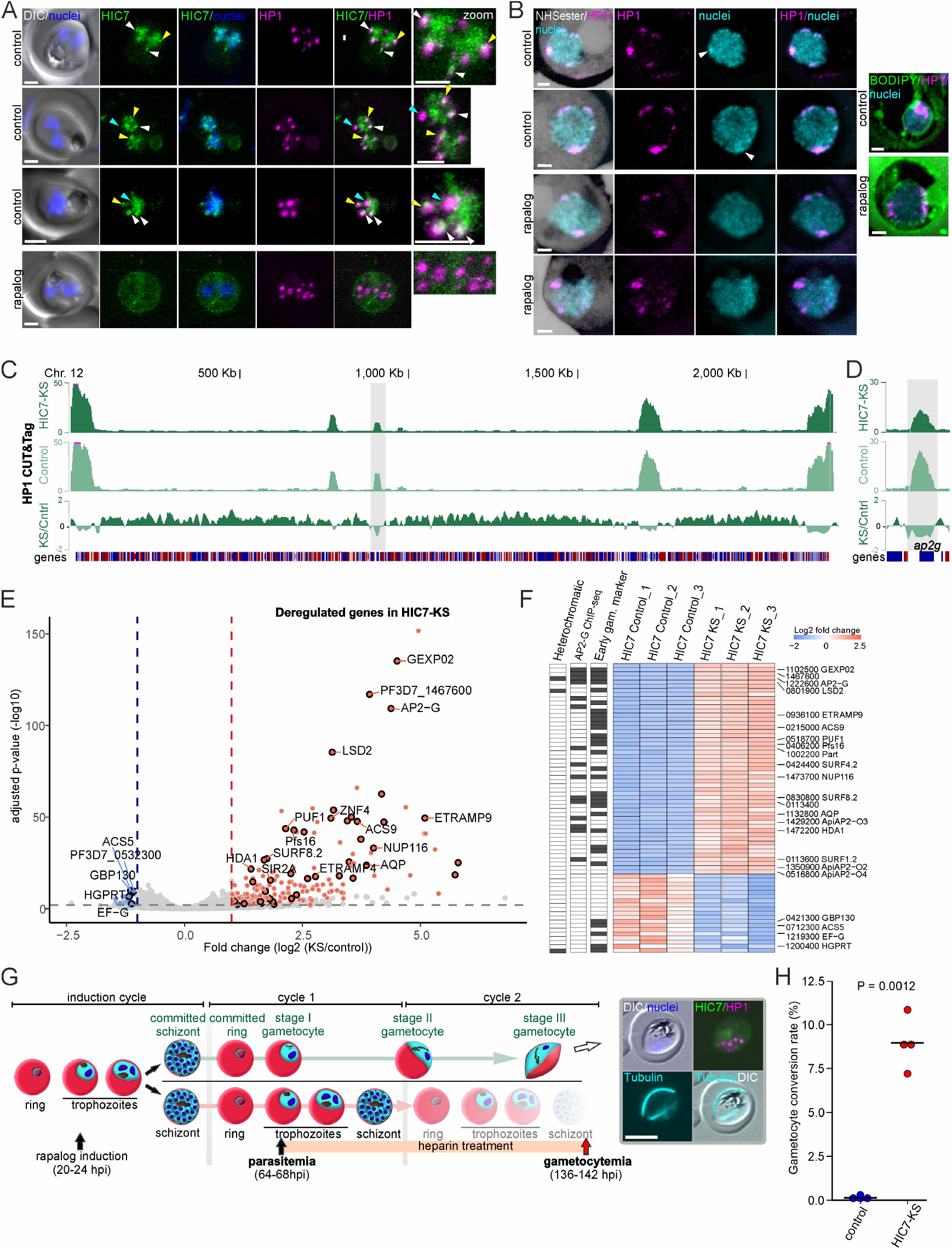
HIC7 influences balance between asexual replication and sexual development. **(A)** Live confocal microscopy images of parasites expressing endogenously tagged HIC7 and episomally co-expressing HP1-mCherry_Lyn-FRB grown in absence (control) and presence of rapalog (induced HIC7-KS). Representative maximum intensity projections from two independent microscopy sessions with at least 10 imaged parasites per session and condition. **(B)** Expansion Microscopy (U-EM) images of parasites expressing endogenously tagged HIC7 and episomally co-expressing HP1-mCherry_Lyn-FRB grown in absence (control) and presence of rapalog (induced HIC7-KS). Anti-mCherry (for HP1-mCherry), Hoechst 33342 and NHS-ester (left panel) or Bodipy (right panel). Images are maximum intensity projections of Z-slices (control: 10-21, 7-15 (left), 12-22 (right); + rapalog: 9-16, 10-19 (left), 21-29 (right)) and representatives from 5 (left panel) and 3 (right panel) imaging sessions with at least 5 imaged parasites per session and condition. **(C)** CUT&Tag tag counts (upper two tracks) and KS over control (KS/Cntrl) log2 ratio tracks (lowest track) of HP1 occupancy over the entire chromosome 12 in HIC7-KS (KS) and control parasites (control). One representative out of 2 independent experiments is shown (see Figure S3C for all replicates). The genomic locus of *ap2-g* is highlighted in grey. **(D)** Zoomed-in view of the *ap2-g* locus from (C). **(E)** Volcano plot of RNA-seq experiments (n = 3 replicates) showing up- (red) and downregulated (blue) genes in late-ring-stage HIC7-KS parasites grown in presence of rapalog (KS) over control (grown without rapalog). Coloured dots represent genes with a base mean expression >= 30, fold change (log2) >= 1 and adjusted p-value <= 0.01. Early gametocyte-specific genes found in Dogga et *al*.^87^ are highlighted with a black border. Hits are marked with their abbreviation (if available) or gene ID. **(F)** Heatmap of the top 51 upregulated and all downregulated genes from (E). Plotted are enrichment values (fold change (log2)) standardized to < 2.5 and > 2 within each replicate. Presence (grey) or absence (white) of these genes in datasets from heterochromatic genes^42^, an early gametocyte-specific marker^87^ and/or an AP2-G target gene in sexually committed rings^88^ is shown on the left. Gene IDs (excluding “PF3D7_”) and abbreviations (if available) of genes are indicated. **(G)** Schematics showing experimental design for gametocyte conversion rate calculation. Timepoints for HIC7-KS induction (rapalog induction) and heparin treatment, as well as parasitemia and gametocytemia determination are indicated. Representative images of tubulin tracker Deep Red stained stage III gametocyte after HIC7-KS as indicated in the scheme and quantified in (H). **(H)** Quantification of gametocyte conversion rate in HIC7-KS parasites (KS) and control parasites (without rapalog addition) from n = 4 biological replicates. Mean and t-test p-value are indicated. Scale bars, 2 μm (1 μm for zoom images); DIC, differential interference contrast; nuclei were stained with Hoechst 33342; hpi, hours post invasion.

To gain insights into HIC7 function using a guilt-by-association approach, we performed proximity labelling with HIC7 (Figure S3A). DiQ-BioID experiments consistently identified HIC7 itself along with a small set of proteins (Figure S3B and Table S3). Several of these were also identified in the HP1 proxiome (Table S1), reinforcing the link between HIC7 and heterochromatin. However, the identified interactors did not provide obvious clues about HIC7’s molecular function.

Given the HIC7-HP1 association, we next investigated whether the growth defect observed upon HIC7 knocksideways could be attributed to alterations in heterochromatin organization. As fluorescence microscopy (Figure 1F and 2A) did not indicate any gross changes after HIC7KS, we used expansion microscopy to assess HP1 in HIC7-KS parasites. In the control, HP1 was detected in several foci located at the fringes of the DNA signal including foci seemingly in contact with the nuclear envelope (Figure 2B control), consistent with its expected location based on fluorescence microscopy^28,29^. Of note, regions of higher Hoechst signal intensity contained HP1, suggesting that heterochromatic regions are apparent in Hoechst-stained expanded cells. After HIC7-KS, the distribution pattern of HP1 did not visibly change (Figure 2B, rapalog), confirming the interpretation from the live fluorescence imaging speaking against global alterations of heterochromatin organisation. To obtain an even more fine-grained picture of heterochromatin organization during HIC7-KS, we profiled genome-wide HP1 occupancy using CUT&Tag^61^. Induction of HIC7-KS in synchronized ring-stage parasites did not result in major heterochromatin reorganization, at either schizont or the subsequent ring stages (Figure 2C and S3C) but showed a modest, global reduction in HP1 occupancy specifically at the ring stage (Figure 2C and S3C, ratio track of KS over control) with an even more pronounced loss of HP1 occupancy over the *ap2-g* locus (Figure 2D). This finding suggests a potential role for HIC7 in maintaining silencing of the gametocyte master transcription factor^65,79^ which is a key sexual commitment regulator.

To understand the impact of HIC7 depletion on gene expression, we investigated transcriptional changes at the end of the first cycle (schizonts) and the beginning of the second cycle (rings). Comparison of RNA-seq data from control and HIC7-KS (+rapalog) parasites at the schizont stage revealed no significant transcriptional alterations (Table S4). However, in ring stage parasites, we observed significant deregulation of 272 genes (Figure 2E and Table S4), with most of them (253 genes) showing increased mRNA abundance. This pool of upregulated genes was not enriched for heterochromatic genes^32^, suggesting that mild overall reduction in heterochromatin occupancy does not lead to deregulation of multigene families. However, they were clearly enriched for early gametocyte markers^80^ and AP2-G-target genes^81^, including *ap2-g, nup116, gexp2* and *lsd2* (Figure 2F and Table S4), suggesting an increased number of parasites induced to become gametocytes among HIC7-KS parasites.

Given these findings, we investigated whether HIC7-KS, similar to HP1-knock-down parasites^14^, leads to an increased gametocyte commitment rate. To measure this, we induced the HIC7-KS in young trophozoites (induction cycle, Figure 2G), just before sexual commitment typically occurs and measured the number of gametocytes arising from the same culture plus (HIC7-KS) and minus (control) rapalog. Further asexual development was prevented with heparin to determine the rate of parasites turning into gametocytes only from the parasites in cycle 1, i.e. those committing in the induction cycle (scheme, Figure 2G). While control parasites exhibited only a low gametocyte conversion rate below 1%, sexual commitment was significantly increased to ∼8% in the HIC7-KS (Figure 2H). Hence, conditional removal of HIC7 from the nucleus substantially increases gametocytogenesis.

Overall, these experiments identify HIC7 as a protein concentrated at the border and partially overlapping with heterochromatic regions that influences the transmission rate of the parasite by maintaining HP1 occupancy at the *ap2-g* locus.

### The protein landscape of euchromatin

Acetylation on histone tails is a universal epigenetic mark of transcriptionally active chromatin, and this is also true in *P. falciparum*, where these marks are enriched in intergenic regions upstream of (active) euchromatic genes^38,39,42^. To characterize the proteome associated with euchromatin, we performed a ProtA-TurboID experiment using a commercial anti-H3K27ac antibody. Mass spectrometric analysis identified 99 high confidence protein hits (Figure 3A and Table S5; see methods for cutoffs), including five ApiAP2 transcription factors, all known bromodomain-containing proteins (acetyl-lysine binding proteins: BDP1-7, SET1, GCN5), and other known euchromatin associated proteins, such as PHD1 and LSD1^45,82^. This confirmed the efficacy of this approach to target euchromatic intergenic regions in the parasite.

**Figure 3.**
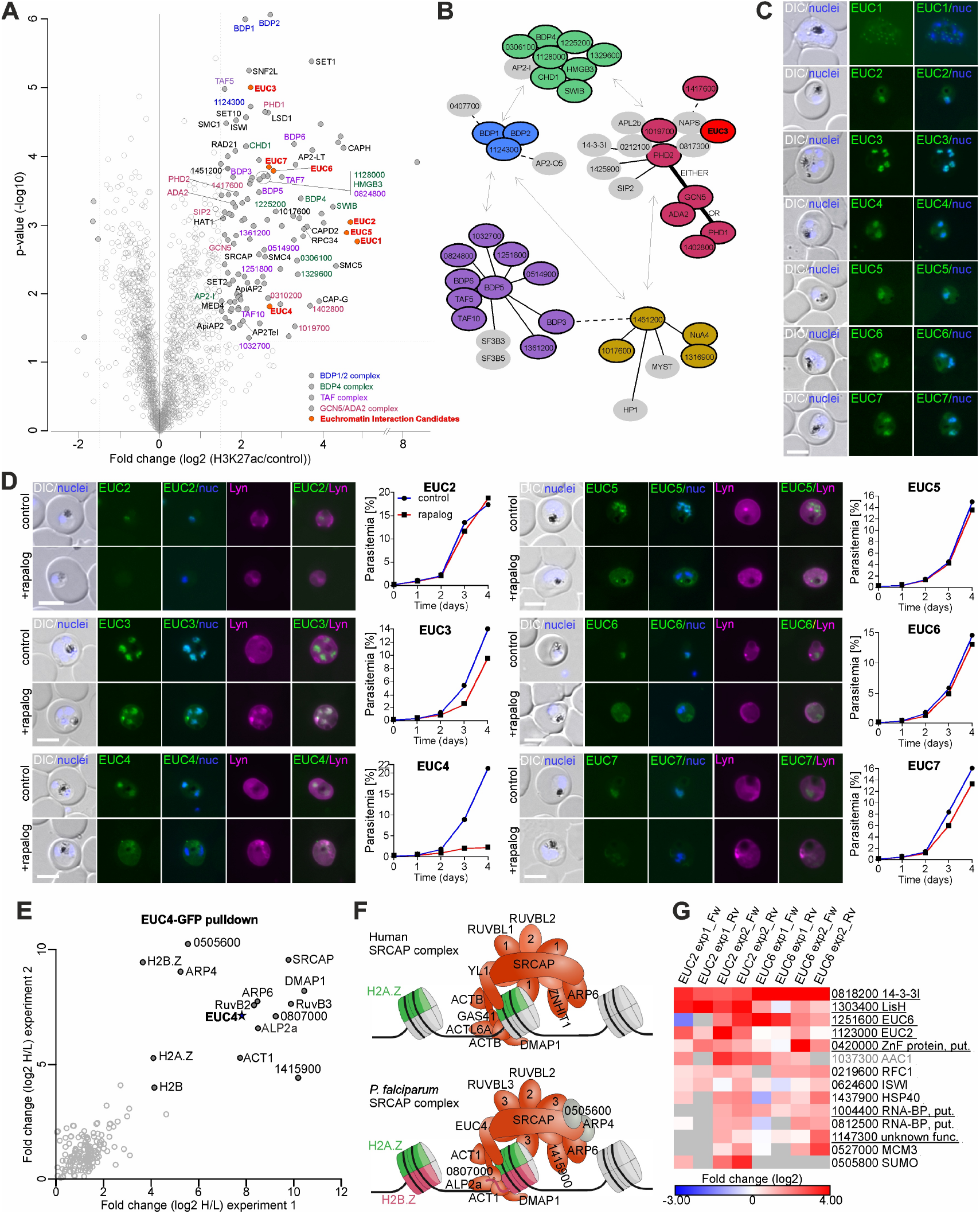
The protein landscape of euchromatin. **(A)** Volcano plot of the H3K27ac-targeting ProtA-TurboID experiment (n = biological 3 replicates). Threshold (dashed lines) log2 enrichment > 1.5 of anti-H3K27ac over control IgG and p-value < 0.05. Relevant euchromatin-associated proteins and euchromatin interaction candidates (EUCs) are marked with their abbreviation (if available) or gene ID (excluding “PF3D7_”). Hits coded as indicated in legend. **(B)** Schematics modified from Hoeijmakers et *al*.^36^ showing the epigenetic reader complexes found in *P. falciparum* colour-coded according to legend from (A). EUC3 is shown associated to GCN5/PHD2 complex, according to interaction data from EUC3 pulldowns from Figure S5B. **(C)** Live fluorescence microscopy images of parasites expressing endogenously tagged EUCs showing their localization in live parasites. Representative images of at least 7 microscopy sessions including an average of 6 fields of view in each. See Figure S4B for full panels. **(D)** Live fluorescence microscopy images of parasites expressing endogenously tagged EUCs and episomally co-expressing Lyn-FRB-mCherry grown in presence of rapalog (induced EUCs-KS) and absence (control). Representative images of at least 4 independent experiments including an average of seven fields of view for each condition. The effect of KS of each EUC on asexual parasite growth is shown in line graph next to image panel (one representative out of 3 independent experiments, see Figure S4C for all replicates). **(E)** Top-right (Figure 3 continued) quadrant of scatterplot of the EUC4-GFP pulldown experiments showing log2 fold enrichment in 2 biological replicates (including 2 technical replicates each heavy vs light label) plotted against each other. Highly enriched proteins in both experiments are marked with their abbreviation (if available) or gene ID (excluding “PF3D7_”). EUC4 as the bait protein is highlighted with a star symbol. **(F)** Schematic comparison of the human SRCAP complex (top) based on structural studies^144,145^ and the proposed model in *P. falciparum* SRCAP complex composition (bottom) from this study. The positions of PF3D7_0505600 and ARP4 in the complex are speculative. **(G)** Heatmap of top hits detected in EUC2 and EUC6 pulldown experiments (see Figure S5C-D for full plots). Plotted are enrichment values (fold change (log2 of GFP/control ratios)) standardized to < 4 and > -3 within each replicate (n = 2 biological and 2 technical replicates). and ribosomal proteins were removed. EUC2/EUC6 complex components are underlined; common GFP-pull-down contaminant (as determined by Hoeijmakers et *al*.^36^), grey font; grey boxes, no value. Scale bars, 5 μm; DIC, differential interference contrast; nuclei (nuc.) were stained with Hoechst 33342.

As a complementary approach, we performed an additional ProtA-TurboID experiment using an antibody against the histone mark H3K4me3, which marks active promoters in eukaryotic cells. Forty-eight proteins passed our thresholds for high confidence hits, of which nearly all (43 proteins) overlapped with the H3K27ac dataset. Several other known euchromatic proteins, although detected, fell just below our enrichment thresholds in the H3K4me3 experiment (Table S5), suggesting a somewhat lower sensitivity with this antibody. Overall, the two datasets were highly concordant, with the anti-H3K27ac antibody yielding a more comprehensive euchromatin proteome. Notably, 32 of the identified euchromatin-associated proteins had been previously assigned to distinct epigenetic complexes^45^ (Figure 3B), and these datasets also include components of the condensin complex (such as SMC4/SMC5, CAPH and CAPD2) and additional proteins with chromatin-associated roles (such as AP2 transcription factors and chromatin remodelling proteins) (Figure 3A and Table S5).

Among the identified proteins, we found 24 without annotated function or reported chromatin association. Of these, seven were selected for further validation and termed Euchromatin-Interaction Candidates: EUC1 to EUC7. All candidates appear in the high confidence lists from both, the H3K27ac and the H3K4me3 experiments, except EUC1, which, while highly enriched, did not meet the cut-off for statistical significance in the H3K4me3 experiment (Table S5). None of the EUCs contain annotated functional domains, except for EUC4, which is annotated as “YL1 nuclear protein, putative” in PlasmoDB. Structural predictions by HHPred^74^ assigned the highest-confidence match to EUC7 (lowest e-value, 100% probability), indicating resemblance to Enhancer of polycomb-like protein (Table S2), which has been confirmed by a parallel study as a component of the NuA4 complex in *Plasmodium*^83^.

Endogenous sandwich-tagging (see Figure S4A and File S1) revealed that six EUCs localize to the nuclei during asexual blood-stage development (Figure 3C and Figure S4B), while EUC1 localized to small foci visible exclusively in late schizonts that were adjacent to the DNA stain but did not evidently overlap with it (Figure 3C). Expression patterns of the EUCs ranged from constitutive across all stages (e.g. EUC3) to stage-specific (e.g. EUC2 and EUC6 which were expressed mainly in tropho zoites and early schizonts) (Figure S4B). The nuclear EUCs were mostly confined to the DNA-stained area of the nucleus where they showed a uniform distribution (Figure 3C and S4B), overall consistent with the diffuse staining patterns observed with histone-acetylation and H3K4me3 antibodies^84^ and clearly differing from the distribution of HICs. An exception was EUC5, which showed focal accumulations in addition to the euchromatin-typical diffuse nuclear distribution (Figure 3C and S4B). Interestingly, this protein was also detected in our heterochromatin proxiome, providing an explanation for the localization pattern and indicating that EUC5 has a dual heterochromatic and euchromatic association.

KS efficiently depleted EUC2, EUC5, EUC6 and EUC7 from the nucleus without impacting parasite growth (Figure 3D and S4C), indicating these proteins are largely dispensable for asexual replication. EUC3 could only be partially removed from the nucleus by KS, but still caused a mild-to-moderate growth defect, suggesting that this protein is potentially important in parasite development. Removal of EUC4 by KS was efficient and led to a severe growth defect (Figure 3D and S4C), indicating an important function in parasite development.

To gain functional insight into EUC4, we performed GFP pull-downs and identified 14 significantly enriched proteins (Figure 3E and Table S6). These interactors showed strong homology to components of the chromatin-remodelling SRCAP complex (Figure 3F and S5A), which, in other eukaryotes, deposits the histone variant H2A.Z^85,86^. We identified *Plasmodium* orthologs for all human SRCAP complex components among EUC4 interactors, along with two parasite-specific proteins (Figure 3E,F and S5A). One of these, ARP4, was previously shown to participate in H2A.Z deposition^87^, strongly supporting the composition and function of this complex. Furthermore, a parallel study provided independent evidence for the existence of this complex through a pull-down using DMAP1, a protein that is however also shared with other complexes in the parasite^83^. Our experiment is, thus, an additional and unequivocal confirmation of the composition of the *Plasmodium* SRCAP/SWR1 complex through the exclusive component EUC4/YL1. Besides H2A.Z, the histone variant H2B.Z was also highly enriched, suggesting that EUC4 and its interactors mediate deposition of the H2A.Z/H2B.Z double-variant nucleosome into euchromatic intergenic regions of the *Plasmodium* genome. Notably, this complex includes a YEATS domain-containing protein (PF3D7_0807000) previously identified as an acetylated H2B.Z histone tail reader^45^, potentially facilitating complex recruitment to euchromatin. Finally, the list of EUC4 interactors also contains the protein PF3D7_0505600, of which the human SRCAP complex lacks an orthologue, but is an orthologue of the Swc5 component of the yeast Swr1 complex, where it is required for the successful histone exchange, most likely by mediating removal of H2A/H2B^88^.

Encouraged by the functional annotation of EUC4, we conducted GFP pull-downs for other EUCs. EUC3 associated with known *P. falciparum* epigenetic complexes^45^, particularly the GCN5/ADA2 complex, and showed weaker, likely indirect interaction with other epigenetic factors, such as the BDP4 and BDP1/BDP2 complexes (Figure 3B, S5B and Table S6). EUC2 and EUC6 co-purified a unique overlapping set of interacting proteins, including each other (Figure 3G, S5C-D and Table S6). Hence, EUC2 and EUC6 are likely part of the same novel complex that includes other factors such as LisH, 14-3-3I, a Zn-finger protein (PF3D7_0420000), an RNA-binding protein (PF3D7_1004400) and PF3D7_1147300, with putative histone/protein- and RNA-binding activities and functions during the trophozoite and early schizont stages when EUC2 and EUC6 are expressed. Given the limited knowledge of RNA-binding proteins in the context of *Plasmodium* chromatin biology, further exploration of this novel complex might provide new insights into for example linking ncRNA to chromatin remodelling or capturing the newly formed RNA for further processing/export.

In summary, proximity biotinylation targeting the H3K27ac and H3K4me3 histone marks enabled identification of a high confidence set of proteins interacting with active promoter regions, including both known and novel components of epigenetic complexes.

### The protein landscape of centromeres

To characterize the centromeric and pericentromeric chromatin we performed proximity biotinylation experiments using CENH3 and SMC1 (Figure 4A). In *P. falciparum* parasites, the histone variant CENH3 has been described as the characteristic component of centromeric chromatin^49^, while the cohesin complex member SMC1 is enriched at pericentromeric regions in trophozoites and schizonts (Grotz, M. et *al*., in preparation). Both proteins were tagged in separate parasite lines, with CENH3 fused to an N-terminal sandwich tag and SMC1 tagged at the C-terminus with GFP (Figure S6A-B and File S1). Fluorescence microscopy of live cells confirmed their expected subnuclear localization, revealing a single focus in each nucleus (or two foci in nuclei undergoing mitosis) closely associated with tubulin at the nuclear periphery for both fusion proteins (Figure 4B,C).

**Figure 4.**
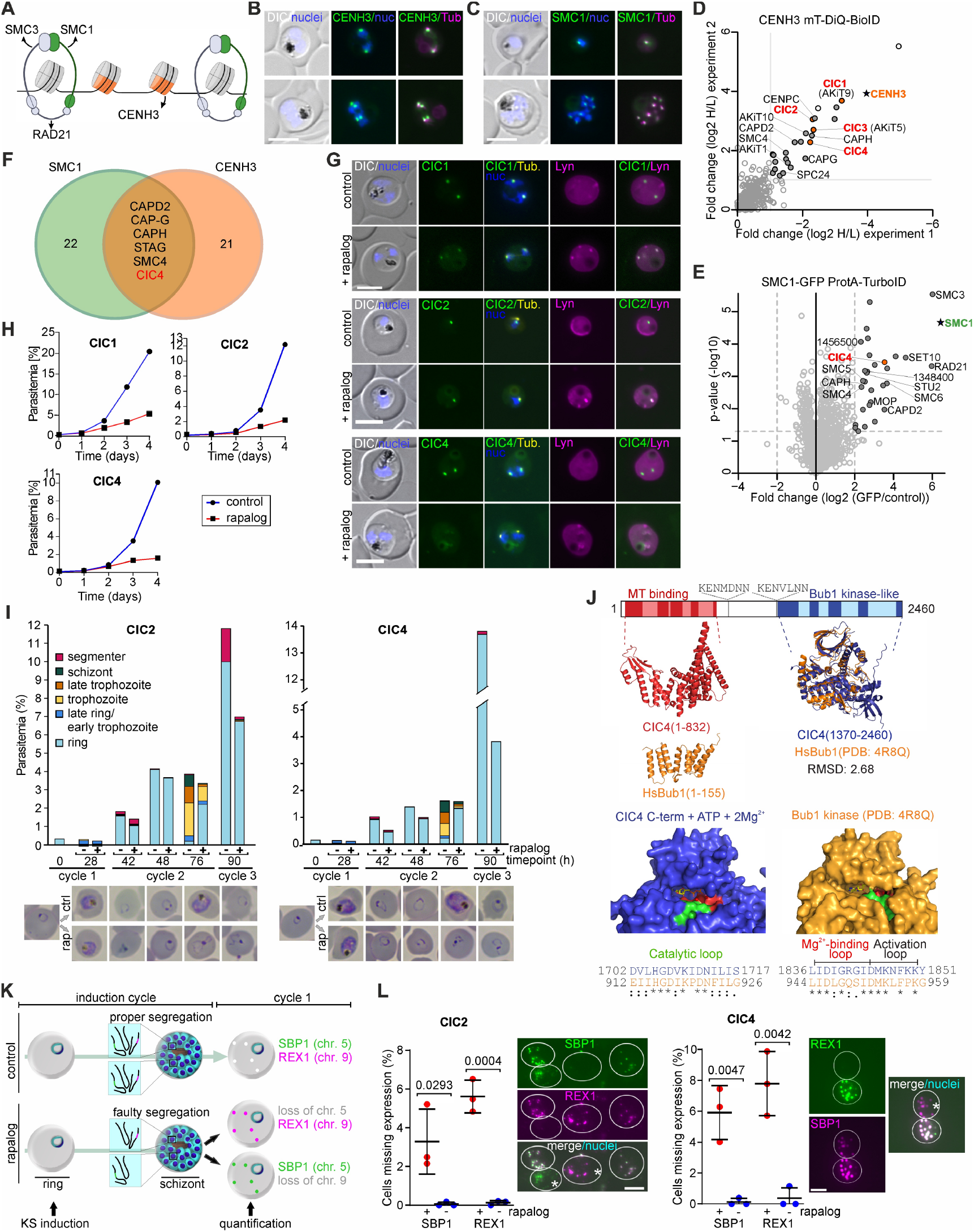
The protein landscape of centromeric and pericentromeric chromatin. **(A)** Schematic of centromeric chromatin (orange, CENH3) with flanking pericentromeric regions associated with the cohesin complex (light blue and green) made with BioRender. **(B-C)** Live fluorescence microscopy images of parasites expressing endogenously taggedCENH3 (B) or SMC1 (C), showing focal distribution of the protein overlapping with tubulin (stained with Tubulin Tracker Deep Red) in live parasites. Representative images shown from 10 (CENH3) or 3 (SMC1) independent microscopy sessions including an average of 7 (CENH3) or 11 (SMC1) fields of view per condition and session. **(D)** Top-right quadrant of scatter plot of the CENH3 mT-DiQ-BioID experiments showing log2 fold change (rapalog over control) of 2 biological replicates (including 2 technical replicates each, heavy vs light label) plotted against each other (threshold > 1). Relevant centromeric chromatin-associated proteins in are marked with their abbreviation (if available) or gene ID (excluding “PF3D7”). Bait (CENH3) is highlighted with a star symbol. See Figure S6H for full plots of all experiments. **(E)** Volcano plot of the SMC1-targeting ProtA-TurboID experiment (n = 3 biological replicates). Threshold (dashed lines) log2 enrichment > 2 of anti-GFP over control IgG and p-value < 0.05. Relevant pericentromere-associated proteins are marked with their abbreviation (if available) or accession number (excluding “PF3D7_”). Bait (SMC1) is highlighted with a star symbol. **(F)** Venn diagram showing the shared proteins from datasets (E, see also Figure S6F, G) of centromeric(CENH3) and pericentromeric- (SMC1) proteins. **(G)** Live fluorescence microscopy images of parasites expressing endogenously tagged CICs and episomally co-expressing Lyn-FRB-mCherry grown in presence of rapalog (induced CICs-KS) and absence (control) and stained with Tubulin Tracker Deep Red. Representative images of at least 5 (CIC1), 4 (CIC2) or 6 (CIC4) independent experiments including an average of 7 fields of view per condition and session. See Figure S7A for full panels. **(H)** Line graphs showing the effect of KS of each CIC on asexual parasite growth (one representative out of 3 independent experiments, see Figure S7B for all replicates). **(I)** Stages and growth based on Giemsa smears (example images shown beneath the graph) in synchronised (0-2 hours synchronisation window) parasites expressing endogenously tagged CIC2 (left) or CIC4 (right) and episomally co-expressing Lyn-FRB-mCherry grown in presence (+ rapalog, to induce CICs-KS) and absence of rapalog (-rapalog) at the indicated time points. Rap, rapalog; ctrl, control. Second biological replicates shown in Figure S7C. **(J)** Schematics of CIC4 (top, not to scale, KEN boxes indicated) and predicted protein structure by alphafold of the N-terminal microtubule (MT) binding domain and the C-terminal Bub1 kinase-like domain. Alignment (by US-align) with human Bub1 protein structure (PDB: 4R8Q) is shown below. Lighter shades of colour indicate flexible loops within folded regions. Surface representations (bottom) of both kinase domains highlight conserved features important for nucleotide and ion-binding. **(K)** Schematics showing experimental design for quantification of missing SBP1 or REX1 signal. Time points for CIC2/CIC4-KS induction and quantification, as well as missing chromosomes due to aneuploidy are indicated. **(L)** Quantification of proportion of cells lacking either SBP1 or REX1 protein expression in CIC2 (top) or CIC4 (bottom) KS parasites (+ rapalog) compared to controls (-rapalog) from n = 3 biological replicates using the assay shown in (K). Mean and t-test p-value are indicated. Example IFA images showing cells missing expression of one of the two antigens. Scale bars, 5 μm; DIC, differential interference contrast; nuclei (nuc.) were stained with Hoechst 33342.

Three different (mT)-DiQ-BioID experiments were conducted with the CENH3-tagged line (using BirA-C^L^, BirAN^L^ and mCherry-FRB-miniTurbo), while ProtA-TurboID experiments targeting SMC1-GFP were performed using an anti-GFP antibody. The high confidence hits (see methods for cut offs) included 27 proteins in the CENH3 proxiome (Figure 4D, S6C-H and Table S7) and 28 proteins in the SMC1 proxiome (Figure 4E and Table S7). Despite the physical proximity of centromeric and pericentromeric chromatin, the proxiomes for the two baits were largely distinct with kinetochore components such as the apicomplexan kinetochore proteins (AKiTs)^89^ specifically detected in the CENH3 vicinity and cohesin complex members being proximal to pericentromeres. Overlapping hits included the condensin I subunits CAPD2, CAP-G, CAP-H and SMC4, a STAG-domain containing protein previously shown to interact with SMC1/SMC3^56,90^, and a putative kinase (PF3D7_ 1112100) (Figure 4F).

From these datasets, we selected four uncharacterized proteins for further validation, hereafter referred to as Centromeric chromatin-Interaction Candidates: CIC1 to CIC4 (Table S7), three of which were identified in the CENH3 proximity proteome and CIC4 which was shared in both datasets. For three of the four candidates (CIC1, CIC2 and CIC4), we successfully produced C-terminally tagged lines (Figure S6I and File S1). All three proteins were found in foci overlapping with nuclear tubulin, confirming a location at centromeric regions (Figure 4G and Figure S7A) and further validating our hit list. Their expression was first detectable from the mid-trophozoite stage on and remained present until the end of the cycle, except for CIC4 which was absent in segmented schizonts (Fig. S7A). Conditional inactivation of either CIC1, CIC2 or CIC4 through KS resulted in a strong growth defect in intraerythrocytic parasites (Figure 4H and Figure S7B), indicating that they are important for asexual replication. However, it is worth noting that nuclear depletion of CIC1 and CIC2 was only partial (Figure 4G) and these experiments might underestimate their importance for parasite growth.

CIC1 and CIC3 have recently been reannotated as the kinetochore components PfAKiT9 and PfAKiT5, respectively, through their homology to *P. berghei* kinetochore proteins^89^. We therefore focussed on the characterization of CIC2 and CIC4. To gauge potential functional roles of these proteins, we determined the developmental stage at which the growth defect occurs upon their nuclear depletion. Both control and parasites with CIC2- or CIC4-KS completed the first 48h cycle and produced almost the same number of new rings (Figure 4I and S7C). However, in the KS parasites, most of these rings failed to develop further, leading to an arrested development in the second cycle while controls continued to grow (Figure 4I and S7C).

Considering that the phenotype manifested with a delay of one developmental cycle and occurred in rings, a stage these proteins are not even expressed, this indicated that mitosis progresses normally in the absence of CIC2 and CIC4 but that a defect occurred impacting the progeny cells. To obtain further hints on their function, we predicted the structure of CIC2 and CIC4 using AlphaFold3^91^. The structural prediction of CIC2 only yielded low-confidence secondary structures, which impeded further comparison to known protein structures. In contrast, the predicted structure of CIC4 allowed similarity searches using foldseek^92^, which identified a high degree of structural similarity to the experimentally solved structure of the human Bub1 kinase (PDB 4R8Q)^93^ in the C-terminal half (best hit). Bub1 is a conserved kinase critical for the spindle assembly checkpoint (SAC)^94^. The C-terminal kinase domain of CIC4 showed both structural and sequence conservation with the HsBub1 (RMSD of 2.68 using US-align^95^), including the nucleotide and ion-binding sites (Figure 4J; iPTM 0.96 for the binding to 2Mg^2+^ ions and 1 ATP molecule in AlphaFold3). The N-terminal domain of CIC4 contains an arrangement of α-helices, a feature also found in the N-terminal TPR domain of HsBub1, which mediates association to the kinetochore, although structures of these domains do not directly align. Furthermore, we detected KEN boxes in CIC4, which in humans serves as anaphase promoting complex recognition signals required for HsBub1 degradation^94^ (Figure 4J). Hence, the structurally highly conserved kinase domain and the presence of additional motifs support the idea that CIC4 may be the *P. falciparum* Bub1 orthologue (henceforth PfBub1-like). The existence of cell cycle check points in *P. falciparum* is controversial^96–99^ but the phenotype of CIC4/PfBub1-like would be consistent with a role similar to proteins of the SAC during mitosis: in yeast, inactivation of Bub1 leads to aneuploidy due to progression of mitosis despite of segregation defects^100^. In the parasite, aneuploidy and associated loss of genes of entire chromosomes in the progeny of the CIC4 KS schizonts would have no effect on the establishment of new rings in RBCs as it can occur without the synthesis of newly synthetised proteins^101^. However, thereafter an aneuploid ring would miss a large number of genes, which would very likely prevent further development.

Aneuploidy is not trivial to detect, as loss or addition of single chromosomes would not significantly alter the total DNA content per nucleus and, as each nucleus can be expected to have different changes, it would not be obvious in a bulk analysis. We therefore resorted to an indirect method to test whether loss of CIC2 and CIC4/Bub1-like leads to aneuploidy on a single-cell level. We chose two exported proteins encoded by genes on two different chromosomes, SBP1 (chromosome 5) and REX1 (chromosome 9). These proteins are expressed from very early ring stage^102^ while the SBP1 and REX1 expressed in the previous cycle is lost with the host cell upon rupture at the end of the previous cycle. Hence, SBP1 and REX1 detected in rings can only arise from newly expressed protein and signifies the presence of the chromosome they are encoded on. Therefore, to test for the presence of these chromosomes in rings after CIC2- or CIC4/Bub1- like-KS, we carried out immunofluorescence assays (IFAs) to detect SBP1 and REX1 in second-cycle rings (see scheme in Figure 4K). For both CIC2 and CIC4 we detected a proportion of cells lacking one or the other antigen (∼ 4-8%, Figure 4L), while such cells were very rare in controls (3 out of 1191 and 4 out of 866 cells for CIC2 and CIC4, respectively). These results support the idea that the delayed onset of the phenotype after CIC2-KS and CIC4/Bub1-like-KS is the result of increased levels of aneuploidy, suggesting a defective SAC.

In summary, we here provide a comprehensive characterization of the rather distinct proteomes of the centromeric and pericentromeric regions. Furthermore, we identify novel centromere associated proteins, including a Bub1-like kinase that could be involved in a so-far-elusive SAC of the malaria parasite.

## DISCUSSION

In this study, we provide a comprehensive catalogue of proteins associated with each chromatin states in *P. falciparum* parasites, namely, heterochromatin, euchromatin and (peri)centromeric chromatin. We obtained high confidence proxiomes by using multiple baits per site and combining different methodologies for proximity labelling. BirA*-based DiQ-BioID is a robust method that has proved highly useful in *Plasmodium* ^57,99,103–106^ and here we further highlight its versality through the systematic survey of proteins associated to distinct chromatin states. By replacing BirA* with miniTurbo^58,59^, we now demonstrate that the mT-DiQ-BioID can label the proxiome of different baits within 30 minutes (and possibly even quicker) and hence improve the method to enable identification of stage-specific or transient interactions. In addition, we also adapted the recently introduced ProtA-turboID^62^ to the malaria parasite, which adds the possibility of profiling the proxiome of non-taggable targets with the use of specific antibodies. Here, we demonstrate its utility to map the proxiome of chromatin-associated proteins (Figure 1D) and posttranslational histone modifications (Figure 3A), but we expect it to be critical to solve limitations of current proximity labelling tools in profiling the proxiome of other non-taggable targets and proteins in, for example, field isolates or *Plasmodium* species that are not genetically tractable. Altogether, we substantially expand the toolbox for analyzing the protein composition of distinct compartments in malaria parasites and our comparisons between the methods, particularly using the same bait, provide guidance for choosing the most suited technology in future studies.

In total, we assigned 214 proteins to specific chromatin domains, highlighting their potential functions in the context of chromatin biology (Figure 5A). Despite the physical proximity of these different domains within the nucleus of about 1 μm in diameter, we identified largely distinct proxiomes for the different chromatin domains (less than 20% of the proteins being shared between any of these domains, Figure 5B) and even between the centromeric and pericentromeric regions (Figure 4F). This highlights the high spatial resolution of our analysis in agreement with DiQ-BioIDs of other kinetochore components, such as the SKA complex^64^. Hence, our approach permits a survey of the distinct composition of these chromatin domains. Interestingly, 14 proteins were detected in both heterochromatin and euchromatin proxiomes (Figure 5B), of which HIC7 and EUC5 indeed showed a localization consistent with both, the heterochromatic and euchromatic nuclear domains (Figures 2A and 3C). Such overlap could be explained by the recruitment of typical euchromatic factors (e.g. SET1, BDP5, MYST) to small euchromatic islands, such as for active antigenic variation genes or promoters of the *var* introns, within the heterochromatic domains^107,108^. Furthermore, we observed several AP2 transcription factors enriched both in the euchromatin and in the heterochromatin (Figure 5B). The presence of these factors in the heterochromatin is somewhat surprising, but since most of these transcription factors are dispensable for intraerythrocytic development^21^, a possible explanation is that they might be tethered to heterochromatin in asexual parasites to prevent undesired binding to their target genes. This would be a unique mechanism to (temporarily) “inactivate” transcription factors that has, to our knowledge, not been described in any organism.

**Figure 5.**
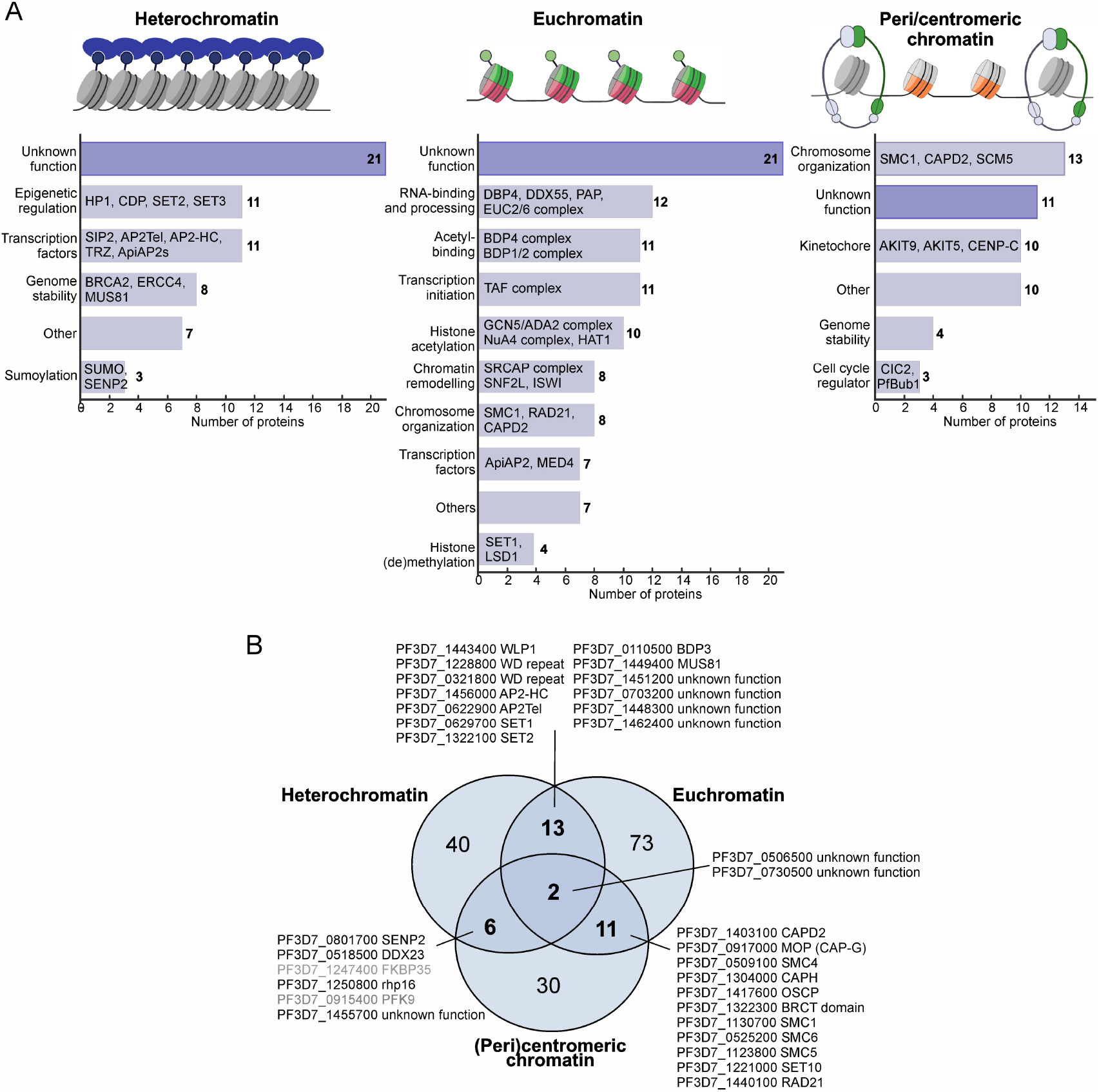
Overview of the protein landscapes of the three chromatin states. **(A)** The relative contribution of proteins to chromatin-associated molecular processes. Chromatin schematics made with BioRender. **(B)** Venn diagram of overlap between high confidence hits of the three chromatin states. Grey font indicates common contaminants of BioID experiments.

Proteins overlapping between the pericentromeric and euchromatic proxiomes (11 proteins, Figure 5B) are enriched for components of the cohesin (e.g. RAD21, SMC1) and condensin (e.g. SMC4, CAPD2) complexes. This is not surprising because, although mostly enriched at pericentromeric regions, cohesins have been shown to have a role in the repression of euchromatic genes as well^56^.

The comprehensive chromatin state catalogue described here provides an opportunity to further investigate the components of each chromatin state and reveal conserved as well as unique parasite-specific chromatin biology (Figure 5A). HIC7 appears to be an interactor of both heterochromatin and euchromatin (Figure 2A and 5B) without any conserved domain or recognizable structure, indicating it belongs to the parasite-specific category. Its inactivation led to a rather specific reduction of heterochromatin over the *ap2-g* locus (Figure 2C), providing a mechanism for the observed increase in gametocyte commitment rates (Figure 2H). While this phenotype is in many ways reminiscent to the overexpression of another heterochromatin-associated, parasite-specific protein, GDV1^35^, these two proteins do not appear to directly interact (Figure S3B). Hence, HIC7 may be a GDV1-independent factor that contributes to the regulatory axis governing the rate of gametocyte commitment and parasite transmission either specifically or through a general effect on heterochromatin (which was hinted at by the general reduction in HP1 occupancy, Figure S3C). The latter option would indicate that HIC7 contributes to the general maintenance of heterochromatin. While HIC7 was found concentrated in HP1-positive areas of the nucleus, it was also found in other DNA-positive but HP1-free regions and in accumulations at the fringe of HP1 foci facing the nuclear centre. Given its non-exclusive co-localization with HP1 it remains to be determined how HIC7 impacts heterochromatin, but one possibility is that it is involved in governing nuclear organization in general.

Euchromatin-associated proteins tend to act in distinct complexes^45^. In fact, many of the proteins identified in the H3K27ac and H3K4me3 proxiomes belong to such complexes (Figure 3B), including euchromatic complexes with broadly conserved function (such as TFIID and GCN5/AGA2 complexes) as well as the parasite-specific BDP1/BDP2 and BDP4 complexes. Yet not all components of these complexes were enriched in the proxiome (grey circles in Figure 3B, e.g. several members of the PHD2-complex) and hence they might represent subcomplexes that are not relevant in the (eu)chromatin context. The novel euchromatin protein candidates identified in this study are also present in various complexes. Their pulldown revealed: i) new components of existing complexes (EUC3, Figure S5B), ii) the composition of the complex responsible for the placement of the parasite-specific H2A.Z and H2B.Z histone pair (EUC4, Figure 3E-F), and iii) a new complex with a likely parasite-specific function, whose role is yet to be characterized in detail (EUC2/6, Figure 3G). Importantly, not all novel euchromatin proximal proteins could be placed to these complexes and hence their analysis will likely reveal novel functions required for gene activation.

We further identified the likely Bub1 checkpoint kinase of the parasite we termed PfBub1-like (Figure 4J), and showed that its inactivation, as well as that of CIC2, leads to a phenotype consistent with aneuploidy (Figure 4K,L), a common consequence of a defective cell cycle check-point during mitosis in eukaryotic cells. The combination of these features provides compelling indication that PfBub1-like kinase regulates mitosis and thereby ensures correct chromatin distribution to daughter cells during schizogony of the malaria parasite. Numerous studies in model organisms have shown that the presence of Bub1 at unattached kinetochores activates the SAC which is a carefully regulated process mediated by interactions of Bub1 with outer kinetochore proteins and multiple check-point proteins (e.g. MAD1)^109^. In contrast, evidence for malaria parasites also relying on SAC regulation during mitosis is only recently emerging. It has been shown that malaria parasites harbour a potential MAD1 homologue^89^ and a paralog of the main cell cycle regulation kinase Aurora B^110,111^, suggesting that the cell cycle regulators proposed to be absent so far might have evaded detection. Our findings of the Bub1-like kinase in the parasite presents a good example of how combination of proteomics, functional and structural analysis can reveal that seemingly parasite-specific proteins may in fact be orthologues of proteins in other organisms, although they were initially not recognized due to sequence divergence. Our analyses of localization and functional characterization suggesting CIC4 to be the Bub1 kinase of the parasite and supports a functional conservation. This is in line with other work wherein localization, interaction and functional data paired with structural predictions revealed evolutionary conservation of proteins of the malaria parasite that were difficult to detect based on the primary sequence level^64,89,112^. Based on this it can be assumed that structural comparisons of the unknown proteins of this resource will reveal further proteins conserved in other organisms and this may help to predict the functions of many more of the *Plasmodium* unknown proteins^113^. Consequently, identifying those that do not have structural similarity to other organisms, we might find truly parasite-specific chromatin-associated factors, which not only reveal exciting new biological processes, but also potential targets for the development of novel compounds with high specificity for the parasite.

## Supporting information

Suppl. file S1. Plasmid sequences

Table S1. HP1 proximity proteome

Table S2. Summary of candidates

Table S3. HIC7 proximity proteome

Table S4. HIC7 KS differential expression

Table S5. Euchromatin proximity proteome

Table S6. EUCs interacting proteins

Table S7. Centromeric proximity proteome

Table S8. Oligonuclotides

## ACKNOWLEDGMENTS

This work was funded by a Leibniz Collaborative Excellence Grant (MalNucFunc; K328/2020 to T.S. and R.B.). The Vermeulen lab is part of the Oncode Institute, which is partly funded by the Dutch Cancer Society. Work in the Vermeulen lab (I.d.K.) was further supported by an ENW-Groot grant (ocenw.groot.2019.017). Y.G. was supported by a Chinese Government Scholarship (CSC). V.H. and T.S. acknowledge funding by the European Research Council (ERC, grant 101021493).

## AUTHOR CONTRIBUTIONS

Conceptualization: G.R.Z., S.M., T.S. and R.B.; investigation: G.R.Z., S.M., J.K., I.K., A.K. V.H., J.G., Y.G., M.G., A.G.S., M.P. and T.S.; methodology: G.R.Z., S.M., J.K., I.K., A.K. V.H. and J.G.; formal analysis: G.R.-Z., S.M., J.K., I.K., A.K. V.H. and J.G.; visualization: G.R.Z., S.M., J.K., A.K., V.H. and J.G.; data curation: G.R.Z., A.K. and J.G.; writing – original draft: G.R.Z. and R.B.; writing – review & editing: J.K., I.K., M.P., A.G.S., M.V. and T.S.; resources: G.R.Z., S.M., I.K., Y.G., M.P. and T.S.; funding acquisition: M.V., T.S and R.B.; supervision: M.V., T.S. and R.B.

## RESOURCE AVAILABILITY

The mass spectrometry proteomics data have been deposited to the ProteomeXchange Consortium via the PRIDE^114^ partner repository. Datasets from different proteomics experiments were deposited separately, but all can be found by the keyword “Pf_Chromatin_proxiome”. The identifiers for each dataset are: PXD065969 for HP1 (mT)-DiQ-BioIDs, PXD066000 for HP1, H3K27an and H3K4me3 ProtA-TurboIDs, PXD066049 for HIC7 DiQBioIDs, PXD066014 for EUC pulldowns, PXD066018 for CENH3 (mT)DiQ-BioIDs, and PXD066044 for SMC1 ProtA-TurboIDs.

Raw and processed RNA-seq and HP1 CUT&Tag dataset of HIC7-KS parasites have been deposited in NCBI’s Gene Expression Omnibus (GEO)^115^ and are accessible through GEO Series accession numbers: GSE305073 and GSE305074.

## SUPPLEMENTAL INFORMATION

*See separate document*.

**Figure S1**. Additional data of HP1 (mT)-DiQ-BioID experiments.

**Figure S2**. Additional data for HICs.

**Figure S3**. Additional data for HIC7.

**Figure S4**. Additional data for EUCs.

**Figure S5**. Interacting proteins of EUC4, EUC3, EUC2 and EUC6.

**Figure S6**. Additional data for SMC1-targeting ProtA-TurboID and CENH3 (mT)-DiQ-BioID experiments.

**Figure S7**. Additional data for CICs.

## METHODS

### Standard parasite culture and synchronization

*P. falciparum* 3D7 parasites^116^ were cultured in human 0+ erythrocytes (RBCs) at 5% haematocrit in RPMI-1640 complete medium containing 0.5% AlbuMAX (Life Technologies) or 10% human serum and 0.2% NaHCO_3_ in an low oxygen atmosphere (3-5% O_2_, 4-5% CO_2_, 90-93% N_2_) at 37 °C^117^. For biotinylation experiments, parasites were grown in modified RPMI with L-glutamine and without biotin and phenol red (US Biological Life Sciences, R9002-01) supplemented with 200 µM Hypoxyxanthine (Merck, H9377) and 0.5% AlbuMAX II (Gibco, 11021037).

Wild type parasites were grown in the absence of antibiotics, whereas genetically modified parasites were cultured in the presence of 2.5 nM WR99210 (Jacobus), 400 μg/ml G418 (Merck), 0.9 μM DSM1 (Sigma-Aldrich) or 2.5 μg/ml BSD (Life Technologies), as detailed below.

When synchronous parasite cultures were required, synchronicity was obtained on ring-stage parasites by 5% sorbitol treatment for 10 min at 37 °C. To achieve a tighter age window, the sorbitol treatment was repeated after the desired time interval.

### Plasmids and parasite lines generation

Genomic modification was achieved using the SLI system^60^ in NF54 (for the SMC1 line) or 3D7 parasites. All plasmids were produced using Gibson assembly^118^ with the primers described in Table S8. Sequencing was carried out to ensure absence of mutations in the cloned regions. All plasmid sequences are shown in File S1.

A target homology region corresponding to a fragment of the 3’ end of the coding sequence (for SMC1, HICs, EUCs and CICs) was amplified from 3D7 genomic DNA and cloned into the pSLI-GlmS vector using NotI and MluI restriction sites (in case of SMC1) or the pSLI-sandwich vector using NotI and AvrII restriction sites (in cases of HICs, EUCs and CICs). The HP1-tagged line was described in Birnbaum et al.^60^ and the CENH3-tagged line was described in Gockel et al.^61^.

For (mT)-DiQ-BioID experiments, the biotin ligase (BirAC^L^, BirA-N^L^ or mCherry-FRB-miniTurbo) was constitutively expressed from an episomal plasmid described elsewhere (BirA^57^ and miniTurbo^61^). The Lyn-FRBmCherry^epi^ plasmid^60^ was used for KS experiments in EUCs and CICs knock-in lines. For HIC8 co-localization with GRASP the pGRASP-mCherry-BSD^nmd3^ plasmid^60^ was used. For HP1 co-localization experiments while simultaneously carrying out KS of HICs, we used the HP1mCherry_Lyn-FRB plasmid, which was obtained by replacing the *Rab5* sequence in the Rab5mCherry^sf3a2^_Lyn-FRB-T2A-yDHODH^nmd3^ plasmid^64^ with the HP1-coding sequence (amplified from 3D7 genomic DNA) between XhoI and AvrII restriction sites (see sequence in File S1).

### Transfection of parasites

For transfection, late schizonts were isolated on a percoll gradient^119^, mixed with 50 μg of DNA (dissolved in 10 μl TE buffer and 90 μl of Amaxa transfection solution: 90 mM NaPO_4_, 5 mM KCl, 0.15 mM CaCl_2_, 50 mM HEPES pH 7.3) and electroporated using the Amaxa system (Nucleofector II AAD-1001N Amaxa Biosystems, Germany), following the U-033 program. After electroporation the parasites were mixed with 300 μl uninfected RBCs and 100 μl RPMI medium and incubated for 60-90 minutes at 37 °C under vigorous shaking, before being transferred to standard culturing conditions.

Transfected parasites for genomic integration were first selected for plasmid uptake with WR99210. Upon re-occurrence, these parasites were further selected for endogenous integration with G418 (C-terminal sandwich integration) or DSM1 (N-terminal sandwich integration). A correct integration of the tags was verified by PCR using gDNA from the modified lines (Figure S2) and using the primers described in Table S8. Parasite lines with BirAC^L^, BirA-N^L^ or mCherry-FRB-miniTurbo were further selected with blasticidine (BSD). Parasites with the LynFRB-mCherry^epi^ plasmid and those with the HP1mCherry_Lyn-FRB plasmid were maintained under DSM1 selection pressure.

### (mT)-DiQ-BioID experiments

DiQ-BioID experiments were carried out as previously described^57^. Asynchronous parasites were grown in a 200 ml culture until a parasitemia of 5-7%. Half of the culture was treated with rapalog for recruitment of the biotinylation enzyme to the target site and the other half was used as a control (background biotinylation). In experiments using BirA*, 250 nM of rapalog and 50 μM of biotin (Sigma-Aldrich), or only biotin in the control plates, were added to the cultures and incubated for 24 hours with a medium and supplements replacement at 12 hours. In experiments using miniTurbo (mT-DiQ-BioID), parasites were grown in biotin-free medium (USBiological Life Sciences) for two days or until the desired harvesting conditions were reached. Rapalog incubation was allowed for 30 minutes (in +rapalog cultures) before the addition of biotin to both (+rapalog and control) cultures, and biotinylation was allowed for another 30 minutes at 37 °C.

Cultures were harvested by centrifugation and washed twice with DPBS. RBCs were lysed using 0.03% saponin in DPBS on ice for 10 minutes. The released parasites were washed five times in DPBS, lysed in 2 ml of lysis buffer (50mM Tris-HCl pH 7.5, 500 mM NaCl, 1% TritonX-100 and 0.4% SDS) containing 1 mM dithiothreitol (DTT, Biomol), 2X protein inhibitor cocktail (Roche) and 1 mM phenylmethylsulfonyl fluoride (PMSF, Roche), and the lysate was frozen at -80 °C. For complete lysis, parasites were submitted to three freeze-thaw cycles and centrifuged at 16,000 g for 10 minutes. The soluble supernatant containing biotinylated proteins was split to obtain two technical replicates per experiment, diluted 2-fold in 50 mM Tris-HCl pH 7.5 containing 2X PIC and 1 mM PMSF and incubated with 25 μl of Streptavidin Sepharose (GE Healthcare) by rotating overnight at 4 °C. The beads were then washed twice in lysis buffer (without SDS), once in dH_2_O, twice in 50 mM Tris-HCl pH 7.5 and three times in 100 mM Triethylammonium bicarbonate buffer pH 8.5 (TEAB, Sigma-Aldrich). The beads were incubated in 50 μl of elution buffer (2 M Urea in 100 mM Tris pH 7.5 containing 10 mM DTT) at room temperature while shaking for 20 minutes, followed by treatment with 50 mM iodoacetamide (IAA) while shaking in the dark for 10 minutes. For on-bead digestion^120^, 0.25 μg of Trypsin/LysC (Promega) was added to each sample and incubated at room temperature for 2 hours. The eluted peptides were collected in the supernatant and the beads were rinsed with an extra 50 μl of elution buffer for 5 minutes at room temperature. The supernatant was again collected and combined with the previous elution. The total 100 μl of elution was added an additional 0.1 μg of Trypsin/LysC and peptides were further digested overnight while shaking at room temperature.

### ProtA-TurboID experiments

All steps for native nuclei collection were performed on ice and using ice-cold solutions. Asynchronous cultures at ∼5% parasitemia were washed with PBS, suspended in 0.05% saponin in PBS for RBC lysis and subsequently loaded on a two-layer sucrose gradient (top layer: 0.1 M; bottom layer: 0.25 M) in cell lysis buffer (CLB, 10 mM Tris pH 8.0, 3 mM MgCl_2_, 0.2% NP40) containing protease inhibitor cocktail (Roche). After 20 minutes centrifugation at 3100 g and 4 °C with acceleration and deceleration set to 1 (Eppendorf 5910 Ri, Rotor S-4x400), the native nuclei recovered from the bottom of the tube were washed with CLB and resuspended in CLB with 20% glycerol and snap-frozen in liquid nitrogen for storage at -80 °C. For ProtA-TurboID experiments the nuclei were treated following the protocol for unfixed cells^62^. Three replicates for control IgG and three replicates for each target antibody were set up in parallel. Each sample with native nuclei was permeabilized in digitonin buffer (0.04% digitonin, 20 mM Hepes pH 7.5, 150 mM NaCl, 0.5 mM spermidine, cOmplete protease inhibitor complex) supplemented with 2 μg of primary antibody: α-H3K27ac (Abcam), α-H3K4me3 (Diagenode), α-HP1^14^, α-GFP (Abcam) or α-IgG (Merck); and incubated at room temperature for 20 minutes while shaking. Then, the nuclei were washed twice with digitonin buffer and resuspended in digitonin buffer with 1.4 μg of recombinant protein A-TurboID fusion protein for 30 minutes rotating at 4 °C, followed by 5 minutes at room temperature. After two washes with digitonin buffer, samples were incubated in biotinylation buffer (digitonin buffer with additional 1 mM ATP, 5 μM biotin and 5 mM MgCl_2_) for 10 minutes shaking at 37 °C. After two washes with wash buffer (20 mM Hepes pH 7.5, 150 mM NaCl, 0.5 mM spermidine, cOmplete protease inhibitor complex), nuclei were resuspended in RIPA buffer and incubated on ice overnight. For a complete nuclear lysis, samples we sonicated (Bioruptor® Pico, Diagenode) for 18x 30” pulses and centrifuged at 11,000 g for 10 minutes. Biotinylated proteins were isolated from the supernatant with 25 μl of Streptavidin Sepharose slurry (GE Healthcare) for 2 hours in rotation at room temperature. Then the beads were washed five times with RIPA buffer and four times with PBS. Subsequently, the beads were resuspended in 50 ul elution buffer and followed the protocol for IAA treatment and on-beads digestion as described in the section above ((mT)-DiQ-BioID experiments).

### Mass Spectrometry (MS) sample preparation

The peptides were prepared as described elsewhere^64^. All samples were loaded on methanol-activated and equilibrated C18 membranes set up on stage-tips^121^. For label-free quantification (ProtA-TurboID experiments), peptides were acidified with 1% TFA before loading to the stage-tip. Peptides were desalted by washing with buffer A (0.1% formic acid), and in the case of (mT)-DiQ-BioID and GFP-pulldown experiments, peptides were dimethyl-labelled as previously described^122^. Technical replicates were processed under label-swap conditions, as follows: rapalog-treated samples carried a “heavy” label and control samples carried a “light” label in the forward experiments; whereas in the reverse experiments, rapalog-treated samples carried a “light” label whereas control samples carried a “heavy” label. In the case of GFP pulldowns, α-GFP and α-IgG control samples were processed in the same way. The “light” label consists of NaBH_3_CN (Merck) in formaldehyde solution (CH_2_O, Sigma-Aldrich). The “heavy” label contains NaBD_3_CN (Sigma-Aldrich) in formaldehyde-^13^C, d2 solution (^13^CD_2_O, Sigma-Aldrich). Each “light” and “heavy”-labelled samples were eluted with buffer B (80% acetonitrile, 0.1% formic acid) into a single tube, whereas label-free samples were eluted separately. The acetonitrile was evaporated using a 20 minute vacuum spin and the samples were reconstituted up to 12 μl with buffer A. Subsequently, 5 μl of the reconstituted sample was analyzed during a 60-minute gradient of increasing buffer B on an Easy-nLC 1000 (Thermo Fisher Scientific) with a 30 cm C18-reverse phase column coupled on-line to an Orbitrap Exploris 480 Mass Spectrometer (Thermo Fisher Scientific). The data was acquired in data-dependent top speed mode in a 3 sec cycle with dynamic exclusion set at 60 sec. The resolution for mass spectrometry was set at 120.000.

### Conditional inactivation (KS) assays

To assess the phenotype upon knock-sideways (KS)^60^ we used the pLyn-FRBmCherry-DHODH^nmd3^ plasmid^60^ for recruitment of the nuclear, sandwich-tagged proteins to the plasma membrane. For KS induction, cultures were split in two and both plates were treated under exact conditions except for the addition of rapalog (Takara, 250 nM final concentration) to one of them, while keeping the rapalog-free dish as control. The removal of the target protein from the nucleus (as compared to the control culture) was confirmed by live-cell fluorescence microscopy.

Flow cytometry was used to monitor the growth of parasites in knock-sideways experiments, by measuring the parasitemia for 5 consecutive days. For each flow cytometry sample, cultures were thoroughly resuspended and a 20 μl sample was diluted 1-in-5 with RPMI containing 0.5 mg/ml dihydroethidium (DHE) and 0.45 mg/ml Hoechst 33342 and incubated for 20 minutes in the dark at room temperature. Stained parasites were then added 400 μl RPMI containing 0.000325% glutaraldehyde. The parasitemia was measured by flow cytometry using an LSR-II cytometer by counting 100,000 events using the FACSDiva software (BD Biosciences).

For stage distribution assessments in CIC2- and CIC4-KS experiments, tightly synchronized parasites were submitted to KS and each parasite stage was manually counted in Giemsa smears at the time points indicated in Figure 4.

### Live cell imaging

Imaging of live parasites was performed as previously described^123^. Parasite nuclei were stained with 4.5 μg/ml Hoechst-33342 (Invitrogen) for 10 minutes at room temperature before imaging. When indicated, parasites were stained with 1 μM Tubulin Tracker™ Deep Red (Thermo Fisher) for 20 minutes at 37 °C. Images were acquired using a Zeiss AxioImager M1 or M2 fluorescence microscope equipped with different filters (Zeiss cubes 44, 49, 64 and 50), Zeiss Plan-apochromat 63x or 100x oil immersion lenses, and a C4742-80 Hamamatsu Orca Digital camera. Images were collected using the AxioVision software (version 4.7), and the brightness and intensity were adjusted and the channels were merged in Corel PhotoPaint (version 2023). 3D stacks of live parasites were obtained using and the Olympus 60x/1.50 UPlanApo oil immersion lens of an Olympus FluoView FV4000 confocal laser scanning microscope and the FluoView software (version 2.6). Images were processed using Imaris 7.7.2 (Bitplane) to adjust for contrast and brightness.

### Gametocytogenesis assay

HIC7 parasites carrying the HP1-mCherry_Lyn-FRB plasmid were synchronized to a 4-hour window. At 20-24 hours post invasion (hpi), HIC7-KS was induced by splitting the culture in two and adding 250 nM rapalog (Takara) to one of them. Two days after, at 20-24 hpi parasitemia was measured by flow cytometry, and heparin (Heparin-Natrium-5000-ratiopharm®; 39,5 I.U. per 1 ml medium) was added to prevent invasion of merozoites. On day 6 (see schematics on Figure 2G), parasites were stained for 25 minutes with 4,5 μg/ml Hoechst-33342 (Invitrogen) and 1 μM Tubulin Tracker™ Deep Red (Invitrogen) for fluorescence microscopy images acquisition. Red blood cells were manually counted from DIC images and gametocytes were counted in the corresponding tubulin-stained layer. At least 2000 red blood cells were counted for each condition. Conversion rate was calculated according to following formula:

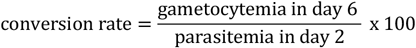

### HP1 genome-wide occupancy profiling

To assess the rearrangements of HP1 occupancy in HIC7-KS compared to control parasites, we performed CUT&Tag^61,124^ in isolated nuclei. Following the protocol by Gockel et al.^61^, parasites were crosslinked with 0.1% formaldehyde (Sigma) for 2 min at 37 °C while shaking. Crosslinking was stopped by addition of glycine at 0.125 M final concentration. Parasite nuclei were then isolated on a sucrose gradient as described above (see ProtA-TurboID section). Nuclei were counted in an automatic hemocytometer (BioRad, TC10 Automated Cell Counter). Approximately 1 million nuclei were used per sample. Nuclei were resuspended in CUT&Tag wash buffer (20 mM HEPES pH 7.5, 150 mM NaCl, 0.5 mM Spermidine, 1x EDTA-free protease inhibitor) containing 0.1% Triton X100 and permeabilized for 10 minutes on ice. Nuclei were pelleted by centrifugation (3500 g, 10 minutes, 4 °C) and resuspended in CUT&Tag wash buffer. Concanavalin A beads (Bangs Laboratories) were activated by resuspending into 10 volumes of Bead Binding Buffer (20 mM HEPES pH 7.5, 10 mM KCl, 1 mM CaCl_2_, 1 mM MnCl_2_), washed once on a magnetic rack and resuspended in the starting volume of bead slurry. Purified nuclei were bound to 10 μl beads per reaction by incubating for 10 minutes rotating at room temperature. The supernatant was removed and 50 μl antibody buffer (CUT&Tag wash buffer; 2 mM EDTA, 0.1% BSA) with primary antibody (0.25 μl polyclonal rabbit αHP1^14^) was added and incubated nutating over night at 4 °C. Unbound primary antibody was removed by washing once with 100 μl CUT&Tag wash buffer and samples were then incubated with secondary antibody (1.2 µg guinea pig anti-rabbit antibody, Antibodies-Online) in 100 μl CUT&Tag wash buffer, 1:100 dilution for 1 hour at room temperature, nutating. The nuclei were washed on a magnetic stand twice with 100 μl CUT&Tag wash buffer and once with 100 μl CUT&Tag 300 Wash Buffer (20 mM HEPES pH 7.5, 300 mM NaCl, 0.5 mM Spermidine, 1x EDTA-free protease inhibitor). 2.5 μl of proteinA/G-Tn5 fusion protein (CUTANA™ pAG-Tn5 for CUT&Tag, Epicypher) was added in 50 μl CUT&Tag 300 wash buffer and incubated nutating for 1 hour. Unbound proteinA/G-Tn5 fusion protein was removed by washing thrice with 100 μl CUT&Tag 300 wash buffer. The nuclei were resuspended in 200 μl tagmentation buffer (CUT&Tag 300 wash buffer, 10 mM MgCl_2_) and incubated in a PCR thermocycler (BioRad, T100) at 37 °C for 1 hour. Tagmentation was stopped and nuclei lysis was facilitated by addition of 10 μl of 0.5M EDTA pH 8, 3 μl of 10% SDS and 1 μl of 50 mg/ml proteinase K. Samples were briefly vortexed and then incubated at 55 °C for 1 hour. DNA fragments were extracted utilizing the DNA Clean & Concentrator - 5 kit (Zymogen, D4014). DNA was eluted from the column with 26 μl of prewarmed elution buffer and DNA concentrations were assessed with Qubit dsDNA High Sensitivity Assay kit (Invitrogen).

For library preparation, up to 50 ng of DNA were amplified using i5 and i7 primers^125^ and using Kapa HiFi polymerase (Roche) with the following program: 58 °C for 5 minutes, 62 °C for 5 minutes (gap filling), 98 °C for 2 minutes, 14 cycles of 98 °C for 20 seconds and 62 °C for 10 seconds, 62 °C for 1 min minute and hold at 4 °C. Post-PCR DNA cleanup was performed by adding 50 μl (1x volume) of AMPure XP bead slurry (Beckman Coulter, A63882) and incubating for 10 minutes at room temperature, washing twice with 80% EtOH on a magnetic rack, and eluting in 16.5 μl of 10 mM Tris-HCl pH 8 for 5 min at room temperature. DNA concentrations of libraries were assessed with Qubit dsDNA High Sensitivity Assay kit (Invitrogen) and library fragment size distribution was accessed by microfluidic gel electrophoresis (Agilent 2100 Bioanalyzer) with the corresponding High Sensitivity DNA Kit (Agilent). Sequencing was performed using an Illumina NextSeq 2000 instrument; 59bp paired-end reads were generated.

### Bulk RNA-seq experimental set-up and library preparation

Knock-sideways was induced in synchronized parasites at ring stage by rapalog addition, while a control culture was handled in parallel without rapalog. Parasites were either harvested at schizont stage on the same cycle, or at ring stage on the next cycle. Cultures harvested at rings were additionally synchronized prior to harvesting at 6-12 hpi (hours post invasion). Total RNA was isolated from TRIzol suspended infected RBC pellet using the Directzol RNA Miniprep kit (Zymo Research) and passing the samples through 2x rounds of DNaseI treatment. The isolated total RNA was quantified using the Qubit™ RNA broad-range assay kit (Invitrogen). The integrity of total RNA (RIN Score >8.0) was verified using the Bioanalyzer RNA 6000 Nano kit (Agilent). Subsequently, 250ng and 500ng of total RNA was used for schizont and ring stage samples, respectively, as input to prepare mRNA library with the Kapa mRNA HyperPrep Kit (Roche) and using 12 and 13 PCR cycles to generate the amplified cDNA library for schizont and ring stage samples, respectively. The cDNA libraries concentration and size distribution were assessed and sequencing was performed as above for DNA libraries.

### GFP-pulldowns

Nuclear extracts were generated as on^126^. In short, isolated nuclei from GFP-tagged lines were incubated in High Salt Buffer (50 mM HEPES pH 7.5, 20% glycerol, 420 mM NaCl, 1.5 mM MgCl_2_, 1 mM DTT, 0.4% NP40, 400 units/ml DNaseI and protease inhibitor) after douncing to break apart possible nuclei clumps. The extracts were slightly diluted to reduce the NaCl concentration to 300 mM, and 0.2 mM EDTA and 50 μg/ml of ethidium bromide were added. The targeted protein complexes were captured in GFP-Trap beads (Chromotek, gta-20) and control pulldowns were done with the same extract in blocked agarose beads (Chromotek, bab-20). False positive proteins were identified from the negative control pulldowns performed with wild-type parasites in Hoeijmakers et al.^45^.

### IFA scoring of presence of antigens

CIC2- and CIC4-tagged parasite lines were synchronized twice using 5% sorbitol 6 hours apart, the culture was split and KS was induced in one dish by adding 250 mM rapalog. Parasites were grown with media change and rapalog replenishment for another 42 hours (one cycle) when they were harvested (rings of the next cycle, confirmed by Giemsa smear). Cells were fixed in 100% acetone as dried film on 10-well slides and IFAs carried out as described^127^. Anti-sera were used in two combinations that each detected REX1 and SBP1: rabbit anti-REX1 (1/1250) with mouse anti-SBP1C (1/625) and mouse anti-REX1 (1/625) and rabbit anti-SBP1C (1/1250)^128^. Secondary antibodies were donkey anti-rabbit conjugated with Alexa Fluor-488 and goat anti-mouse conjugated with Alexa Fluor-594 (Thermo Ficher) diluted 1:2000. For each condition, wells with both combinations of antibodies were prepared. Wells with the swamped primary antibodies were individually scored and the average of both wells was used as one biological replicate. For scoring, all cells with a Hoechst signal and at least 4 dots indicative of Maurer’s cleft staining in either channel were included to determine the total number of cells in a section. Of these sections, the percentage of cells without one of the two signals were then determined. At least 75 cells were scored per well and condition. Scoring was done blinded to the condition of the samples analyzed.

### Ultrastructural expansion microscopy (U-ExM)

Sample preparation for U-ExM was performed as previously described^129^ with minor changes. The harvested parasites were fixed with 4% formaldehyde/0.0075% glutaraldehyde for 15 minutes at 37 °C. Glass coverslips (10 mm diameter) were coated with 0.5 mg/ml Concanavalin A (ConA) in water for 20 minutes at room temperature. After removal of excess ConA, the parasites were sedimented onto the coverslips (1% haematocrit) and incubated for 10 minutes at room temperature. The unbound RBCs and parasites were carefully washed off with 1x PBS until only a faint red-golden layer was left. For protein anchoring, the coated cells were incubated with pre-warmed anchoring solution (1.4% formaldehyde, 2% acrylamide in 1x PBS) overnight at 37 °C. For gelation of the anchored parasites on the next day, 90 μl monomer solution (19% sodium acrylate, 10% acrylamide, 0.1% N,N’-methylenebisacylamide in 1x PBS) were mixed with 5 μl of 10% TEMED and 5 μl 10% of APS, placed on an ice-cold parafilm (30 μl per gel) and immediately a coverslip was placed on top of the gel drop. After gelation, the gels were separated from the coverslips by incubation for 15 minutes in denaturation buffer (200 mM SDS, 200 mM NaCl, 50 mM Tris, pH 9) at 37 °C with gently shaking. The gels were then transferred into 1.5 ml Eppendorf tubes and the embedded proteins were denatured in 1 ml denaturation buffer for 90 minutes at 95 °C. This was followed by a first expansion round of the gels with water in a 10 ml dish for 30 minutes on a platform shaker. Next, the expanded gels were shrunken again by removing the water and adding 1x PBS to the dish and incubate for 15 minutes shaking. The shrunken gels were cut into three parts, each of which was transferred into an 1.5 ml Eppendorf tube for blocking in 3% bovine serum albumin/1x PBS for 30 minutes with constant shaking. Subsequently, the gels were incubated with rabbit polyclonal anti-mCherry (1:500; Abcam) overnight at room temperature and after washing the next day, with goat anti-rabbit Alexa Fluor^®^ 633 antibody (1:500; Thermo Fisher) for 2 hours at 37 °C. Following washing steps, the gels were incubated with 10 μg/ml DyLight488-NHS-ester (Thermo Fisher) in 1x PBS (for protein staining) and 2 μg/ml Hoechst 33342 (for DNA staining). Following staining, the gels were expanded in water and, for gels with membrane staining, they were stained with Bodipy TR Ceramide (1:500, Invitrogen) for 1 hour in water and placed on poly-L-lysine pre-treated ibidi slides for imaging. The expanded parasites were imaged on an Olympus FluoView FV4000 confocal laser scanning microscope using the Olympus 60x/1.50 UPlanApo oil immersion lens and FluoView software (version 2.6). The samples were quickly scanned first using the green laser (NHS signal) and the UV laser (Hoechst signal) only and then, in an area with properly stained parasites, 3D stacks with the specific fluorophore signals were captured using respective lasers. The image stacks were processed using Imaris 7.7.2 software, brightness and intensity of images were adjusted, and the channels were merged in Corel Photo Paint (version X6).

### Mass spectrometry data analysis

Raw files were analyzed using MaxQuant^130^ (version 1.6.6.0 for HP1, H3K27ac, H3K4me3, EUC3, CENH3 and SMC1; or version 2.1.4.0 for HIC7, EUC4, EUC6; version 2.4.2.0 for EUC2), using default parameters except for the following. Multiplicity was set at 2, with the light labels (DimethLys0 and DimethNter0) and heavy labels (DimethLys8 and DimethNter8) selected. Deamidation (NQ) was added as a variable modification together with oxidation (M) and acetyl (N-term). Match-between-runs and (except for ProtA-TurboID experiments) re-quantify options were enabled with default parameters and iBAQ values were calculated. Mass spectra were compared to peptide masses from the *Plasmodium falciparum* 3D7 annotated proteome (PlasmoDB release 59). The output ProteinGroup file was submitted to Perseus software package (version 1.4.0.20; or version 1.5.0.15 for ProtA-TurboID experiments)^131^. Proteins were filtered against ‘peptides only identified by site’, ‘reverse’ and ‘potential contaminant’ positive values. In (mT)-DiQ-BioID and GFP-pulldown datasets, H/L normalized values were log2 transformed and reverse experiments were (-x) transformed to obtain rapalog/control ratios. In ProtA-TurboID datasets, intensity values were transformed by log2 and imputation was applied to replace missing values using Perseus default settings (normal distribution, width of 0.3, down shift of 1.8). One-sample (dimethyl-labelled experiments) or two-sample (for label-free ProtA-Turbo experiments) t-tests were run to calculate the mean fold change (t-test difference) and -log10 t-test p-values. Twosided Benjamini-Hochberg test was used to calculate a corrected FDR (q-value).

To obtain high confidence datasets for each chromatin state, we included proteins that meet strict cut-offs for three parameters combined: fold change (log2 ratio) values, t-test p-values and unique peptides detected for each protein in the MS run. For all experiments, minimum thresholds were drawn at (-log10) p-values of 1.3 and 4 unique peptides. Fold change cut-offs were drawn at a value of 1 for HP1 and CENH3 (mT)-DiQ-BioIDs, at 1.5 for HP1 and H3K27ac ProtA-TurboIDs, at 0.8 for H3K4me3 ProtA-TurboID, and at 2 for SMC1 ProtA-TurboID.

### CUT&Tag data analysis

Raw sequencing data was subset to 2.2 M reads per sample using seqtk (v1.2) to minimize differences due to sequencing depth. Paired sequencing reads were mapped (pair-end) against the reference genome PlasmoDB v26 3D7 using bowtie2 (v2.5.2)^132^. Reads were filtered by mapping quality >= 30 and mitrochondrial as well as apicoplast reads were removed with samtools (v1.18)^133^. BigWig files normalized to read per million per kilobase (RPMK) were created using deeptools (v3.5.4)^134^. BedGraph files were generated for visualization purposes on the UCSC genome browser^135^ with bedtools (v2.31.0)^136^ normalized to library size. Log2 ratio tracks were generated by running the multiBigwigSummary command from deeptools (v3.5.4)^134^ using 2000 bp bins on both +rapalog and control bigwig files. A pseudocount of 1 was added to all values to prevent divisions by 0, and log2 values were calculated in bash and appended to a new bedGraph file for visualization on UCSC genome browser^135^. Detailed & customizable scripts can be found on our github page (https://github.com/bartfai-lab/DiBioCUTnTag-Analysis) or Zenodo (10.5281/zenodo.15638929).

### RNA-seq data analysis

The fastq files from sequencing were evaluated for quality using the FASTQC package (v0.11.9). Trim-galore package (v0.6.7) was used to trim off adapter sequence and bases with base calling quality score <30. The trimmed fastq were aligned against the *P. falciparum* genome reference (Pf strain 3D7 v68 fasta; edited to remove mitochondrial and apicoplast genome) using the splice-aware STAR aligner (v2.7.10a)^137^. The aligned bams were filtered for uniqueness and high mapping quality (samtools view -q 255) using the samtools package (v1.15.1). The resulting bam file was processed for removing duplicate alignments using the MarkDuplicate function in Picard package (v2.18.23-SNAPSHOT). Finally raw gene-counts were generated using featureCounts (v2.0.1 from subread package)^138^ in a strandedness and paired-end aware manner (feature-Counts -s 2 -p -B). Feature assignment of > 90% was reported for all samples.

Differential gene expression analysis was performed in RStudio (R version 4.3.3, “Angel Food Cake”) using the Deseq2 package (v1.40.2)^139^. The ring and schizont stage analysis was done independently. The Deseq2 object ‘dds’ was pre-filtered to retain only genes with >10 sum raw-counts in at least 3 samples for ring-stage and 2 samples for schizont-stage (the lowest sample size per stage). Post library size estimation, estimation of gene dispersion and negative binomial general linear model (GLM) fit, normalized gene counts per sample were obtained. Variance stabilizing transformation was applied to gene counts (vst(dds, blind=FALSE)) for visualization purposes. Differential gene expression analysis across control and rapalog treatment conditions was done with the results() function in Deseq2. Genes were filtered as differentially expressed or stage-specific markers if their expression baseMean>=30, absolute(log2Fold-Change)>=1 and p.adjusted<=0.01 compared to reference. Volcano plot for differential expression visualization was generated using the ggplot2 function (v3.5.1)^140^. Scaled gene expression heatmaps were generated using the pheatmap (v1.0.12) package. Supplementary information regarding i) expression of gametocytogenesis marker genes expression in the malaria cell atlas (sexual vs asexual), ii) AP2-G genomic target enrichment across early committed and sexual stages and iii) HP1 enrichment across asexual stage parasites was manually curated from relevant publications for comparison without marker list.

## REFERENCES

1. Lorentz, A., Ostermann, K., Fleck, O., and Schmidt, H. (1994). Switching gene swi6, involved in repression of silent mating-type loci in fission yeast, encodes a homologue of chromatin-associated proteins from Drosophila and mammals. Gene 143, 139–143. 10.1016/0378-1119(94)90619-x.

2. Moris, N., Pina, C., and Arias, A.M. (2016). Transition states and cell fate decisions in epigenetic landscapes. Nat. Rev. Genet. 2016 17, 693–703. 10.1038/nrg.2016.98.

3. Mahmood, T., He, S., Abdullah, M., Sajjad, M., Jia, Y., Ahmar, S., Fu, G., Chen, B., and Du, X. (2024). Epigenetic insight into floral transition and seed development in plants. Plant Sci. 339. 10.1016/j.plantsci.2023.111926.

4. Greenberg, M.V.C., and Bourc’his, D. (2019). The diverse roles of DNA methylation in mammalian development and disease. Nat. Rev. Mol. Cell Biol. 20, 590–607. 10.1038/s41580-019-0159-6.

5. Janssen, S.M., and Lorincz, M.C. (2022). Interplay between chromatin marks in development and disease. Nat. Rev. Genet. 23, 137–153. 10.1038/s41576-021-00416-X.

6. Klemm, S.L., Shipony, Z., and Greenleaf, W.J. (2019). Chromatin accessibility and the regulatory epigenome. Nat. Rev. Genet. 20, 207–220. 10.1038/s41576-018-0089-8.

7. Gourisankar, S., Krokhotin, A., Wenderski, W., and Crabtree, G.R. (2024). Context-specific functions of chromatin remodellers in development and disease. Nat. Rev. Genet. 25, 340–361. 10.1038/S41576-023-00666-X,.

8. Li, X., and Fu, X.D. (2019). Chromatin-associated RNAs as facilitators of functional genomic interactions. Nat. Rev. Genet. 20, 503–519. 10.1038/s41576-019-0135-1.

9. Bell, O., Burton, A., Dean, C., Gasser, S.M., and Torres-Padilla, M.E. (2023). Heterochromatin definition and function. Nat. Rev. Mol. Cell Biol. 24, 691–694. 10.1038/s41580-023-00599-7.

10. Klughammer, J., Romanovskaia, D., Nemc, A., Posautz, A., Seid, C.A., Schuster, L.C., Keinath, M.C., Lugo Ramos, J.S., Kosack, L., Evankow, A., et al. (2023). Comparative analysis of genome-scale, base-resolution DNA methylation profiles across 580 animal species. Nat. Commun. 14, 1–23. 10.1038/s41467-022-34828-y.

11. Niederhuth, C.E., Bewick, A.J., Ji, L., Alabady, M.S., Kim, K. Do, Li, Q., Rohr, N.A., Rambani, A., Burke, J.M., Udall, J.A., et al. (2016). Widespread natural variation of DNA methylation within angiosperms. Genome Biol. 17, 1–19. 10.1186/S13059-016-1059-0.

12. Bunnik, E.M., Venkat, A., Shao, J., McGovern, K.E., Batugedara, G., Worth, D., Prudhomme, J., Lapp, S.A., Andolina, C., Ross, L.S., et al. (2019). Comparative 3D genome organization in apicomplexan parasites. Proc. Natl. Acad. Sci. U. S. A. 116, 3183–3192. 10.1073/pnas.1810815116.

13. Blombach, F., and Werner, F. (2024). Chromatin and gene regulation in archaea. Mol. Microbiol. 123. 10.1111/mmi.15302.

14. Brancucci, N.M.B., Bertschi, N.L., Zhu, L., Niederwieser, I., Chin, W.H., Wampfler, R., Freymond, C., Rottmann, M., Felger, I., Bozdech, Z., et al. (2014). Heterochromatin protein 1 secures survival and transmission of malaria parasites. Cell Host Microbe 16, 165–176. 10.1016/j.chom.2014.07.004.

15. Lopez-Rubio, J.J., Mancio-Silva, L., and Scherf, A. (2009). Genomewide Analysis of Heterochromatin Associates Clonally Variant Gene Regulation with Perinuclear Repressive Centers in Malaria Parasites. Cell Host Microbe 5, 179–190. 10.1016/j.chom.2008.12.012.

16. Voss, T.S., and Brancucci, N.M. (2024). Regulation of sexual commitment in malaria parasites — a complex affair. Curr. Opin. Microbiol. 79, 102469. 10.1016/j.mib.2024.102469.

17. Reyser, T., Paloque, L., Augereau, J.M., Di Stefano, L., and Benoit-Vical, F. (2024). Epigenetic regulation as a therapeutic target in the malaria parasite Plasmodium falciparum. Malar. J. 23, 44. 10.1186/s12936-024-04855-9.

18. World Health Organization (2024). World malaria report 2024. https://www.who.int/teams/global-malaria-programme/reports/world-malaria-report-2024.

19. Bozdech, Z., Llinás, M., Pulliam, B.L., Wong, E.D., Zhu, J., and DeRisi, J.L. (2003). The Transcriptome of the Intraerythrocytic Developmental Cycle of Plasmodium falciparum. PLOS Biol. 1, e5. 10.1371/journal.pbio.0000005.

20. Rovira-Graells, N., Gupta, A.P., Planet, E., Crowley, V.M., Mok, S., De Pouplana, L.R., Preiser, P.R., Bozdech, Z., and Cortés, A. (2012). Transcriptional variation in the malaria parasite Plasmodium falciparum. Genome Res. 22, 925–938. 10.1101/GR.129692.111.

21. Modrzynska, K., Pfander, C., Chappell, L., Yu, L., Suarez, C., Dundas, K., Gomes, A.R., Goulding, D., Rayner, J.C., Choudhary, J., et al. (2017). A Knockout Screen of ApiAP2 Genes Reveals Networks of Interacting Transcriptional Regulators Controlling the Plasmodium Life Cycle. Cell Host Microbe 21, 11–22. 10.1016/j.chom.2016.12.003.

22. Gardner, M.J., Hall, N., Fung, E., White, O., Berriman, M., Hyman, R.W., Carlton, J.M., Pain, A., Nelson, K.E., Bowman, S., et al. (2002). Genome sequence of the human malaria parasite Plasmodium falciparum. Nature 419, 498–511. 10.1038/nature01097.

23. Hoeijmakers, W.A.M., Stunnenberg, H.G., and Bártfai, R. (2012). Placing the Plasmodium falciparum epigenome on the map. Trends Parasitol. 28, 486–495. 10.1016/J.PT.2012.08.006.

24. Trelle, M.B., Salcedo-Amaya, A.M., Cohen, A.M., Stunnenberg, H.G., and Jensen, O.N. (2009). Global histone analysis by mass spectrometry reveals a high content of acetylated lysine residues in the malaria parasite Plasmodium falciparum. J. Proteome Res. 8, 3439–3450. 10.1021/pr9000898.

25. Coetzee, N., Sidoli, S., Van Biljon, R., Painter, H., Llinás, M., Garcia, B.A., and Birkholtz, L.M. (2017). Quantitative chromatin proteomics reveals a dynamic histone post-translational modification landscape that defines asexual and sexual Plasmodium falciparum parasites. Sci. Rep. 7, 1–12. 10.1038/s41598-017-00687-7.

26. Grüning, H. von, Coradin, M., Mendoza, M.R., Reader, J., Sidoli, S., Garcia, B.A., and Birkholtz, L.-M. (2021). A dynamic and combinatorial histone code drives malaria parasite asexual and sexual development. bioRxiv, 2021.07.19.452879. 10.1101/2021.07.19.452879.

27. Salcedo-Amaya, A.M., Van Driel, M.A., Alako, B.T., Trelle, M.B., Van Den Elzen, A.M.G., Cohen, A.M., Janssen-Megens, E.M., Van De Vegte-Bolmer, M., Selzer, R.R., Iniguez, A.L., et al. (2009). Dynamic histone H3 epigenome marking during the intraerythrocytic cycle of Plasmodium falciparum. Proc. Natl. Acad. Sci. U. S. A. 106, 9655–9660. 10.1073/pnas.0902515106.

28. Freitas, L.H., Hernandez-Rivas, R., Ralph, S.A., Montiel-Condado, D., Ruvalcaba-Salazar, O.K., Rojas-Meza, A.P., Mâncio-Silva, L., Leal-Silvestre, R.J., Gontijo, A.M., Shorte, S., et al. (2005). Telomeric heterochromatin propagation and histone acetylation control mutually exclusive expression of antigenic variation genes in malaria parasites. Cell 121, 25–36. 10.1016/j.cell.2005.01.037.

29. Duraisingh, M.T., Voss, T.S., Marty, A.J., Duffy, M.F., Good, R.T., Thompson, J.K., Freitas, L.H., Scherf, A., Crabb, B.S., and Cowman, A.F. (2005). Heterochromatin silencing and locus repositioning linked to regulation of virulence genes in Plasmodium falciparum. Cell 121, 13–24. 10.1016/j.cell.2005.01.036.

30. Salcedo-Amaya, A.M., Hoeijmakers, W.A.M., Bártfai, R., and Stunnenberg, H.G. (2010). Malaria: could its unusual epigenome be the weak spot? Int. J. Biochem. Cell Biol. 42, 781–784. 10.1016/j.biocel.2010.03.010.

31. Flueck, C., Bartfai, R., Volz, J., Niederwieser, I., Salcedo-Amaya, A.M., Alako, B.T.F., Ehlgen, F., Ralph, S.A., Cowman, A.F., Bozdech, Z., et al. (2009). Plasmodium falciparum heterochromatin protein 1 marks genomic loci linked to phenotypic variation of exported virulence factors. PLoS Pathog. 5. 10.1371/journal.ppat.1000569.

32. Fraschka, S.A., Filarsky, M., Hoo, R., Niederwieser, I., Yam, X.Y., Brancucci, N.M.B., Mohring, F., Mushunje, A.T., Huang, X., Christensen, P.R., et al. (2018). Comparative Heterochromatin Profiling Reveals Conserved and Unique Epigenome Signatures Linked to Adaptation and Development of Malaria Parasites. Cell Host Microbe 23, 407-420.e8. 10.1016/j.chom.2018.01.008.

33. Connacher, J., Josling, G.A., Orchard, L.M., Reader, J., Llinás, M., and Birkholtz, L.M. (2021). H3K36 methylation reprograms gene expression to drive early gametocyte development in Plasmodium falciparum. Epigenetics and Chromatin 14, 1–15. 10.1186/s13072-021-00393-9.

34. Voss, T.S., Bozdech, Z., and Bártfai, R. (2014). Epigenetic memory takes center stage in the survival strategy of malaria parasites. Curr. Opin. Microbiol. 20, 88–95. 10.1016/j.mib.2014.05.007.

35. Filarsky, M., Fraschka, S.A., Niederwieser, I., Brancucci, N.M.B., Carrington, E., Carrió, E., Moes, S., Jenoe, P., Bártfai, R., and Voss, T.S. (2018). GDV1 induces sexual commitment of malaria parasites by antagonizing HP1-dependent gene silencing. Science 359, 1259–1263. 10.1126/science.aan6042.

36. Toenhake, C.G., Voorberg-van der Wel, A., Wu, H., Kanyal, A., Nieuwenhuis, I.G., van der Werff, N.M., Hofman, S.O., Zeeman, A.M., Kocken, C.H.M., and Bártfai, R. (2023). Epigenetically regulated RNA-binding proteins signify malaria hypnozoite dormancy. Cell Rep. 42. 10.1016/j.celrep.2023.112727.

37. Michel-Todó, L., Bancells, C., Casas-Vila, N., Rovira-Graells, N., Hernández-Ferrer, C., González, J.R., and Cortés, A. (2023). Patterns of Heterochromatin Transitions Linked to Changes in the Expression of Plasmodium falciparum Clonally Variant Genes. Microbiol. Spectr. 11. 10.1128/spectrum.03049-22.

38. Bártfai, R., Hoeijmakers, W.A.M., Salcedo-Amaya, A.M., Smits, A.H., Janssen-Megens, E., Kaan, A., Treeck, M., Gilberger, T.W., Franc oijs, K.J., and Stunnenberg, H.G. (2010). H2A.Z demarcates intergenic regions of the plasmodium falciparum epigenome that are dynamically marked by H3K9ac and H3K4me3. PLoS Pathog. 6. 10.1371/journal.ppat.1001223.

39. Chaal, B.K., Gupta, A.P., Wastuwidyaningtyas, B.D., Luah, Y.H., and Bozdech, Z. (2010). Histone deacetylases play a major role in the transcriptional regulation of the Plasmodium falciparum life cycle. PLoS Pathog. 6. 10.1371/journal.ppat.1000737.

40. Hoeijmakers, W.A.M., Salcedo-Amaya, A.M., Smits, A.H., Françoijs, K.J., Treeck, M., Gilberger, T.W., Stunnenberg, H.G., and Bártfai, R. (2013). H2A.Z/H2B.Z double-variant nucleosomes inhabit the AT-rich promoter regions of the Plasmodium falciparum genome. Mol. Microbiol. 87, 1061. 10.1111/mmi.12151.

41. Petter, M., Selvarajah, S.A., Lee, C.C., Chin, W.H., Gupta, A.P., Bozdech, Z., Brown, G. V., and Duffy, M.F. (2013). H2A.Z and H2B.Z double-variant nucleosomes define intergenic regions and dynamically occupy var gene promoters in the malaria parasite Plasmodium falciparum. Mol. Microbiol. 87, 1167–1182. 10.1111/mmi.12154.

42. Ngwa, C.J., Kiesow, M.J., Papst, O., Orchard, L.M., Filarsky, M., Rosinski, A.N., Voss, T.S., Llinás, M., and Pradel, G. (2017). Transcriptional Profiling Defines Histone Acetylation as a Regulator of Gene Expression during Human-to-Mosquito Transmission of the Malaria Parasite Plasmodium falciparum. Front. Cell. Infect. Microbiol. 7. 10.3389/fcimb.2017.00320.

43. Azizan, S., Selvarajah, S.A., Tang, J., Jeninga, M.D., Schulz, D., Pareek, K., Herr, T., Day, K.P., De Koning-Ward, T.F., Petter, M., et al. (2023). The P. falciparum alternative histones Pf H2A.Z and Pf H2B.Z are dynamically acetylated and antagonized by PfSir2 histone deacetylases at heterochromatin boundaries. MBio 14. 10.1128/mbio.02014-23.

44. Tang, J., Chisholm, S.A., Yeoh, L.M., Gilson, P.R., Papenfuss, A.T., Day, K.P., Petter, M., and Duffy, M.F. (2020). Histone modifications associated with gene expression and genome accessibility are dynamically enriched at Plasmodium falciparum regulatory sequences. Epigenetics Chromatin 13. 10.1186/S13072-020-00365-5.

45. Hoeijmakers, W.A.M., Miao, J., Schmidt, S., Toenhake, C.G., Shrestha, S., Venhuizen, J., Henderson, R., Birnbaum, J., Ghidelli-Disse, S., Drewes, G., et al. (2019). Epigenetic reader complexes of the human malaria parasite, Plasmodium falciparum. Nucleic Acids Res. 47, 11574–11588. 10.1093/nar/gkz1044.

46. Josling, G.A., Petter, M., Oehring, S.C., Gupta, A.P., Dietz, O., Wilson, D.W., Schubert, T., Längst, G., Gilson, P.R., Crabb, B.S., et al. (2015). A Plasmodium Falciparum Bromodomain Protein Regulates Invasion Gene Expression. Cell Host Microbe 17, 741–751. 10.1016/j.chom.2015.05.009.

47. Balaji, S., Madan Babu, M., Iyer, L.M., and Aravind, L. (2005). Discovery of the principal specific transcription factors of Apicomplexa and their implication for the evolution of the AP2-integrase DNA binding domains. Nucleic Acids Res. 33, 3994–4006. 10.1093/nar/gki709.

48. Kelley, J.M., McRobert, L., and Baker, D.A. (2006). Evidence on the chromosomal location of centromeric DNA in Plasmodium falciparum from etoposide-mediated topoisomerase-II cleavage. Proc. Natl. Acad. Sci. U. S. A. 103, 6706–6711. 10.1073/pnas.0510363103.

49. Hoeijmakers, W.A.M., Flueck, C., Françoijs, K.J., Smits, A.H., Wetzel, J., Volz, J.C., Cowman, A.F., Voss, T., Stunnenberg, H.G., and Bártfai, R. (2012). Plasmodium falciparum centromeres display a unique epigenetic makeup and cluster prior to and during schizogony. Cell. Microbiol. 14, 1391–1401. 10.1111/J.1462-5822.2012.01803.X.

50. Voorberg-van der Wel, A., Zeeman, A.M., van Amsterdam, S.M., van den Berg, A., Klooster, E.J., Iwanaga, S., Janse, C.J., van Gemert, G.J., Sauerwein, R., Beenhakker, N., et al. (2013). Transgenic Fluorescent Plasmodium cynomolgi Liver Stages Enable Live Imaging and Purification of Malaria Hypnozoite-Forms. PLoS One 8, e54888. 10.1371/journal.pone.0054888.

51. Iwanaga, S., Kato, T., Kaneko, I., and Yuda, M. (2012). Centromere Plasmid: A New Genetic Tool for the Study of Plasmodium falciparum. PLoS One 7, e33326. 10.1371/journal.pone.0033326.

52. Iwanaga, S., Khan, S.M., Kaneko, I., Christodoulou, Z., Newbold, C., Yuda, M., Janse, C.J., and Waters, A.P. (2010). Functional Identification of the Plasmodium Centromere and Generation of a Plasmodium Artificial Chromosome. Cell Host Microbe 7, 245–255. 10.1016/j.chom.2010.02.010.

53. Batugedara, G., Lu, X.M., Saraf, A., Sardiu, M.E., Cort, A., Abel, S., Prudhomme, J., Washburn, M.P., Florens, L., Bunnik, E.M., et al. (2020). The chromatin bound proteome of the human malaria parasite. Microb. genomics 6. 10.1099/MGEN.0.000327.

54. Pandey, R., Abel, S., Boucher, M., Wall, R.J., Zeeshan, M., Rea, E., Freville, A., Lu, X.M., Brady, D., Daniel, E., et al. (2020). Plasmodium Condensin Core Subunits SMC2/SMC4 Mediate Atypical Mitosis and Are Essential for Parasite Proliferation and Transmission. Cell Rep. 30, 1883-1897.e6. 10.1016/j.celrep.2020.01.033.

55. Uhlmann, F. (2016). SMC complexes: From DNA to chromosomes. Nat. Rev. Mol. Cell Biol. 17, 399–412. 10.1038/NRM.2016.30;SUBJMETA.

56. Rosa, C., Singh, P., Chen, P., Sinha, A., Claës, A., Preiser, P.R., Dedon, P.C., Baumgarten, S., Scherf, A., and Bryant, J.M. (2023). Cohesin contributes to transcriptional repression of stage-specific genes in the human malaria parasite. EMBO Rep. 24. 10.15252/embr.202357090.

57. Birnbaum, J., Scharf, S., Schmidt, S., Jonscher, E., Maria Hoeijmakers, W.A., Flemming, S., Toenhake, C.G., Schmitt, M., Sabitzki, R., Bergmann, B., et al. (2020). A Kelch13-defined endocytosis pathway mediates artemisinin resistance in malaria parasites. Science 367, 51–59. 10.1126/science.axx4735.

58. Roux, K.J., Kim, D.I., Raida, M., and Burke, B. (2012). A promiscuous biotin ligase fusion protein identifies proximal and interacting proteins in mammalian cells. J. Cell Biol. 196, 801–810. 10.1083/jcb.201112098.

59. Branon, T.C., Bosch, J.A., Sanchez, A.D., Udeshi, N.D., Svinkina, T., Carr, S.A., Feldman, J.L., Perrimon, N., and Ting, A.Y. (2018). Efficient proximity labeling in living cells and organisms with TurboID. Nat. Biotechnol. 36, 880–887. 10.1038/nbt.4201.

60. Birnbaum, J., Flemming, S., Reichard, N., Soares, A.B., Mesén-Ramírez, P., Jonscher, E., Bergmann, B., and Spielmann, T. (2017). A genetic system to study Plasmodium falciparum protein function. Nat. Methods 14, 450–456. 10.1038/nmeth.4223.

61. Gockel, J., Ramón-Zamorano, G., Kimmel, J., Spielmann, T., and Bártfai, R. (2025). CUT&Tag and DiBioCUT&Tag enable investigation of the AT-rich epigenome of Plasmodium falciparum from low-input samples. Cell Reports Methods, 101–110. 10.1016/j.crmeth.2025.101110.

62. Santos-Barriopedro, I., van Mierlo, G., and Vermeulen, M. (2022). Off-the-shelf proximity biotinylation using ProtA-TurboID. Nat. Protoc. 18, 36–57. 10.1038/s41596-022-00748-w.

63. Wichers, J.S., Wunderlich, J., Heincke, D., Pazicky, S., Strauss, J., Schmitt, M., Kimmel, J., Wilcke, L., Scharf, S., von Thien, H., et al. (2021). Identification of novel inner membrane complex and apical annuli proteins of the malaria parasite Plasmodium falciparum. Cell. Microbiol. 23. 10.1111/cmi.13341.

64. Kimmel, J., Schmitt, M., Sinner, A., Jansen, P.W.T.C., Mainye, S., Ramón-Zamorano, G., Toenhake, C.G., Wichers-Misterek, J.S., Cronshagen, J., Sabitzki, R., et al. (2023). Gene-by-gene screen of the unknown proteins encoded on Plasmodium falciparum chromosome 3. Cell Syst. 14, 9-23.e7. 10.1016/j.cels.2022.12.001.

65. Coleman, B.I., Skillman, K.M., Jiang, R.H.Y., Childs, L.M., Altenhofen, L.M., Ganter, M., Leung, Y., Goldowitz, I., Kafsack, B.F.C., Marti, M., et al. (2014). A Plasmodium falciparum histone deacetylase regulates antigenic variation and gametocyte conversion. Cell Host Microbe 16, 177–186. 10.1016/j.chom.2014.06.014.

66. Sethumadhavan, D.V., Tiburcio, M., Kanyal, A., Jabeena, C.A., Govindaraju, G., Karmodiya, K., and Rajavelu, A. (2022). Chromodomain Protein Interacts with H3K9me3 and Controls RBC Rosette Formation by Regulating the Expression of a Subset of RIFINs in the Malaria Parasite. J. Mol. Biol. 434, 167601. 10.1016/j.jmb.2022.167601.

67. Flueck, C., Bartfai, R., Niederwieser, I., Witmer, K., Alako, B.T.F., Moes, S., Bozdech, Z., Jenoe, P., Stunnenberg, H.G., and Voss, T.S. (2010). A major role for the Plasmodium falciparum ApiAP2 protein PfSIP2 in chromosome end biology. PLoS Pathog. 6. 10.1371/journal.ppat.1000784.

68. Sierra-Miranda, M., Vembar, S.S., Delgadillo, D.M., Ávila-López, P.A., Herrera-Solorio, A.M., Lozano Amado, D., Vargas, M., and Hernandez-Rivas, R. (2017). PfAP2Tel, harbouring a non-canonical DNA-binding AP2 domain, binds to Plasmodium falciparum telomeres. Cell. Microbiol. 19, e12742. 10.1111/cmi.12742.

69. Carrington, E., Cooijmans, R.H.M., Keller, D., Toenhake, C.G., Bártfai, R., and Voss, T.S. (2021). The ApiAP2 factor PfAP2-HC is an integral component of heterochromatin in the malaria parasite Plasmodium falciparum. iScience 24, 102444. 10.1016/j.isci.2021.102444.

70. Shang, X., Wang, C., Fan, Y., Guo, G., Wang, F., Zhao, Y., Sheng, F., Tang, J., He, X., Yu, X., et al. (2022). Genome-wide landscape of ApiAP2 transcription factors reveals a heterochromatin-associated regulatory network during Plasmodium falciparum blood-stage development. Nucleic Acids Res. 50, 3413–3431. 10.1093/nar/gkac176.

71. Aurrecoechea, C., Brestelli, J., Brunk, B.P., Dommer, J., Fischer, S., Gajria, B., Gao, X., Gingle, A., Grant, G., Harb, O.S., et al. (2009). PlasmoDB: a functional genomic database for malaria parasites. Nucleic Acids Res. 37, D539–D543. 10.1093/nar/gkn814.

72. Rosenberg, S.C., and Corbett, K.D. (2015). The multifaceted roles of the HORMA domain in cellular signaling. J. Cell Biol. 211, 745–755. 10.1083/jcb.201509076.

73. Ninova, M., Godneeva, B., Chen, Y.C.A., Luo, Y., Prakash, S.J., Jankovics, F., Erdélyi, M., Aravin, A.A., and Fejes Tóth, K. (2020). The SUMO Ligase Su(var)2-10 Controls Hetero- and Euchromatic Gene Expression via Establishing H3K9 Trimethylation and Negative Feedback Regulation. Mol. Cell 77, 571-585.e4. 10.1016/j.molcel.2019.09.033.

74. Zimmermann, L., Stephens, A., Nam, S.Z., Rau, D., Kübler, J., Lozajic, M., Gabler, F., Söding, J., Lupas, A.N., and Alva, V. (2018). A Completely Reimplemented MPI Bioinformatics Toolkit with a New HHpred Server at its Core. J. Mol. Biol. 430, 2237–2243. 10.1016/j.jmb.2017.12.007.

75. Ambekar, S. V., Beck, J.R., and Mair, G.R. (2022). TurboID Identification of Evolutionarily Divergent Components of the Nuclear Pore Complex in the Malaria Model Plasmodium berghei. MBio 13. 10.1128/mbio.01815-22.

76. Weiner, A., Dahan-Pasternak, N., Shimoni, E., Shinder, V., von Huth, P., Elbaum, M., and Dzikowski, R. (2011). 3D nuclear architecture reveals coupled cell cycle dynamics of chromatin and nuclear pores in the malaria parasite Plasmodium falciparum. Cell. Microbiol. 13, 967–977. 10.1111/j.1462-5822.2011.01592.x.

77. Iglesias, N., Paulo, J.A., Tatarakis, A., Wang, X., Edwards, A.L., Bhanu, N. V., Garcia, B.A., Haas, W., Gygi, S.P., and Moazed, D. (2020). Native Chromatin Proteomics Reveals a Role for Specific Nucleoporins in Heterochromatin Organization and Maintenance. Mol. Cell 77, 51-66.e8. 10.1016/j.molcel.2019.10.018.

78. Robinson, M.S., Sahlender, D.A., and Foster, S.D. (2010). Rapid Inactivation of Proteins by Rapamycin-Induced Rerouting to Mitochondria. Dev. Cell 18, 324. 10.1016/J.DEVCEL.2009.12.015.

79. Sinha, A., Hughes, K.R., Modrzynska, K.K., Otto, T.D., Pfander, C., Dickens, N.J., Religa, A.A., Bushell, E., Graham, A.L., Cameron, R., et al. (2014). A cascade of DNA-binding proteins for sexual commitment and development in Plasmodium. Nature 507, 253–257. 10.1038/nature12970.

80. Dogga, S.K., Rop, J.C., Cudini, J., Farr, E., Dara, A., Ouologuem, D., Djimdé, A.A., Talman, A.M., and Lawniczak, M.K.N. (2024). A single cell atlas of sexual development in Plasmodium falciparum. Science 384. 10.1126/science.adj4088.

81. Josling, G.A., Russell, T.J., Venezia, J., Orchard, L., van Biljon, R., Painter, H.J., and Llinás, M. (2020). Dissecting the role of PfAP2-G in malaria gametocytogenesis. Nat. Commun. 11. 10.1038/s41467-020-15026-0.

82. Miao, J., Wang, C., Lucky, A.B., Liang, X., Min, H., Adapa, S.R., Jiang, R., Kim, K., and Cui, L. (2021). A unique GCN5 histone acetyltransferase complex controls erythrocyte invasion and virulence in the malaria parasite Plasmodium falciparum. PLOS Pathog. 17, e1009351. 10.1371/journal.ppat.1009351.

83. Kalamuddin, M., Shakri, A.R., Wang, C., Min, H., Li, X., Cui, L., and Miao, J. (2024). MYST regulates DNA repair and forms a NuA4-like complex in the malaria parasite Plasmodium falciparum. mSphere 9. 10.1128/msphere.00140-24.

84. Petter, M., Lee, C.C., Byrne, T.J., Boysen, K.E., Volz, J., Ralph, S.A., Cowman, A.F., Brown, G. V., and Duffy, M.F. (2011). Expression of P. falciparum var Genes Involves Exchange of the Histone Variant H2A.Z at the Promoter. PLOS Pathog. 7, e1001292. 10.1371/journal.ppat.1001292.

85. Ranjan, A., Wang, F., Mizuguchi, G., Wei, D., Huang, Y., and Wu, C. (2015). H2A histone-fold and DNA elements in nucleosome activate SWR1-mediated H2A.Z replacement in budding yeast. Elife 4, 1–11. 10.7554/ELIFE.06845.

86. Liang, X., Shan, S., Pan, L., Zhao, J., Ranjan, A., Wang, F., Zhang, Z., Huang, Y., Feng, H., Wei, D., et al. (2016). Structural basis of H2A.Z recognition by SRCAP chromatin-remodeling subunit YL1. Nat. Struct. Mol. Biol. 23, 317–323. 10.1038/NSMB.3190,.

87. Liu, H., Cui, X.Y., Xu, D.D., Wang, F., Meng, L.W., Zhao, Y.M., Liu, M., Shen, S.J., He, X.H., Fang, Q., et al. (2020). Actin-related protein Arp4 regulates euchromatic gene expression and development through H2A.Z deposition in blood-stage Plasmodium falciparum. Parasites and Vectors 13, 1–15. 10.1186/S13071-020-04139-6.

88. Sun, L., and Luk, E. (2017). Dual function of Swc5 in SWR remodeling ATPase activation and histone H2A eviction. Nucleic Acids Res. 45, 9931–9946. 10.1093/nar/gkx589.

89. Brusini, L., Pacheco, N.D.S., Tromer, E.C., Soldati-Favre, D., and Brochet, M. (2022). Composition and organization of kinetochores show plasticity in apicomplexan chromosome segregation. J. Cell Biol. 221. 10.1083/jcb.202111084.

90. Hillier, C., Pardo, M., Yu, L., Bushell, E., Sanderson, T., Metcalf, T., Herd, C., Anar, B., Rayner, J.C., Billker, O., et al. (2019). Landscape of the Plasmodium Interactome Reveals Both Conserved and Species-Specific Functionality. Cell Rep. 28, 1635-1647.e5. 10.1016/j.celrep.2019.07.019.

91. Abramson, J., Adler, J., Dunger, J., Evans, R., Green, T., Pritzel, A., Ronneberger, O., Willmore, L., Ballard, A.J., Bambrick, J., et al. (2024). Accurate structure prediction of biomolecular interactions with AlphaFold 3. Nature 630, 493–500. 10.1038/s41586-024-07487-w.

92. van Kempen, M., Kim, S.S., Tumescheit, C., Mirdita, M., Lee, J., Gilchrist, C.L.M., Söding, J., and Steinegger, M. (2024). Fast and accurate protein structure search with Foldseek. Nat. Biotechnol. 42, 243–246. 10.1038/s41587-023-01773-0.

93. Kang, J., Yang, M., Li, B., Qi, W., Zhang, C., Shokat, K.M., Tomchick, D.R., Machius, M., and Yu, H. (2008). Structure and Substrate Recruitment of the Human Spindle Checkpoint Kinase Bub1. Mol. Cell 32, 394–405. 10.1016/j.molcel.2008.09.017.

94. Kim, T., and Gartner, A. (2021). Bub1 kinase in the regulation of mitosis. Animal Cells Syst. (Seoul). 25, 1–10. 10.1080/19768354.2021.1884599.

95. Zhang, C., Shine, M., Pyle, A.M., and Zhang, Y. (2022). US-align: universal structure alignments of proteins, nucleic acids, and macromolecular complexes. Nat. Methods 19, 1109–1115. 10.1038/s41592-022-01585-1.

96. Voß, Y., Klaus, S., Guizetti, J., and Ganter, M. (2023). Plasmodium schizogony, a chronology of the parasite’s cell cycle in the blood stage. PLOS Pathog. 19, pe1011157. 10.1371/journal.ppat.1011157.

97. Kops, G.J.P.L., Snel, B., and Tromer, E.C. (2020). Evolutionary Dynamics of the Spindle Assembly Checkpoint in Eukaryotes. Curr. Biol. 30, R589–R602. 10.1016/j.cub.2020.02.021.

98. Matthews, H., Duffy, C.W., and Merrick, C.J. (2018). Checks and balances? DNA replication and the cell cycle in Plasmodium. Parasites and Vectors 11. 10.1186/s13071-018-2800-1.

99. Machado, M., Klaus, S., Klaschka, D., Guizetti, J., and Ganter, M. (2023). Plasmodium falciparum CRK4 links early mitotic events to the onset of S-phase during schizogony. MBio 14. 10.1128/mbio.00779-23.

100. Hoyt, M.A., Totis, L., and Roberts, B.T. (1991). S. cerevisiae genes required for cell cycle arrest in response to loss of microtubule function. Cell 66, 507–517. 10.1016/0092-8674(81)90014-3.

101. Fréville, A., Moreira-Leite, F., Roussel, C., Russell, M.R.G., Fricot, A., Carret, V., Sissoko, A., Hayes, M.J., Diallo, A.B., Kerkhoven, N.C., et al. (2025). Malaria parasites undergo a rapid and extensive metamorphosis after invasion of the host erythrocyte. EMBO Rep. 26, 2545–2573. 10.1038/s44319-025-00435-3.

102. Grüring, C., Heiber, A., Kruse, F., Ungefehr, J., Gilberger, T.W., and Spielmann, T. (2011). Development and host cell modifications of Plasmodium falciparum blood stages in four dimensions. Nat. Commun. 2, 1–11. 10.1038/ncomms1169.

103. Kehrer, J., Frischknecht, F., and Mair, G.R. (2016). Proteomic analysis of the plasmodium berghei gametocyte egressome and vesicular bioid of osmiophilic body proteins identifies merozoite trap-like protein (MTRAP) as an essential factor for parasite transmission. Mol. Cell. Proteomics 15, 2852–2862. 10.1074/mcp.M116.058263.

104. Schnider, C.B., Bausch-Fluck, D., Brühlmann, F., Heussler, V.T., and Burda, P.-C. (2018). BioID Reveals Novel Proteins of the Plasmodium Parasitophorous Vacuole Membrane. mSphere 3. 10.1128/mSphere.00522-17.

105. Geiger, M., Brown, C., Wichers, J.S., Strauss, J., Lill, A., Thuenauer, R., Liffner, B., Wilcke, L., Lemcke, S., Heincke, D., et al. (2020). Structural Insights Into PfARO and Characterization of its Interaction With PfAIP. J. Mol. Biol. 432, 878–896. 10.1016/j.jmb.2019.12.024.

106. Boucher, M.J., Ghosh, S., Zhang, L., Lal, A., Jang, S.W., Ju, A., Zhang, S., Wang, X., Ralph, S.A., Zou, J., et al. (2018). Integrative proteomics and bioinformatic prediction enable a high-confidence apicoplast proteome in malaria parasites. PLOS Biol. 16, e2005895. 10.1371/journal.pbio.2005895.

107. Miao, J., Fan, Q., Cui, L., Li, X., Wang, H., Ning, G., Reese, J.C., and Cui, L. (2010). The MYST family histone acetyltransferase regulates gene expression and cell cycle in malaria parasite Plasmodium falciparum. Mol. Microbiol. 78, 883–902. 10.1111/j.1365-2958.2010.07371.x.

108. Bryant, J.M., Baumgarten, S., Dingli, F., Loew, D., Sinha, A., Claës, A., Preiser, P.R., Dedon, P.C., and Scherf, A. (2020). Exploring the virulence gene interactome with CRISPR/dCas9 in the human malaria parasite. Mol. Syst. Biol. 16, 9569. 10.15252/msb.20209569.

109. Zhang, Y., Song, C., Wang, L., Jiang, H., Zhai, Y., Wang, Y., Fang, J., and Zhang, G. (2022). Zombies Never Die: The Double Life Bub1 Lives in Mitosis. Front. Cell Dev. Biol. 10, 870745. 10.3389/FCELL.2022.870745/FULL.

110. Nagar, A., Yanase, R., Zeeshan, M., Ferguson, D.J.P., Abel, S., Pashley, S.L., Mishra, A., Eze, A., Rea, E., Brady, D., et al. (2025). Plasmodium ARK1 regulates spindle formation during atypical mitosis and forms a divergent chromosomal passenger complex. bioRxiv, 2025.08.27.672654. 10.1101/2025.08.27.672654.

111. Roques, M., Niu, C., Brochet, M., and Brusini, L. (2025). A Modular Chromosomal Passenger Complex Rewires Chromosome Segregation in Plasmodium berghei. bioRxiv, 2025.09.05.674402. 10.1101/2025.09.05.674402.

112. Sayers, C., Pandey, V., Balakrishnan, A., Michie, K., Svedberg, D., Hunziker, M., Pardo, M., Choudhary, J., Berntsson, R., and Billker, O. (2024). Systematic screens for fertility genes essential for malaria parasite transmission reveal conserved aspects of sex in a divergent eukaryote. Cell Syst. 15, 1075-1091.e6. 10.1016/j.cels.2024.10.008.

113. Behrens, H.M., and Spielmann, T. (2024). Identification of domains in Plasmodium falciparum proteins of unknown function using DALI search on AlphaFold predictions. Sci. Rep. 14, 1–12. 10.1038/s41598-024-60058-x.

114. Perez-Riverol, Y., Bandla, C., Kundu, D.J., Kamatchinathan, S., Bai, J., Hewapathirana, S., John, N.S., Prakash, A., Walzer, M., Wang, S., et al. (2025). The PRIDE database at 20 years: 2025 update. Nucleic Acids Res. 53, D543–D553. 10.1093/nar/gkae1011.

115. Edgar, R., Domrachev, M., and Lash, A.E. (2002). Gene Expression Omnibus: NCBI gene expression and hybridization array data repository. Nucleic Acids Res. 30, 207–210. 10.1093/nar/30.1.207.

116. Walliker, D., Quakyi, I.A., Wellems, T.E., McCutchan, T.F., Szarfman, A., London, W.T., Corcoran, L.M., Burkot, T.R., and Carter, R. (1987). Genetic analysis of the human malaria parasite Plasmodium falciparum. Science 236, 1661–1666. 10.1126/SCIENCE.3299700,.

117. Trager, W., and Jensen, J.B. (1976). Human malaria parasites in continuous culture. Science 193, 673–675. 10.1126/SCIENCE.781840,.

118. Gibson, D.G., Young, L., Chuang, R.Y., Venter, J.C., Hutchison, C.A., and Smith, H.O. (2009). Enzymatic assembly of DNA molecules up to several hundred kilobases. Nat. Methods 6, 343–345. 10.1038/NMETH.1318,.

119. Rivadeneira, E.M., Wasserman, M., and Espinal, C.T. (1983). Separation and Concentration of Schizonts of Plasmodium falciparum by Percoll Gradients. J. Protozool. 30, 367–370. 10.1111/j.1550-7408.1983.tb02932.x.

120. Hubner, N.C., Nguyen, L.N., Hornig, N.C., and Stunnenberg, H.G. (2015). A quantitative proteomics tool to identify DNA-protein interactions in primary cells or blood. J. Proteome Res. 14, 1315–1329. 10.1021/pr5009515.

121. Rappsilber, J., Mann, M., and Ishihama, Y. (2007). Protocol for micro-purification, enrichment, pre-fractionation and storage of peptides for proteomics using StageTips. Nat. Protoc. 2, 1896–1906. 10.1038/nprot.2007.261.

122. Boersema, P.J., Raijmakers, R., Lemeer, S., Mohammed, S., and Heck, A.J.R. (2009). Multiplex peptide stable isotope dimethyl labeling for quantitative proteomics. Nat. Protoc. 4, 484–494. 10.1038/nprot.2009.21.

123. Grüring, C., and Spielmann, T. (2012). Imaging of live malaria blood stage parasites. Methods Enzymol. 506, 81–92. 10.1016/B978-0-12-391856-7.00029-9.

124. Kaya-Okur, H.S., Wu, S.J., Codomo, C.A., Pledger, E.S., Bryson, T.D., Henikoff, J.G., Ahmad, K., and Henikoff, S. (2019). CUT&Tag for efficient epigenomic profiling of small samples and single cells. Nat. Commun. 10, 1–10. 10.1038/s41467-019-09982-5.

125. Buenrostro, J.D., Wu, B., Litzenburger, U.M., Ruff, D., Gonzales, M.L., Snyder, M.P., Chang, H.Y., and Greenleaf, W.J. (2015). Single-cell chromatin accessibility reveals principles of regulatory variation. Nature 523, 486–490. 10.1038/nature14590.

126. Kensche, P.R., Hoeijmakers, W.A.M., Toenhake, C.G., Bras, M., Chappell, L., Berriman, M., and Bártfai, R. (2015). The nucleosome landscape of Plasmodium falciparum reveals chromatin architecture and dynamics of regulatory sequences. Nucleic Acids Res. 44, 2110–2124. 10.1093/nar/gkv1214.

127. Spielmann, T., Fergusen, D.J.P., and Beck, H.P. (2003). etramps, a new Plasmodium falciparum gene family coding for developmentally regulated and highly charged membrane proteins located at the parasite-host cell interface. Mol. Biol. Cell 14, 1529–1544. 10.1091/mbc.e02-04-0240.

128. Mesén-Ramírez, P., Reinsch, F., Blancke Soares, A., Bergmann, B., Ullrich, A.K., Tenzer, S., and Spielmann, T. (2016). Stable Translocation Intermediates Jam Global Protein Export in Plasmodium falciparum Parasites and Link the PTEX Component EXP2 with Translocation Activity. PLOS Pathog. 12, e1005618. 10.1371/journal.ppat.1005618.

129. Liffner, B., and Absalon, S. (2021). Expansion microscopy reveals Plasmodium falciparum blood-stage parasites undergo anaphase with a chromatin bridge in the absence of mini-chromosome maintenance complex binding protein. Microorganisms 9. 10.3390/microorganisms9112306.

130. Cox, J., and Mann, M. (2008). MaxQuant enables high peptide identification rates, individualized p.p.b.-range mass accuracies and proteome-wide protein quantification. Nat. Biotechnol. 26, 1367–1372. 10.1038/nbt.1511.

131. Tyanova, S., Temu, T., Sinitcyn, P., Carlson, A., Hein, M.Y., Geiger, T., Mann, M., and Cox, J. (2016). The Perseus computational platform for comprehensive analysis of (prote)omics data. Nat. Methods 13, 731–740. 10.1038/NMETH.3901,.

132. Hohjoh, H., and Singer, M.F. (1997). Sequence-specific single-strand RNA binding protein encoded by the human LINE-1 retrotransposon. EMBO J. 16, 6034–6043.

133. Li, H., Handsaker, B., Wysoker, A., Fennell, T., Ruan, J., Homer, N., Marth, G., Abecasis, G., and Durbin, R. (2009). The Sequence Alignment/Map format and SAMtools. Bioinformatics 25, 2078–2079. 10.1093/BIOINFORMATICS/BTP352,.

134. Ramírez, F., Ryan, D.P., Grüning, B., Bhardwaj, V., Kilpert, F., Richter, A.S., Heyne, S., Dündar, F., and Manke, T. (2016). deepTools2: a next generation web server for deep-sequencing data analysis. Nucleic Acids Res. 44, W160–W165. 10.1093/nar/gkw257.

135. Perez, G., Barber, G.P., Benet-Pages, A., Casper, J., Clawson, H., Diekhans, M., Fischer, C., Gonzalez, J.N., Hinrichs, A.S., Lee, C.M., et al. (2025). The UCSC Genome Browser database: 2025 update. Nucleic Acids Res. 53, D1243–D1249. 10.1093/nar/gkae974.

136. Quinlan, A.R., and Hall, I.M. (2010). BEDTools: A flexible suite of utilities for comparing genomic features. Bioinformatics 26, 841–842. 10.1093/BIOINFORMATICS/BTQ033,.

137. Dobin, A., Davis, C.A., Schlesinger, F., Drenkow, J., Zaleski, C., Jha, S., Batut, P., Chaisson, M., and Gingeras, T.R. (2012). STAR: ultrafast universal RNA-seq aligner. Bioinformatics 29, 15. 10.1093/BIOINFORMATICS/BTS635.

138. Liao, Y., Smyth, G.K., and Shi, W. (2014). FeatureCounts: An efficient general purpose program for assigning sequence reads to genomic features. Bioinformatics 30, 923–930. 10.1093/bioinformatics/btt656.

139. Love, M.I., Huber, W., and Anders, S. (2014). Moderated estimation of fold change and dispersion for RNA-seq data with DESeq2. Genome Biol. 15, 1–21. 10.1186/s13059-014-0550-8.

140. Wickham, H. (2016). ggplot2: Elegant Graphics for Data Analysis (Springer International Publishing) 10.1007/978-3-319-24277-4.

141. Morpheus https://software.broadinstitute.org/morpheus/.

142. Feng, Y., Tian, Y., Wu, Z., and Xu, Y. (2018). Cryo-EM structure of human SRCAP complex. Cell Res. 28, 1121–1123. 10.1038/s41422-018-0102-y.

143. Yu, J., Sui, F., Gu, F., Li, W., Yu, Z., Wang, Q., He, S., Wang, L., and Xu, Y. (2024). Structural insights into histone exchange by human SRCAP complex. Cell Discov. 10, 1–18. 10.1038/s41421-023-00640-1.

