## Supplementary Figures for "Protein landscape of the chromatin states in the malaria parasite *Plasmodium falciparum*"

### Author affiliation and footnotes

[1] Bernhard Nocht Institute for Tropical Medicine, Hamburg, Germany
[2] Department of Molecular Biology, Radboud Institute for Molecular Life Sciences, Radboud University, Nijmegen, the Netherlands
[3] Department of Molecular Biology, Radboud Institute for Molecular Life Sciences, Oncode Institute, Radboud University Nijmegen, Nijmegen, the Netherlands
[4] Mikrobiologisches Institut - Klinische Mikrobiologie, Immunologie und Hygiene, Universitätsklinikum Erlangen, Friedrich-Alexander-Universität (FAU) Erlangen-Nürnberg, Erlangen, Germany
[5] Division of Molecular Genetics, Netherlands Cancer Institute, Amsterdam, Netherlands
[6] These authors contributed equally
[7] These authors contributed equally
[8] Lead contact

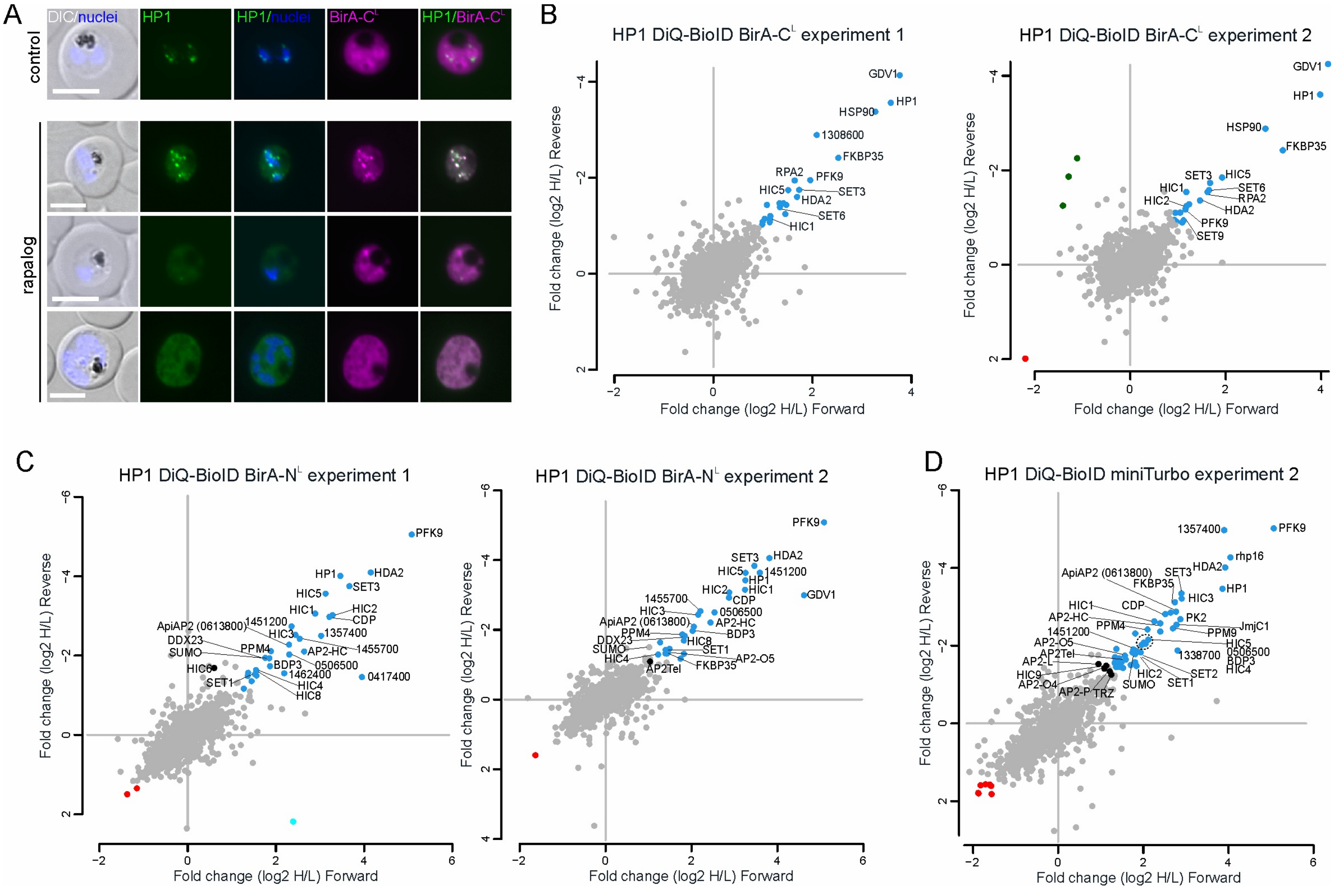


**Figure S1**. **Additional data of HP1 (mT)-DiQ-BioID experiments**. **(A)** Live fluorescence microscopy images showing recruitment of the biotin ligase BirA-C$^L$ to HP1 upon rapalog addition (rapalog) compared to control (cytoplasmic distribution). Representative images of 2 biological replicates with 44 (control) and 64 (rapalog) parasites analyzed per condition and session. **(B-D)** Full scatter plots showing log2 fold change of rapalog over control (H/L) from all individual BirA-C$^L$ (B), BirA-N$^L$ (C) and miniTurbo-(D)DiQ-BioID experiments with technical replicates (Forward and Reverse) plotted against each other. The mT-DiQ-BioID experiment 1 contains only one technical replicate and therefore was not plotted. Outliers (beyond 1.5x interquartile range) identified in both technical replicates are marked with their abbreviation (if available) or gene ID (excluding “PF3D7_”). Scale bars, 5 µm; DIC, differential interference contrast; nuclei (nuclei) were stained with Hoechst 33342.

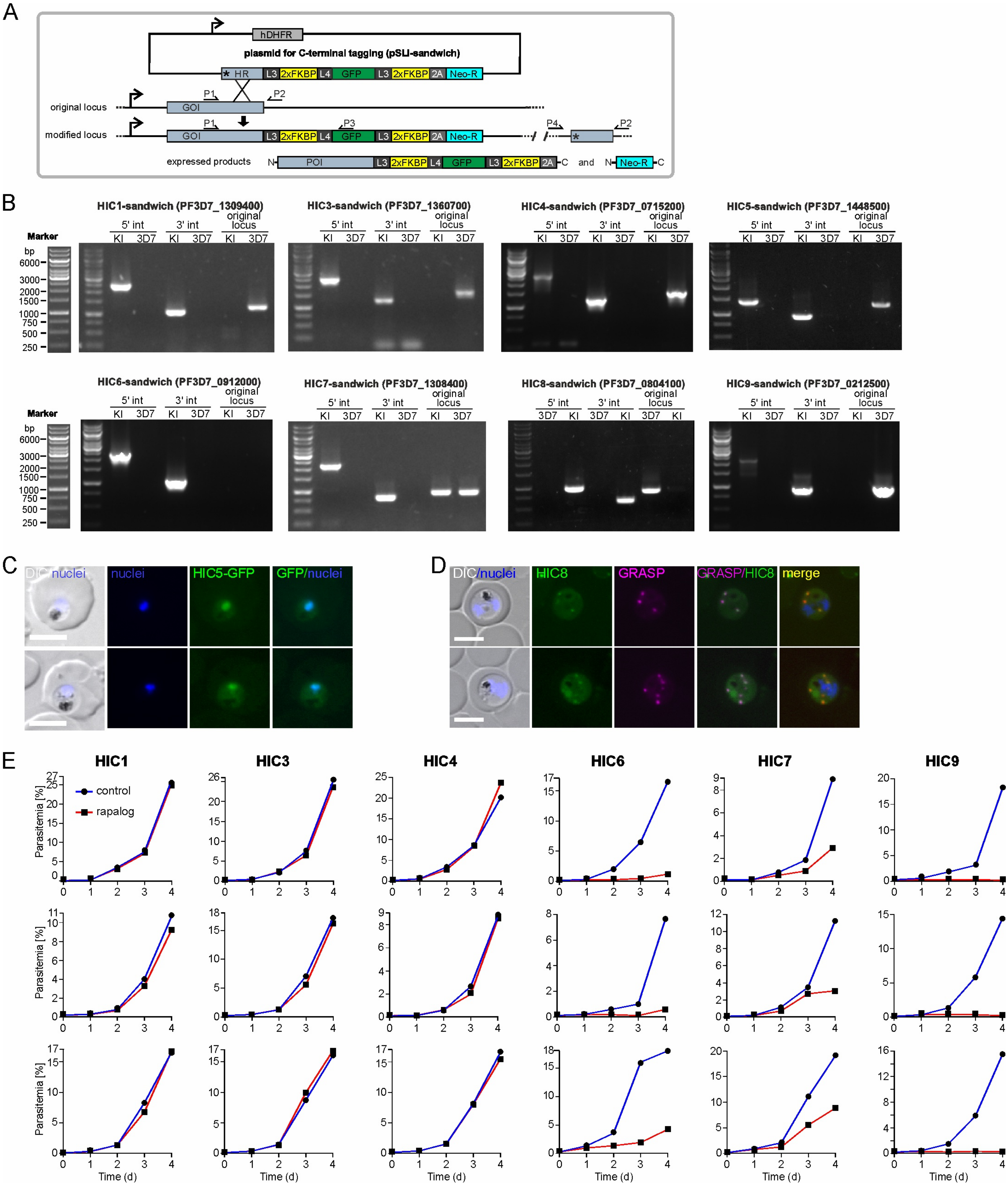


**Figure S2**. **Additional data for HICs. (A)** Schematics of the selection-linked integration (SLI)[60] strategy for genome modification. Analyzed proteins in this study were endogenously tagged with 2xFKBP-GFP-2xFKBP tag introduced by the integration of the pSLI-sandwich plasmid (top) via the homology region (HR) of the gene of interest (GOI) in the genome, resulting in the expression of the SLI-resistance (Neo-R (conferring G418 resistance)) and protein of interest (POI) with the tag (bottom). *, STOP codon; L3 and L4, linkers; 2A, T2A skip peptide; Neo-R, G418 resistance; hDHFR, human dihydrofolate reductase. Small arrows indicate position of primers used for assessing correct integration of the plasmid into the genome shown in (B). **(B)** Agarose gels with PCR products amplified from genomic DNA of HIC-sandwich knock in lines (KI) compared to 3D7 wildtype parasites (3D7) confirming the genomic integration of pSLI-sandwich plasmid into the correct locus. PCR products show 5' integration (5' int) using primers P1 and P3, 3' integration (3' int) using primers P4 and P2 and absence of the unmodified original locus using primers P1 and P2. Primer positions are indicated in (A) and primer sequences are in Table S8. Marker fragment length is indicated in base pairs on the left.

**(C)** Live fluorescence microscopy images of parasites expressing endogenously tagged HIC5, showing HIC5 nuclear distribution. **(D)** Fluorescence microscopy images of parasites expressing endogenously tagged HIC8 and episomally co-expressing GRASP-mCherry, showing HIC8 co-localization with GRASP. **(E)** Flow cytometry growth curves of all three biological replicates of HICs-KS for which one is also shown in the main figures. Scale bars, 5 µm; DIC, differential interference contrast; nuclei (nuclei) were stained with Hoechst 33342.

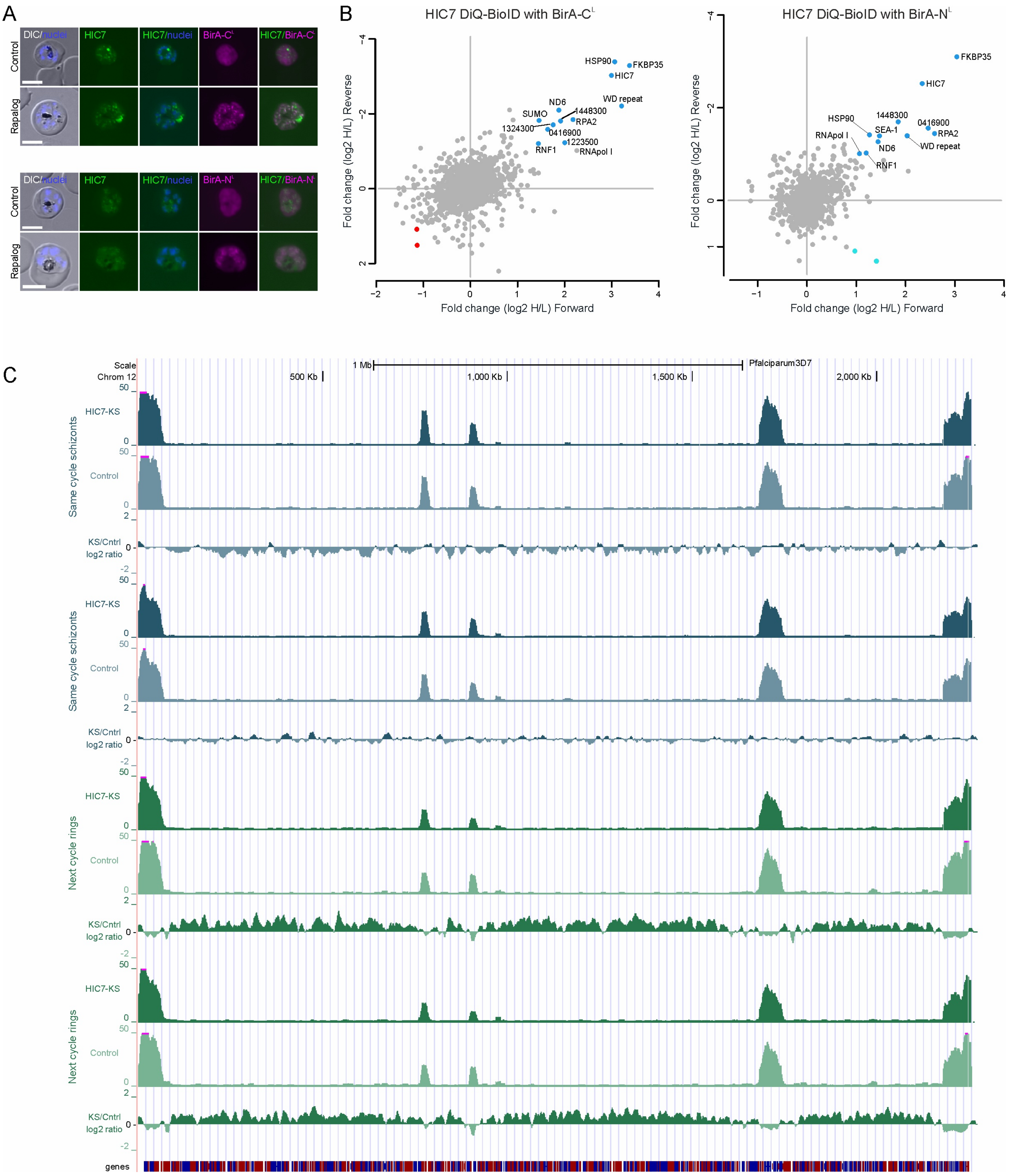


**Figure S3**. **Additional data for HIC7**. **(A)** Live fluorescence microscopy showing recruitment of the biotin ligase BirA-$C^L$ (top panel) and BirA-$N^L$ (bottom panel) to HIC7 upon rapalog addition (rapalog) compared to cytoplasmic distribution in control parasites (control). Representative images are shown from independent experiments in which 10 (control) and 12 (rapalog) fields of view were analyzed for the BirA-$C^L$ experiment, and 6 (control) and 9 (rapalog) fields of view were analyzed for the BirA-$N^L$ experiment. **(B)** Scatter plots of HIC7 DiQ-BioID experiments using BirA*-$C^L$ (left) or BirA*-$N^L$ (right) showing log2 fold enrichment of 2 technical replicates (Forward and Reverse) plotted against each other. Outliers (beyond 1.5x interquartile range) identified in both technical replicates are highlighted in blue and are marked with their abbreviation (if available) or gene ID (excluding “PF3D7_”). **(C)** Genome browser tracks of HP1 occupancy on chromosome 12 from CUT&Tag data. Pair-wise comparison of HP1 CUT&Tag tag counts (top 2 tracks each) and fold change (log2 ratio, bottom track each) in schizont (blue tracks) and ring-stage (green tracks) parasites wherein HIC7-KS was induced (KS) compared to control parasites (control) from 2 biological replicates each is shown. Scale bars, 5 µm; DIC, differential interference contrast; nuclei were stained with Hoechst 33342.

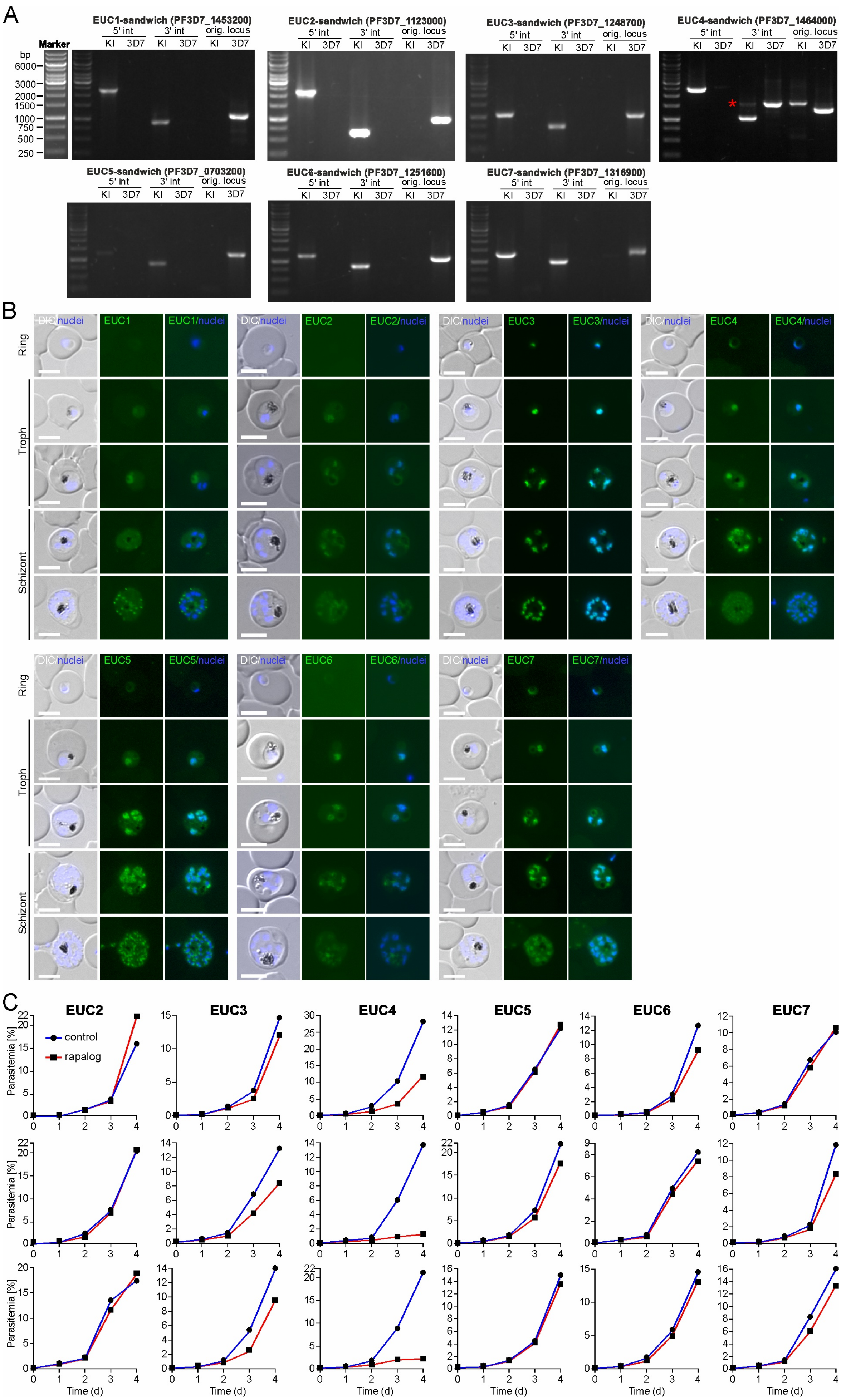


**Figure S4. Additional data for EUCs. (A)** Agarose gels with PCR products amplified from genomic DNA of EUC-sandwich knock in lines (KI) compared to 3D7 wildtype parasites (3D7) confirming the genomic integration of pSLI-sandwich plasmid into correct EUCs loci. PCR products show 5' integration (5' int) using primers P1 and P3, 3' integration (3' int) using primers P4 and P2 or absence of the unmodified original locus using primers P1 and P2. A red asterisk (*) marks unspecific PCR products. Primer positions are indicated in Figure S2A and primer sequences are in Table S8. Marker fragment length is indicated in base pairs on the top left. **(B)** Full panels of live fluorescence microscopy images of parasites expressing endogenously tagged EUCs showing their expression through the asexual parasite development (ring, trophozoite and schizonts) (see also Figure 3C). For each EUC representative images of at least 7 microscopy sessions including an average of 6 fields of view in each are shown. **(C)** Flow cytometry growth curves of all three biological replicates of EUC-KS (rapalog, red line) and control parasites (blue line) shown in Figure 3D. Scale bars, 5 µm; DIC, differential interference contrast; nuclei were stained with Hoechst 33342.

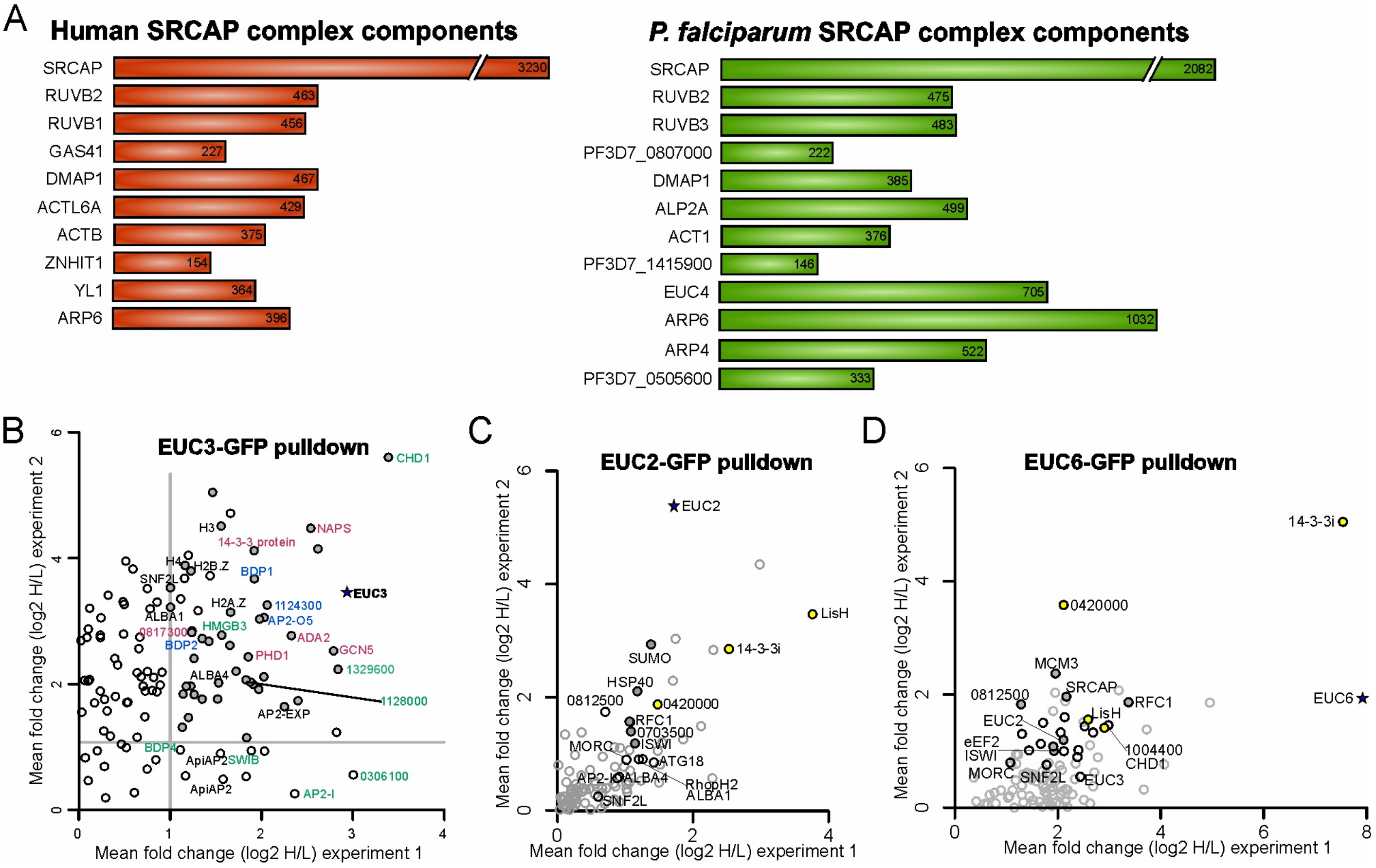


**Figure S5**. **Interacting proteins of EUC4, EUC3, EUC2 and EUC6**. **(A)** SRCAP complex components in human (left,[144,145]) and P. falciparum (right, this study). Names or gene IDs and length (in amino acids) are indicated. **(B-D)** Top-right quadrant of scatterplots of EUC3-GFP (B), EUC2-GFP (C) and EUC6-GFP (D) pull downs showing mean fold-change enrichment of 2 biological replicates (including 2 technical replicates each) plotted against each other. Enriched proteins in both experiments are marked with their abbreviation (if available) or gene ID (excluding “PF3D7_”). Bait proteins are highlighted with a star symbol. In (B), epigenetic reader proteins are named and colour-coded according to the complexes described by Hoeijmakers et al[36]. In (C) and (D), proteins that belong to the EUC2/EUC6 complex are highlighted with a yellow dot.

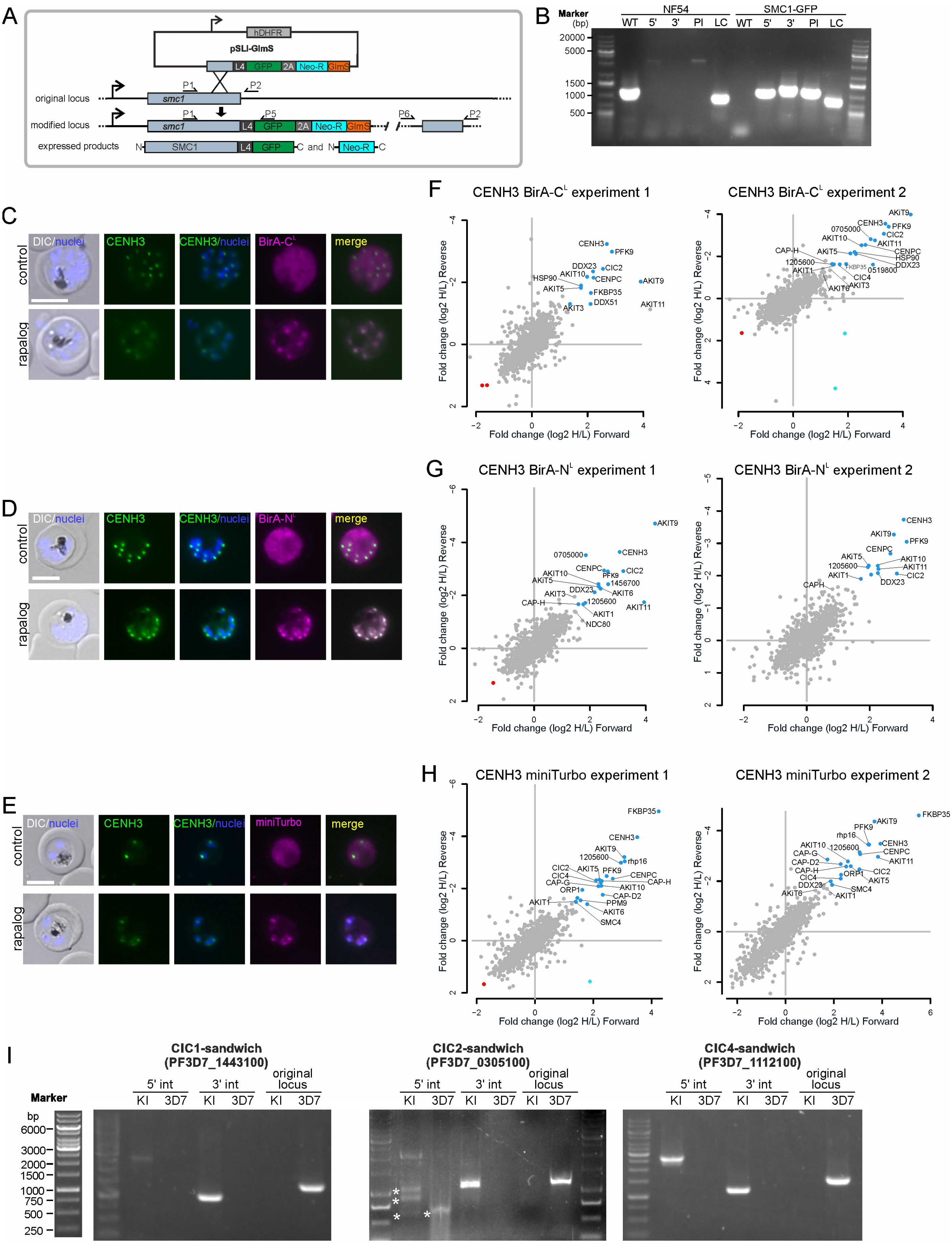


**Figure S6**. **Additional data for SMC1-targeting ProtA-TurboID and CENH3 (mT)-DiQ-BioID experiments. (A)** Schematics of the selection-linked integration (SLI)[60] strategy for genome modification of SMC1. SMC1 was endogenously tagged with GFP introduced by the integration of the pSLI-SMC1-glmS plasmid (top) into the homology region (HR) of the *smc1* genomic locus, resulting in the expression of the GFP tagged SMC1 protein, the SLI-resistance (Neo-R, conferring resistance to G418) and the glmS ribozyme (bottom). L4, linker; 2A, T2A skip peptide; Neo-R, G418 resistance; hDHFR, human dihydrofolate reductase. Small arrows indicate position of primers used for assessing correct integration of the plasmid into the genome shown in (B). **(B)** Agarose gel with PCR products amplified from genomic DNA of the SMC1-GFP-glmS knock in parasite line (KI) compared to NF54 wildtype parasites (NF54) confirming correct modification of the *smc1* genomic locus. PCR products show the unmodified locus (WT) using primers P1 and P2, 5' integration (5') using primers P1 and P5, 3' integration (3') using primers P6 and P2 or the plasmid (Pl) using primers P5 and P6. LC is *Pfs16* genomic amplification as loading control. Primer positions are indicated in (A) and primer sequences in Table S8. Plasmid sequences are shown in File S1. Marker fragment length is indicated in base pairs. **(C-E)** Live fluorescence microscopy showing recruitment of the biotin ligase BirA-$C^L$ (C), BirA-$N^L$ (D) or miniTurbo (E) to CENH3 upon rapalog addition (rapalog) compared to control (cytoplasmic distribution). Representative images from 2 biological replicates with an average of 6 fields of view each. **(F-H)** Full scatterplots showing log2 fold change of rapalog over control from all individual BirA-$C^L$ (F), BirA-$N^L$ (G) and miniTurbo-DiQ-BioID (H) experiments from Figure 4D with technical replicates (Forward and Reverse) plotted against each other. Outliers (beyond 1.5x interquartile range) identified in both technical replicates are highlighted in blue (up) and red (down) and marked with their abbreviation (if available) or gene ID (excluding "PF3D7_"). **(I)** Agarose gels with PCR products amplified from genomic DNA of CICs-sandwich knock in lines (KI) compared to 3D7 wildtype parasites (3D7) confirming the genomic integration of pSLI-sandwich plasmid into correct CICs loci. PCR products show 5' integration (5' int) using primers P1 and P3, 3' integration (3' int) using primers P4 and P2 or absence of the unmodified original locus using primers P1 and P2. Primer positions are indicated in Figure S2A and primer sequences are in Table S8. Marker fragment length is indicated in base pairs on the top left. Asterisks shows non-specific bands. Scale bars, 5 µm; DIC, differential interference contrast; nuclei were stained with Hoechst 33342.

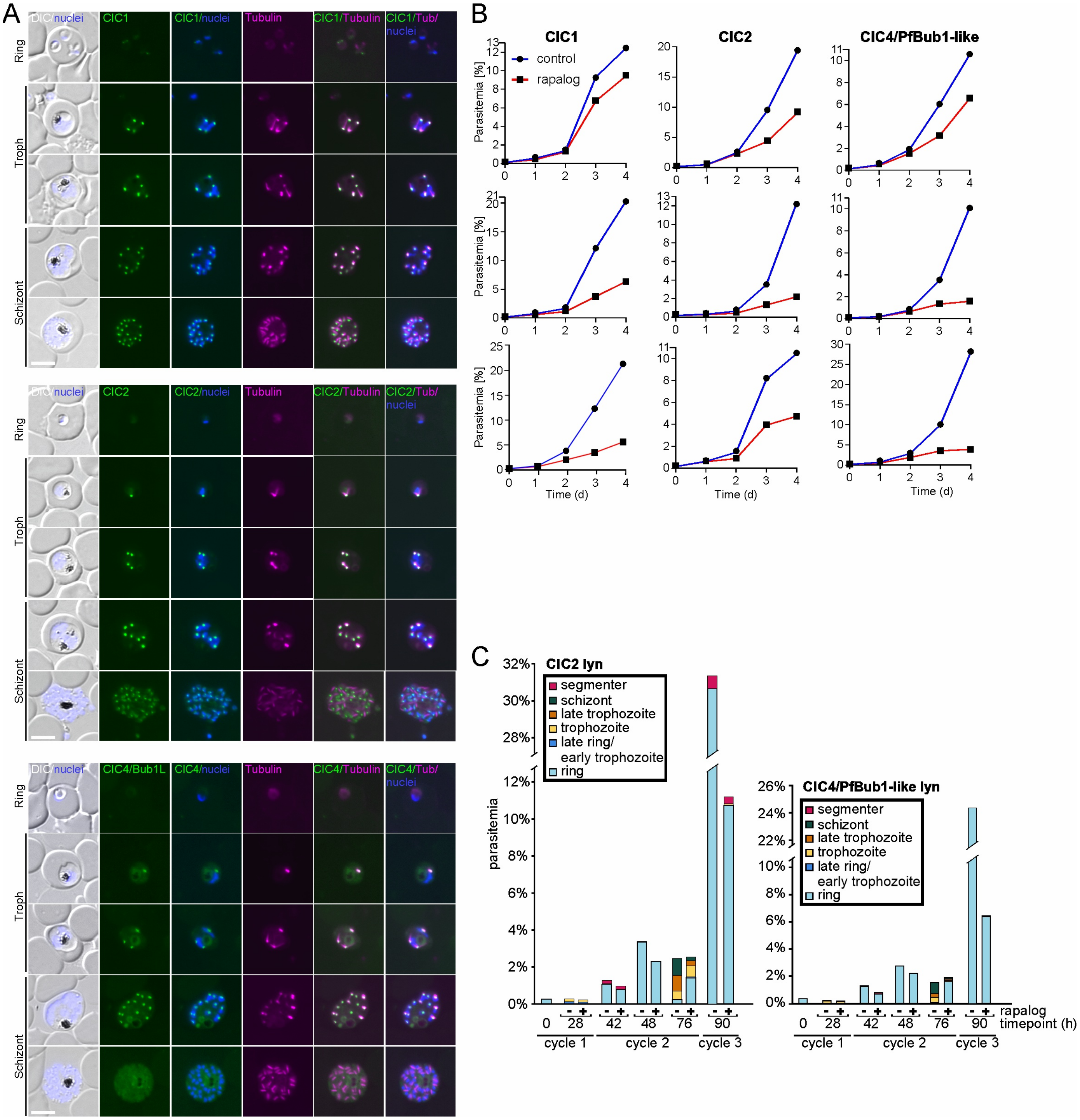


**Figure S7**. **Additional data for ClCs**. **(A)** Full panels of live fluorescence microscopy images of parasites expressing endogenously tagged ClCs showing their expression and association with tubulin (stained with Tubulin Tracker Deep Red) through asexual parasite development (ring, trophozoite and schizonts). Representative images of at least 5 (ClC1), 4 (ClC2) or 6 (ClC4/PfBub1-like) independent experiments including an average of 7 fields of view per condition and session. **(B)** Flow cytometry growth curves of all three biological replicates of ClCs-KS (rapalog, red line) and control parasites (blue line) shown in Figure 4H. **(C)** Second biological replicates (first replicates in Figure 4I) of stages and growth based on Giemsa smears in synchronised (0-2 hours synchronisation window) parasites expressing endogenously tagged ClC2 and ClC4/PfBub1-like and episomally co-expressing Lyn-FRB-mCherry grown in presence (+rapalog, to induce ClCs-KS) and absence of rapalog (- rapalog) at the indicated time points. Scale bars, 5 µm; DIC, differential interference contrast; nuclei (nuclei) were stained with Hoechst 33342.
