## Supplementary material for "Protein landscape of the chromatin states in the malaria parasite *Plasmodium falciparum*": Suppl. file S1. Plasmid sequences

**>pSLI-sandwich**

XXXX = hDHFR resistance

INSERT = GOI homology region

XXXX = Linker

XXXX = FKBP

XXXX = GFP

XXXX = T2A skip peptide

XXXX = G418 resistance

ggatatggcagcttaatgttcgtttttcttatttatatatttataccaattgattgtatttataactgtaaaaatgtgtatgttgtgtgcatatttttttttgtgcatgcacatgcatgtaaatagctaaaattatgaacattttattttttgttcagaaaaaaaaaactttacacacataaaatggctagtatgaatagccatattttatataaattaaatcctatgaatttatgaccatattaaaaatttagatatttatggaacataatatgtttgaaacaataagacaaaattattattattattattatttttactgttataattatgtgtctccttcaatgattcataaatagttggacttgatttttaaaatgtttataatatgattagcatagttaaataaaaaaagttgaaaaattaaaaaaaaacatataaacacaaatgatggtttttccttcaatttcgatatcaatttatagaaacaaaatatatacttgtataattttatttttttatataaatcattacatatataattatacaatattttttctaagagataattatatattaatatatataaaaaaaggtgttttttttttttttttttatttttatttttattttatggtaatattttattttccttattttataaattatattagtttatatgtgattaattttatatattatcaatttatatatttttaaatgcttacttaattatctttttttttttttttttttttttttcccctctttttatattaatttatttttgaaaaaattgatatatatatatatatataatatatatatatacatgtagtagtattaaacaatgtataatatatataaataatatatttatatatttcatttcaattttaattttttttggttttttttttttttctttttgtcatatttaaaaaaaattatattcatataagttatgcattttttataaacattattcaatatatgtataatataatatatatatatatattaatgtattattccaatgtgcatgataaaagaaaaaaataatatttataaaaaaaaagaaaaataaaacaaaaaaagaaaaaaaaaaaaaaaaaaaaaaaaatacaaaaataaataatataatttataattatatattcttgtcacaataaaaatatatatatatatatatatatttataatatgtatattttaaactagaaaaggaataactaatattttatttattatcattcaagatttatattttataataataaatacctaatagaaatatatcaggatccatgcatggttcgctaaactgcatcgtcgctgtgtcccagaacatgggcatcggcaagaacggggactacccctggccaccgctcaggaacgaatttagatatttccagagaatgaccacaacctcttcagtagaaggtaaacagaatctggtgattatgggtaagaagacctggttctccattcctgagaagaatcgacctttaaagggtagaattaatttagttctcagcagagaactcaaggaacctccacaaggagctcattttctttccagaagtctagatgatgccttaaaacttactgaacaaccagaattagcaaataaagtagacatggtctggatagttggtggcagttctgtttataaggaagccatgaatcacccaggccatcttaaactatttgtgacaaggatcatgcaagactttgaaagtgacacgttttttccagaaattgatttggagaaatataaacttctgccagaatacccaggtgttctctctgatgtccaggaggagaaaggcattaagtacaaatttgaagtatatgagaagaatgattaagcttatttaataatagattaaaaatattataaaaataaaaacataaacacagaaattacaaaaaaaatacatatgaattttttttttgtaatcttccttataaatatagaataatgaatcatataaaacatatcattattcatttatttacatttaaaattattgtttcagtatctttaatttattatgtatatataaaaataacttacaattttattaataaacaatatatgtttattaattcatgttttgtaatttatgggatagcgattttttttactgtctgtatttttcttttttaattatgttttaattgtattttatttttattattgttctttttatagtattattttaaaacaaaatgtattttctaagaacttataataataataaatataaattttaataaaaattatatttatcttttacaatatgaacataaagtacaacattaatatatagcttttaatatttttattcctaatcatgtaaatcttaaatttttctttttaaacatatgttaaatatttatttctcattatatataagaacatatttattaaatctagaattctatagtgagtcgtattacaattcactggccgtcgttttacaacgtcgtgactgggaaaaccctggcgttacccaacttaatcgccttgcagcacatccccctttcgccagctggcgtaatagcgaagaggcccgcaccgatcgcccttcccaacagttgcgcagcctgaatggcgaatggcgcctgatgcggtattttctccttacgcatctgtgcggtatttcacaccgcatatggtgcactctcagtacaatctgctctgatgccgcatagttaagccagccccgacacccgccaacacccgctgacgcgccctgacgggcttgtctgctcccggcatccgcttacagacaagctgtgaccgtctccgggagctgcatgtgtcagaggttttcaccgtcatcaccgaaacgcgcgagacgaaagggcctcgtgatacgcctatttttataggttaatgtcatgataataatggtttcttagacgtcaggtggcacttttcggggaaatgtgcgcggaacccctatttgtttatttttctaaatacattcaaatatgtatccgctcatgagacaataaccctgataaatgcttcaataatattgaaaaaggaagagtatgagtattcaacatttccgtgtcgcccttattcccttttttgcggcattttgccttcctgtttttgctcacccagaaacgctggtgaaagtaaaagatgctgaagatcagttgggtgcacgagtgggttacatcgaactggatctcaacagcggtaagatccttgagagttttcgccccgaagaacgttttccaatgatgagcacttttaaagttctgctatgtggcgcggtattatcccgtattgacgccgggcaagagcaactcggtcgccgcatacactattctcagaatgacttggttgagtactcaccagtcacagaaaagcatcttacggatggcatgacagtaagagaattatgcagtgctgccataaccatgagtgataacactgcggccaacttacttctgacaacgatcggaggaccgaaggagctaaccgcttttttgcacaacatgggggatcatgtaactcgccttgatcgttgggaaccggagctgaatgaagccataccaaacgacgagcgtgacaccacgatgcctgtagcaatgccaacaacgttgcgcaaactattaactggcgaactacttactctagcttcccggcaacaattaatagactggatggaggcggataaagttgcaggaccacttctgcgctcggcccttccggctggctggtttattgctgataaatctggagccggtgagcgtgggtctcgcggtatcattgcagcactggggccagatggtaagccctcccgtatcgtagttatctacacgacggggagtcaggcaactatggatgaacgaaatagacagatcgctgagataggtgcctcactgattaagcattggtaactgtcagaccaagtttactcatatatactttagattgatttaaaacttcatttttaatttaaaaggatctaggtgaagatcctttttgataatctcatgaccaaaatcccttaacgtgagttttcgttccactgagcgtcagaccccgtagaaaagatcaaaggatcttcttgagatcctttttttctgcgcgtaatctgctgcttgcaaacaaaaaaaccaccgctaccagcggtggtttgtttgccggatcaagagctaccaactctttttccgaaggtaactggcttcagcagagcgcagataccaaatactgtccttctagtgtagccgtagttaggccaccacttcaagaactctgtagcaccgcctacatacctcgctctgctaatcctgttaccagtggctgctgccagtggcgataagtcgtgtcttaccgggttggactcaagacgatagttaccggataaggcgcagcggtcgggctgaacggggggttcgtgcacacagcccagcttggagcgaacgacctacaccgaactgagatacctacagcgtgagctatgagaaagcgccacgcttcccgaagggagaaaggcggacaggtatccggtaagcggcagggtcggaacaggagagcgcacgagggagcttccagggggaaacgcctggtatctttatagtcctgtcgggtttcgccacctctgacttgagcgtcgatttttgtgatgctcgtcaggggggcggagcctatcgaaaaacgccagcaacgcggcctttttacggttcctggccttttgctggccttttgctcacatgttctttcctgcgttatcccctgattctgtggataaccgtattaccgcctttgagtgagctgataccgctcgccgcagccgaacgaccgagcgcagcgagtcagtgagcgaggaagcggaagagcgcccaatacgcaaaccgcctctccccgcgcgttggccgattcattaatgcagctggcacgacaggtttcccgactggaaagcgggcagtgagcgcaacgcaattaatgtgagttagctcactcattaggcaccccaggctttacactttatgcttccggctcgtatgttgtgtggaattgtgagcggataacaatttcacacaggaaacagctatgaccatgattacgccaagctatttaggtgacactatagaatactcgcggccgctaa**INSERT**cctaggTCAGGATTGAGATCAAGATCTGCTGCTGCTGGTGCTGGTGGTGCTGCTAGAGCTGCTctgcagAGAGGAGTACAAGTTGAAACAATATCACCAGGAGATGGTCGTACATTTCCAAAAAGAGGTCAAACTTGTGTTGTACATTATACTGGAATGCTTGAAGATGGAAAGAAATTTGATTCATCTCGTGATAGAAATAAACCATTTAAATTTATGCTAGGTAAACAAGAAGTAATACGAGGTTGGGAAGAAGGAGTTGCTCAAATGAGTGTAGGTCAAAGAGCAAAACTTACTATATCTCCAGATTATGCTTATGGTGCAACTGGACATCCAGGTATAATTCCACCTCATGCAACTCTTGTATTTGATGTGGAGCTTCTAAAACTAGAAACTAGAGGTGTTCAGGTTGAAACAATTTCACCTGGAGATGGCAGAACCTTTCCTAAAAGAGGACAGACTTGCGTAGTTCATTATACAGGCATGCTAGAGGATGGTAAGAAATTTGATTCTAGTCGAGATAGAAATAAGCCATTCAAGTTTATGCTAGGTAAACAGGAAGTAATAAGAGGTTGGGAAGAGGGTGTAGCACAGATGTCAGTTGGACAAAGAGCAAAGTTAACAATATCACCAGATTATGCATACGGTGCAACAGGCCATCCTGGCATCATCCCTCCACATGCAACTTTAGTATTCGACGTTGAATTGTTAAAGTTAGAGACAacgcgtGCTAGAGGTGCTGCTGCTGGTGCTGGAGGTGCAGGTAGACGTACGATGAGTAAAGGAGAAGAACTTTTCACTGGAGTTGTCCCAATTCTTGTTGAATTAGATGGTGATGTTAATGGGCACAAATTTTCTGTCAGTGGAGAGGGTGAAGGTGATGCAACATACGGAAAACTTACCCTTAAATTTATTTGCACTACTGGAAAACTACCTGTTCCATGGCCAACACTTGTCACTACTTTCGCGTATGGTCTTCAATGCTTTGCGAGATACCCAGATCATATGAAACAGCATGACTTTTTCAAGAGTGCCATGCCCGAAGGTTATGTACAGGAAAGAACTATATTTTTCAAAGATGACGGGAACTACAAGACACGTGCTGAAGTCAAGTTTGAAGGTGATACCCTTGTTAATAGAATCGAGTTAAAAGGTATTGATTTTAAAGAAGATGGAAACATTCTTGGACACAAATTGGAATACAACTATAACTCACACAATGTATACATCATGGCAGACAAACAAAAGAATGGAATCAAAGTTAACTTCAAAATTAGACACAACATTGAAGATGGAAGCGTTCAACTAGCAGACCATTATCAACAAAATACTCCAATTGGCGATGGCCCTGTCCTTTTACCAGACAACCATTACCTGTCCACACAATCTGCCCTTTCGAAAGATCCCAACGAAAAGAGAGACCACATGGTCCTTCTTGAGTTTGTAACAGCTGCTGGGATTACACATGGCATGGATGAGCTCTACAAAGTCGACGCCAGGGGAGCAGCCGCAGGAGCAGGGGGGGCAGGAAGGCGTGGTGTTCAGGTCGAGACTATTAGCCCTGGAGATGGACGCACGTTTCCTAAGCGTGGACAGACATGCGTAGTTCACTACACAGGTATGTTGGAGGACGGTAAAAAGTTCGACAGCTCACGCGACCGCAATAAACCTTTCAAGTTTATGCTTGGCAAGCAGGAGGTTATTCGTGGATGGGAGGAGGGTGTAGCACAGATGTCTGTTGGACAGCGTGCTAAGTTGACAATTTCACCTGACTATGCTTATGGCGCTACGGGCCATCCCGGGATCATTCCGCCACATGCGACTCTGGTATTCGACGTTGAATTATTAAAGTTAGAGACAgctagaggggccgctgcaggtgctggtggagctggaagaCGTGGAGTACAAGTAGAGACTATCTCTCCAGGTGACGGTCGCACTTTCCCAAAGCGTGGCCAAACCTGTGTTGTACATTACACTGGTATGCTGGAGGATGGGAAAAAGTTCGATTCCAGTCGCGACCGTAACAAACCGTTCAAATTCATGTTGGGAAAGCAGGAAGTGATCCGCGGGTGGGAGGAAGGCGTGGCGCAAATGAGCGTCGGTCAGCGGGCTAAATTGACCATTTCCCCTGACTACGCGTATGGGGCTACTGGGCACCCAGGGATTATTCCGCCTCACGCTACACTTGTGTTTGATGTCGAACTTTTGAAACTGGAAACTGTCGACGGAGAAGGAAGAGGAAGTTTATTAACATGTGGAGATGTAGAAGAAAATCCAGGACCAATGATTGAACAAGATGGATTGCACGCAGGTTCTCCGGCCGCTTGGGTGGAGAGGCTATTCGGCTATGACTGGGCACAACAGACAATCGGCTGCTCTGATGCCGCCGTGTTCCGGCTGTCAGCGCAGGGGCGCCCGGTTCTTTTTGTCAAGACCGACCTGTCCGGTGCCCTGAATGAACTGCAGGACGAGGCAGCGCGGCTATCGTGGCTGGCCACGACGGGCGTTCCTTGCGCAGCTGTGCTCGACGTTGTCACTGAAGCGGGAAGGGACTGGCTGCTATTGGGCGAAGTGCCGGGGCAGGATCTCCTGTCATCTCACCTTGCTCCTGCCGAGAAAGTATCCATCATGGCTGATGCAATGCGGCGGCTGCATACGCTTGATCCGGCTACCTGCCCATTCGACCACCAAGCGAAACATCGCATCGAGCGAGCACGTACTCGGATGGAAGCCGGTCTTGTCGATCAGGATGATCTGGACGAAGAGCATCAGGGGCTCGCGCCAGCCGAACTGTTCGCCAGGCTCAAGGCGCGCATGCCCGACGGCGAGGATCTCGTCGTGACCCATGGCGATGCCTGCTTGCCGAATATCATGGTGGAAAATGGCCGCTTTTCTGGATTCATCGACTGTGGCCGGCTGGGTGTGGCGGACCGCTATCAGGACATAGCGTTGGCTACCCGTGATATTGCTGAAGAGCTTGGCGGCGAATGGGCTGACCGCTTCCTCGTGCTTTACGGTATCGCCGCTCCCGATTCGCAGCGCATCGCCTTCTATCGCCTTCTTGACGAGTTCTTCTAACTCGAG

**Inserts:**

**>HIC1 homology region**

ATCAAAACCAATGCAAATAGCCTTTTTAAATTTGTTGATAAAAATGAAAGTAATGAAGAAGGTATTATAAATATAAATACATATGACAAAGAAAAAATCCATCATGAGATAAATGAAAATATGCAAGAGCAAAAATTAATTATACCAAATCATGACCATTTGTTAAATAATATACAAGATAAAAATGGTGGCAACATTATTTATAATAATAATAATATTTCAAAACGACAGCAGCAAAAAGGAAAAGAACATAAAGAAGATGAACATAAAAAGGAACAACATAAAAAGGAAGAACATAAAGAAGATGAACATAAAAAGGAAGAACATAAAGAAAATGAACATAAAAAGGAAGAGCATAAAGAAGAAGAACATAAAGAAGAAGAACATAAAGAAGAAGAACATAAAGAAGAAGAACATAAAGAAGATGAACATCAACATGTACAACATCAACATGTACAACATCAACATGTACAACATCAACATGTACAACATCAACATAAACAACATCATCATAAACTTGTTAATAATAAGGATGACCAAATTAATAATGATATCCAAAAATTATACAATGATGTTCATAATATGTGCATTAAAACAAAATATGTTAATAAAGATATTATTACTAAAGGACTTGGTATATATCCTTTACTAGCGAAACCATTACTACATAGACTTATTAAAGAAAATATAATTAAAGAAAAATTTATAAAAAATAAAGGATATGAAAGTAATATATATATACCCATGCAAAATACATTTAACAAAAAGAATAATAAAGACGTCCTAAGGTGTGATACTATGTCCACTACAAATTCGACAAACGACGAAGAGGATAATGAAAAAAAAAATAAAGTAAGTAAA

**>HIC2 homology region**

AACCATATCGGATACTTGGGAAGTAAACATTTAAAAAGAGAACAATATGAAGATAGTTCTATTATACAAAAAGATGATAATAATGATAATAATAATAATAATAATAATAATAAAAATGCCTATGGAGATATAAAAGGTAAAAATACATTTATAAAAGAGACCGTTAATGAAAATCCACAATTACTACACGATTGTTGTGATTTAGAAGATAACATAAAATATAAAATGCTCAATCAAAAGAGTAAAGATCGAATTGTAAAAATAAAAAGGAAAATAAAGAAATTTATAAAACTCACCAAAGGAGAATTAAAACTCTTGTCCAAAGTATTACATCATAAATTCTTACTTAATGAAATTAGTGCTATAGGTAATTATGAACAAAAAACTCAAAACAATTTGGAAATATTAGAATTAGAAGAATTTAAATTAAGAAAGGATTTCCTAAATATGTATTGTGATTGTCTATCTATTGAAATAACAAATCATGAAAAGGAATTCTTAAAAGAGTTTCAATTATTTATGCCAGACGTAAATATTATGGCAGACAAAAATATTTTATATGTACTTAGGCATTCAAAGAATGAAAAAAATTGTGTGTTTAAATATTTCTTGGATGTAAAAGAGACAAGTCATAAAAATGGTAATATCGATCATCTA

**>HIC3 homology region**

GATAATTTGATTCCTTTCGACGTTTTTAATATAAATGAAGCTCATTTATCGTTAGACCAAACGCAACAAATAATTCCCTCCGATACAATTAATAACACAGAAAATAATAATACTAGTACAAATAATAATAATAATAATAATAAAAACTTTTATTTAAAATTTGAAGAGAGGTTTAATATGAATAATGATAGAATAATTACAAATAAATTACCAGATGAAAATAATTTCCAAGCTTTAAATGAACTGAATAAAGAACTTAGAAACGCACACAATCTTAGTATAATAAATTATGATAATAATAATAATAATAACAACAACAATAATAACAACAACAATAATAATAATAATAAGTTAAATAAGAAGAAAAAGAAGAAGAAAAATTCAATTACTACAAATATAAATAAAAAGAATAACAACAAGATTAATCGTAGTAATAATATTAATAATTCCTATTCAAGTTTTAGAAGAATTGATAGTATATGTAATAATTTTAACTTCTCTATGATAAAGTATAGTAATAAACATAATAATAATAATAACAATAATAGTAATAATATTAATAGTAATAATATTAATAGTAATAATATTAATAATAATAACAATAAGAGAAAAAATACAAATAATAAAACTATTTATCAATATATTAATCCTAACAATACAAATCAACATTTTATCAAACGATTAAGGAAAAATAAACGAATTAGTAAAAAAGACAAAAACCTAAAAGAAAATCAAAAAAAGAAAATTAAACGAAAA A

**>HIC4 homology region**

AACAGGACAATTGATAACGAACCTGTTGTGAAAGAATCTATTGTAAACGAATCTGTTGTGAAAGAATATGTTGTGAAAGAATATGTTGTGAACGAATCCATTACGAACGAATCCATTACGAACGAATCTATCACGAACGAATCTATTACGAACGAATCTATTATGAACGAATCAATTACGAGTGAACAAAGTAATAAGTCCATTGAAAAAAATTCATTTGAAAAAAGCTATTCTGTTGAAGTGGTTTCATTAAACGATAGTACCGATATAAATGAAAACCCTAGAAATTTAGAAAATAAAAATGCTCCATTAAAAAATGTAAAAATGTTTGAAAAGAAGGTGTCAGAAACACCCGCAATGGAATTAAGGCTTTCAAAAAAACATAGTCTTTTAAAAAAGAGAGAGAACTTAGAAATGCTACATCAGAAATGTAATGAAATTGATGAAAATAAATGTGCATCAGGATTTGTAAATTATATATTACAAAATATTCAGAAAAGAGATGGTAAAGAAACATTTGATGATTTATTAAATTATATATTAAAATTAAAAGAAATATATAATATGTCAACAATGGATTTTATAGGCGCCTTCTTGATATTAGAGTCATTCAATTTAAATATTATAAATGAATATCCATTACGTGAATGGATACTAGTTACATTTCATTATATAAAAGGAATGATCACCTTACAGAATATTAAATTTATAATTAAATCATTAGATTTAAATCATTTAATTATTAGTAATATAACAGCTGCATTCTATATGAATATTAAACCTATTAAAGTAGATGAAGAAAATATAGAACGTGTACTTAATCTCTTATCAGAAGTTGTCATAAGAAGATCTTCTCGATTTAAACTTCTTAAAACAGATACTGAAAAGCAAAATCAGCAGGATGAAAAATTAGGATCCTGCTGGCAACCAGTTCACACAATAAAAGATTCA

**>HIC5 homology region**

CAGCACAATGATATAAACAACAATCAAAATTATACAAATCAAAATAACAATAATATTACTAATCAAAATCAATCATGTCTTTTATTGTGTCAGCTACATTGTACTATTGTATCACGTCAACACGAAATAGCTGATAGTTTATATAAAAAATTAAATTTCGGAAATAAAAATGTTACTCTACCAAATAATATAAAAAAAAATACAAATATTTATTTGTTCAATCTCTATTTTGATGAAAATATGAAAACATTATTATACATTATACCATCCAAAAAgtaaaataatttaaaaaaaataaaaatatatataaaaacagatatatttttcttcctatatattaatacatatattcatattaaattttttttttttttcatttttttagAATTTATAAATCCTTAAATAGTAATTGGAAGGTTGGTTTAAAATGTAGGATACAAAAACCACTATGCGCTACTAACTATAATGCATATTCAGCTAGAATCAGGAGGgtaaaaataaaaaaaattataaaaataatgaaaatataaacacatatatgttttaatatgaatatataatgtattgttacaaattttttttttttttttcacgttatatatatatatatatacacaaatttatatatacatatatatattattttttatagATTATTTACGATCCTCAACATTTATGGAAATGTGTAACGGTGAAAAAAGACACTCAGAAATTTAATAAAAACTCAATAAATCAAAAAGTTAACTTTTGGGAATTAACTAAAAAGAAAAACGTG

**>HIC6 homology region**

GATATGTTAAATAAATCAAAAATGGAGGATTTGTTAATTATATTTAAGCTTATAGAAAATTGTAATACGAAATATATTAATGTTTGTAGTATGTTTGTTGAATATATCAATATAAAATATAATGCAAGTAATAAATATATGGATGATCAAAATTATAATGAAGACAACAAATGTATAAAAGATAAGAAGATAAAAAATGGTAGTAATAATTTAAATGATCGTAATAAAAATAAGGGTACTAACCATTACATAATTAAAAATGAAATAAATAATGAAGGAGAATCATTTCAAGAATTATTTTATAAATTCATTATAGCATTATTATCTCTTTTCTATTTTATAAAATTAATACATATACCATCCTCTTTTCTTAAAGGCAAAGAAGATCCAGAAGACTCATTTAACGAAAATCAAGAACATAATAATGTAAACATTTATTTTGATCAAGAAAATAATACTTATAATGATGATATAAATGAAAAATATTTTCCTCAAGAAATTATACATAGCAAAAATTTAAAAAAATCAAAAGATGTGGAAGATATATCAACAAATAAGGGAGAAATAGAGATGCTTTTATCAAACAATATTGAAAAGTCTTTACCACAAAAATATCATGATAATACTAATTTAACAAATGATTTAACATATAAAAAATATATACTAGATATATTAAGTAACATAAATATAAAAAAATTAATATTTGGAAAGGCATACATG

**>HIC7 homology region**

TGGTAACAAAACCATTCTTTCTTCTTTTCTAAATGTTTTCCAAAATATGCAATTATTAAAATGTAATAATTATCTTAATTGTGAAGATATTAAAATATTAACTAACAAACTTGTATATAATTCTGTATATACAGATATAAATAAGAACAAGAATTTAACACGAATAGACGAAAATAAGAAACAAGCAGAAAATAATAATCCTATGGATAATGTATATATTAATAACATTTTTGATAGAATTAAAAATGACTTATATAAAGAAAACTTTATTACATATGGAAATATAAATAATACTCAAGAAGAAGATGACAAAATTCATAATCCCGATGATAATCATCATCATATGGAAAAAACTAAAGAAAATAATATATTTAAAAATAATTTGTATTATATCAAAATGAATAATATGAAGAATATATGTTTATATGATGATTATGATATTTTCAATAATTTAAAAACGAAAAATGAAGCTCTCAATTGTGTAAATGCTATTGTAACAGATATGAATTCTTCAGCCAAAAAAAATGACAACGCAAATTTTAAGAAAAGAAAAACAATCGACTCGTCTTTGAATGAGGTTAGAAATTCGATACGTGAAAGGGACTTAAGAAGGCTTAACAGAGAAAAGAGTAAAATCTTA

**>HIC8 homology region**

GAGAAATCCTATAATGAAGAGGGAACTAGCCAAACGGATAATAGTGATAAAGAAACTGAACAAATGGAAAATAGCGAAGCGGAAACTAGCGAAAATGACAAAGATGAAAATAAAACTAGCGAAAATGACATAAATGAAAATAAAACTAGCGAAAATGACATAAATGAAAATAAAACTAGCGAAAATGACATAAATGAAAATAAAACTAGCGAAAATGACATAAATGAAAATAAAACTAGCGAAAATAATAAAAATAAAGAAGATTCTATTCAAAATGATAAAAATAAAGAAGAATCTATTCAAAATGATAAAAATAAAGAAGAAGCTATCCAAAATGATAAAAATAAAGAAGAAGCTATCCAAAATGATAAAAACAAGGATTTAAAGGAGTATATGGAAAATAACGAACAAAGTATGAATCAAAGTGATGACAATGCATTGACTAATACAAGCTATGAAATAAAAAAAAATGATGATGTAATAAATAAAGAGGAGGATGATATTATGAATGGTGAAATACCTAATAGTAATTATATAAATGAATTACATATTATACTAAAAGAATTAAAAATAGAAGAGAACAAAATGTTAGAAAATATAATTGAAAGATATAAAATAATATATGGTATGAATAAAGAAGTAGATTATAATATGTTAGAAGAAGATGTGAGTAATTTTCATAATAGTATAACACATGAAGAAAATATGTTGAAAAATAAAAAACTAGCCCTTTTGAGAAAAAAAAGAAATATATTAAAAATGTATTATTCTTATTCGTCTGATAATAGTGAGGATGACGAA

**>HIC9 homology region**

GCTATTGATAAGGATGATAGGAAAACATTATATTATTTATATAATGTAAATTATTGTTTTAATGATCAAAATAATAATAATAATAATAATAATAATAATATGAATAATTCAAATATATTTGGTAATCCTCATAATCCAGAATTGGTAGTTTTAAGTTATAAAAATTATTGTTTTTATATTGTATGGATTTCTAATTTATTATTAAATCATAAAATAGAATATGAATCATTAATATATGTATATTTAAAAATATATAATAATACAAAACATCAAAAAGCATCATTATTAGAAGAACATGAAATTACCATTATTTTATATTACTTATTTACTATGTGGATTAATGGAGATAAAAATAATTCTTCCTTTTTCTTTTATAATGAAGAAAAAAGTTATCATGAAAAAAATAATCTTGGATATTTTTTAAAAGATACTTATGGAAATTTACAATATATAAATAATTATTCTCTAATTACATTATTAAAAGAATTATTATCAAGAGCAGAATTCTATTTTGAATATACATCAGATGCTGCATCAAGAAATTATGATATTAGTTGTATAATTCTTGATTCTTCTATATATGATAATGAAAAAGTTAAAAAATCGTCAGTAGATATTCTAAAAAAATTCTTTGATAATATATACATAACCTTTAATGATGTTCTTATTAAAATACCATTCTTAAATCAAATTTTGGAATTCCAAAAAATACACGAATTTTTAAAGAGGTTTAAAATATATCTCGATGAAATTATTTGTCGCATAACCTTAACCGAGGGAAACTATATC

**>EUC1 homology region**

GGTAAAAATCCTTTAATCCATGAGCTTCTTTTGTATCTCATAAATAACACTAGGTGTAAAATTATAATgtaagaaaaaaaaaaaaaaaaaaaaaaaaaaaaaaaaaaaatataaatataaataaatatatacatatatatatatatatatatatatatatatatatatatatatatatatatattgtggataccctatgaatgccatgcaatatgtaatatataatatgctttatatttatatttatatttatattttcattattttatttttagGTTGGACAACTGTAATTTGGAAAACCCTTTTTTAATAAACGTGAACAAAAATATTGATCAATTAAATGAGAATAATTATTTGACCTTATTATCTTTGAGATATAATTATTTTAGTAGTGTTAAGGAATTAATAAAATTTGTAAACAATTTAATATCGAAATGTAAATCCTTAAAAACGATTCGTATTTATACTAGTAACTTTAGTGAAGACGATATGAAACTTATAGAAAAAAATATAATAAAAAAAGATGTATTATCATTTATAAGTACTTATGCTTTAATACCTAGTAGTAAAAAATTAAAAAGAAAAGTCCTAAAATTAAATAAATATAAAATTAATAAGGATGTCGAATATAATACTGAACTAACTGAACGTGAAAAGAAACTTATTGAACAATTTTTTGATAGACAAAATACAAAAACTGATAAAGAAGAAAAAAAAGATGATTTGAACACATTAAATAGATATAATTTAAATGATATAAAAAATAGGAATATTAAATATAAATTTAAAAACAATAAGATA

**>EUC2 homology region**

CCCCTTCCATAGGAAAAATAAATGGTGATATAAATCATAATTATGTTGACTCACATTTAAATAATATAAACGTAAAAAATAGAGCAACCATAGTTAATGAAGACGAACATTTATATACAAATAATAATTATATAGAACTAGTTAAAAATGATGATATTATATTAAATGGTAATTATGTAAATTCATCTTCTGATGAACAACAAACAAATACATTTGATGTGAATTTACATGATAATAATAATAATAATAAGTTAATAGAAAATATATTAAAACCGTATGTTAAAGAGGAAGAAGAAGAACTTGAATATTCCAAAAACCTTCGAAAAATTGCAACAAAATGGACAGATGAGGAAAAAGATATTTATTTTAATATTTTTGGAAAGGAAGGAAAAAATTGGGATGCTCTTTATGAAGCTTTAGAGCCATTTGGGAAAACAAAAGATCAAATAAAAAATTTTTATCAAAACACAATAGCAAAGAGGAAAAAAAGCCATATAATTCCA

**>EUC3 homology region**

GGTAAGTGCATCATCTATTGGTATGATAAATCCTGGTCATATGAATAGTTCGCATAGTGGTAGTATAATGAATAAGGATAAATTTAGAGACCTCCAATTGAATAGTAATAACAACAACAATAATAATAATAATAATAATTATGTAAAAGTTAATCCTTTTGATCATCCCATGAAAAATATGAATCCAAATTTAATGGATGATAAATATCCCAAAGATAATAATAATATGATGATGATGATGATGAACAGTGTTTACAATATGATGGGATCAGATCATTTCCATATGAATAATAGTCATATGATGGATCCGATGGGTTCACCTAAAATGAATGATAATATGTATAACCATAATAATATGAATTTTGAAAACACCAACACTAATCACAATAATCACAATAATCACAATAATCATAATAATCTTAATAATCTTAATAATCTTAATAATCATAATAATCATAATAATAGAATACAGGATAAGCTTCCTGAA

**>EUC4 homology region**

CCCCCAATAATTCCAGATAATAAAGAGAAACAAATTAATAATGATAAGAACACAATTTTGTCAAATCGAAATTTAAATGATAATACAAATGATTATCCAAATGATTATCCAAATGATTATCCAAATGATAATACAAAAGATGAAGAAAAAAATGGACTTATCTTAAATCCAATGAATTTGGAGAATTTAGACCAATGTGAAGATTTAAATATACAATATCTAAGTCAAATAGATGATATAAATTTTGATAATTTAAATAAAAATAAATCTAATAATAATAATAATAATAATATATTAAATTCTAATAATGAAAATATGTATGATGATAATAAAAATAAGTCAAATGTATTAAATCAAGATATTAAGAAAAACAAAATTTTAGCTATTCCAAATTATGAAGAGAAACAATATTATATTGTCACCAATGAAAATGAGCTAAACATTTATAGTAATTTTAATGATCAAAAAGATAACATAAAAGAATTAAAAAATCAAAAACAAGAAAAACAAAAAATTTGTGCTATAACCAATTTAGAAGGTAAATATTTTGACCCCCTAACACAACAATATTACAATAATGCAGAGGCATTTAAAAAATTACGTTTATATTATCATCAAATATCATACAATAATTTAAATAAGGATATATGTCTAACAGTCAAACATTTCAAAAACAAATTAGCCGAAATAGAACAATACGTAAATAGTAATAATAATCATAATAGTTGTAATGATATACCTTGTAATAATAATGTGAAACAT

**>EUC5 homology region**

GCTGTTAAGGTTAATATTTCTGCATATAAAGGATTTAAACAAATAAAATGGACTAAATGTTATGACATGTTAATATCTTTATCAAATGTAGGAAATAACATATTAGTATTAAAAAATAAACATTTAAATAACTGGAGTTTTTTTATTTCTGATTTTGAATTAATAGATTCGAATATTGAAGTTATTGAAGATGATAACGAATTTGATATAATAGAAAATATCCAACCGAAACATAAAAATATGGAAGATAATATATGGAAATATATAATTATATATTTAAATTTATTTATTAAAAATAATGTATTTATTCAAAATTTAGTGCATCCATCTTCTCTTTTTCCTATTTTATATACAGGTTATTATATTAATAGTCCTAAGTATTTTTTTTATAACTTATCTTATGATTTAACAAAACAAAACTTTTTAACCTATCAATCAGATGATGATGATAAATGTAGTAAACCAAAATATGTAAAGGATAGAAATAATTGTAATGAATTTGTTAATATAAAGGAACATGTGGAATCATATATGAATACAGACCAAGAAATATGTACAAATATAAATGGAGAGATGAATCATATGGAAAAGGCAAATTTACAAATTAAAGACAATTCAAATAATATTGAATTAATAACAACCTCTATGAAATATTATTTGTATTACAAAAATATTATTAACTTTTCAATA

**>EUC6 homology region**

GTGAATCTAGTCCTAGTTCATCAAGTAGTAGTATGCATGATATAACATATCATAAAAACTTATTTGAGGATGATAAAATAAGCATAGTTAAATATTTTATGAATTATAGTGATGATGATAATAAGAATACATATCAAGATTATTTAGTACAAGAAAATAATATAAATAATAATGTACATAATAATGGAGTAGTGAGAGATATTAGTGAGATAGGAAATGATAAAAATAAGAAAAATGAAAATACATATATGAACGAATTAAATAATTTGAAATATATAAAAAATGATATATATTCTACAAAAAATTATATAAACAAAACAAATCAATTATTAAAATTATATTCGAATGATAAACTGAATGTTCTACAATATTATGATGAAAAATTTAAAGAAAGAAAAAACAATGAAAACGATGGTAGTGAATATGCTCAAAAATCTTCTATCGATGAAGTTCATAATATTCATTATGAATCTAAAATGTATAATGTGAACTTTTGTATTAAAAAATATTTACGTAATATGAAATATGTAAACTTGAAAAGATTAAAGACAATAAATGAAAAAAATTGTAAGCTATATGTAAAGGGTTGTACCCACTTTTTAAATTTAAACAAATTTAATGATGCCCTGTCTTGTCTTAGTAAACTACAAAACCTGAAAGGGAAAAAGAAAAAAAAAAAGGAAAAGCAAAAAAAAACG

**>EUC7 homology region**

CGTACATCCATTACCAATGATCACGATGCAAATAATACTGATATTAAACTCAATTATTTTATAAATAAAGGGAATAAAATAAATATGAATGAACCTGATTATAATACTAACTTATTATTTCATAAAAATAACACTGAAGAAAATATGTTAAATCAATATATACAGAATAAGAACAATTTATATAATATAAACAATATAAATAATATAAATAGTAACATAAAAATTAATGAAGACAATACAAAAATGGTATCGCATAATATACTATCCTTGGAACATCAAAATTTAAAAAATCAAGATTTCTATGAAAAATATGTTAGCACAAATATAACGAACAATTATAATAATAATAATAATAATAATATTCCTTACCCTTTGGTTAGTAGGAATGATAAATATGCTCCGTGTATTTTTTCAAAAGAACAAAATAATAAAACAATATTTCATTTTGAAGATATGATTGACCCCTTAATGAATAAAGACAATTATTTATTTTGCTTGAGTAAGTATGTGTACGTACAAAAAAAAATATTATTTTATTTGGAACACTTATTAAATTTCGATAATTACAATTTTAAGAGTAAAAAAAAAGAGCTACCCAACAATAAGGATATGAACATTAATTTAAAAATTAACACTAACGATAATATAAATAATGATAGTCTAAATAATGATAATATAAATAAGGACAATATAAATTGTGATGATATTAATAATAAAAAAAATGATGATCTTATTAATAATACATTATCTGAACTTAACGGAACATGTCTTAAAAGTAAAGTTTCACATAAAAGCTTAAAACATAAAACAGCAAAGGATAACAATATTAAG

**>CIC1 homology region**

TGCATCAACAAAATTCATTGGGCAGTATTTTAAACAAAACTAATAGTTTTAATAAAAATGATAATAATAGTACACAAGAAGAAATTAAAGTAAAAAGAAGAAGAGGTAGACCTAAAGCAAGTGAAGTAGCTGCTATTAATATTATGAATAATAAATTAAAACAAAATGATATAAGAAAATATTATAGCAAAAATAGTGATAATGGATTTATGAAAAAAAATAGTATGAACGAAAAAAATAAAGAACAACAAAATGAAGAAGCAAATACTAATGTTTTTAAAATTATTAAAAGTTCAATACAAGAAAATTCTTCTTTTATGAAAAAATCACCAAACGAAATTGTAGAATTAGAAAAAAATTACAAAAATAATATAAAAAAAGGAGTGACATGTATACCTTTAAATTATCAAAGCAAAGGTGGTGCAATGGCTATTATATTAATAGGAACAGATACTACATATGGGCCTGTAAAAAATTCCTATGGATTTATGACTTTTCTTGTATTAGATTGTCATACAAATAATTTTTTTATTGATACAGGAATTAAAAATAACGTTATAGAATGTGAAAAACATATGCAACTTTTAATAAGTCCAGGTGATATGTACATGTTTAAAAATCAAAGTCACGAAATGGAAGCAAGATTATTATTAATTGTTTGTAATAAAATGAATCAACCGTTTGACGTAAAAATGCATATAAAAGACCCACAAATGAACAACGAAAAA

**>CIC2 homology region**

ATGTGGGTATGATGAGAAGTCGTGTGATATATATGAAACCTAATGATATAAATAAATTAATATACTCCTTTGAAAGATTAAATGAAAATATATTCGATTGTGCTGATTATATATATAAACATATTTTTAAAAATAAGAAAGAAAAAATAAATGACGAAAAAAAATTCTATAAAAATATTTATTTATGGTATTTGAATTGTATTCATATTATTAATTTACAAATATATGTAATGTGTCACTTTTATAATAATTATCATAATACTACAATTCATATAAAAAATAAGAAGAAAAATGTAAACATAAATAATATGAACAATAATAATAATAATAATAATAATAATAATATTCTATACCATTCTTTACAAATCAATATGAATATAATAAAAGGAATTAATAAGATAAAACATAATGTTGAACGATATCTAAAAGAAACTAAGCAATATTATGAAATTATAAAAAATAAAGAAGAAATAAAAGAAGGAAAAAAAAAAAAAGACAATAAAAATGAAACAAAATATTCTCTTGTACAAAATAATCAAACGGATGCAAATATATTCAATATAACTAATGAGGAATGTATCGAAGATTATGAAGAATTAAATTCATTTTTAGATTTTAACTTAGATAAACAATCAGGTATATTGTTGGAAGGTACTACAACTAATACAAATAATAATTTATTAGATCAAAAGGTTAATGAAAATGTTTATAAAATGTCTAATTATCTGGTGGATGAGGACTGGTTAATGTATATACGGTCTCTAATC

**>CIC3 homology region**

GATATAAAGTGGGATATGGATTTGGAATATTCACTTAACCAAATTAGAGAAGACTGGAAAAATGAAAACAAACAAAATAAAAAAATTATACAAAAAAAGgtaatacaaatgataagaaatatatatatatatatatatataaatttacatatgtgatatattaatatattaatatatgaataattttttttgtagATTATTGAGGGTATTAAACACCGAGTTGGGTTATTAGATTATCTGAACAAAGTAGAAATAAACTTGAACGCCTTTGCAAAAATTATTGAAGGGAGAATGATATCAAATGATGACGTAAAAAGTCAATTATTATCCATACAAGATATGAGTGAAGAAGGGTTACTTTCGAAGCTTTTAGAAAATTTGAAATGTAATAAAATTGATATAAATGCTGAATCTTTTAAAACACCAACTTCTTTTTTTGTTTCTCATCACTTACCTATGACAGTATGTTCATATAGTGACCATTCAAATAATCATGAATTAAAGGATATGAATGAAACTAATAGTAATGCTGATATTAAAGATGATGAAAAAAAAGTGACTGATGAAAATGATGCCATAGATACTCAAAATAAAAATGAACACAACGAC

**>CIC4 homology region**

GAAAAAAGTACGCACAATTTTGGGAAATGTTTTTTCATATTTTATTAAATTTTTGTAATATATATGAATTAAATCATATACGATATAATTCAAATGAAGAGGTTCTGAATAATAACTCAGATGATGATTTGTATGTTCATAAATTAATAAATAAAACAGAGAATTATTATTTTGATTTTAAGAAAAATAATTGGCAAGAAATTCATAAAAGTCCTCAAAGAAATGATTCAAATATTTCTAACGATAGTAATAAAAATATAATTAATTTATCGGATAGATGTTATAGTAACATACCATATGAAAATAAGTCAACTAATTTATATTATATTGATGGAGTGGAAAATCACAATTCCAAAAATATAATAAATAATCATCAAGGTCATAAAGATATAATATTGTCAACAAATAAAAATATCCCAACAAAACATCCCAATGAACATATTAAATCATATGAAAACAATGTATTAATAAATAATGATAATAATAAATTATCGATTTCTAAAAATGGAAATTATAAAAATACAATACTAAGAAAAGATATACAAGGACATAACTACAATAGCATGGGAAAAAACAGCAAATATTTTTTTATAAAAAATTCCATATCAAATTTTAAATGTATAAGGAAAAGCTTGTATAAACATAAATATGAAAATAAAACTAAGGATCAAAATAAAGATGAAATGAAAAATAAAAATATACATCCCCCTGAGGAAAATACACAAAGTGGAAAATTGAAAAGAAAAATCATATTTTTTGAAAACGAACAAAATAAAACCAAATGTAGAAAATTAAATAATGAAAAACAATATTATAATAATGAATATAAATATAACGAAGCTAAGAGATCTACAAGATATATTAATAAACTCATGAAAAAAAAAGCAATCTTTATCCTTCTAAATTTAAAAAGGTCTATTGAGCGCATATTTGATGAAGACGAGGAGAAGCAAACAATCTTGTTAAACGAACTTTATAACGCCTCCACTTTTTTT

**>HP1-mCherry_Lyn-FRB**

XXXX = HP1

XXXX = Linker

XXXX = mCherry

XXXX = Lyn

XXXX = FRB

XXXX = T2A skip peptide

XXXX = yDHODH

CGGCCGCTAACGTAACAGACTTAGGAGGAGATCTtaaAGTCTCCTTTTCTTTATTTTACAGTTGTGAATTATATTCATCTCATTTATTCTTTCTTTTTTAATAATTCAAGGTAAACGTTTTTATATTTTATCATATTATTATGGTTCATAACGTGTTGTATATATATATATATATATATATTATATATATTTATTTTTTTTTTAATAATAAAATTTTAAATTATAAAAATAAAAAAAACAAAAAAATATAATAATTTAATAAAATATAAACCTAAATATGTAATTAATTATTCTCTTTATTTTAAATTAAAAAGGAAAAAATATATATAATAAATAATGGTAAGGAAGAATCCTTATAAAATAAATATACATATAATTTTATTGAGTTAATATATAATAAGTATTATAAAAATTATATATATGTATATATATAAATATATGTTATATATATATATTTATTATATATACATAATATTATAGAATATTGACGGAAAAATATTAAAAAAAAAAAAAAAAAAAATTATATAAAATATAATTCTAAAAATTTCAGGATCCCTATTTATAAAAAAAACAAATTAACTTCAAATTCTTTTTCTTTTTTTTCATGTATATATATATATATATATATATATATATATATAATTTTTCAAAATTTTTTTAAGATATAATAAATATATTATGTATATTTTATCTATAACATTTATATTTTAAATTGAAAAAAAATAAAATAATACATAAAAGATAAATATAATGAAGTAACAATATTATTATATAATTAAATAATTGAATGAATATATATATATATATATATATATGTTAAAATTGATGAATGAATTATAATGAAAGTTAACTAAATGTAAAATATAAGAATAATATATATATTATCGAAGAAGCAATAAAATTTTTATGATATTTAGATTTCAAGTAAAACATGTATGAATAATATACATATATATATATATATATATATATATATATATATATATATATATGTACATATATACATATACATAATATATAAAATATAATTTTAAAAGTAActcgagATGACAGGGTCAGATGAAGAATTTGAAATTGGTGATATACTTGAAATAAAAAAAAAGAAGAATGGTTTTATTTATTTAGTAAAATGGAAAGGATATTCAGATGATGAGAATACTTGGGAACCCGAAAGTAATTTAATACATTTGACAACATTTAAGAAAAAGATGGAAAGCTTAAAAACGAATTTTTTATCGAAAGCTAATGAGACAAATGGTGATGGGAAAATTTTGAAAAATCATATATTAGCACCAACACAAGAAGACGATAGTATTAAATCAAAAGGTAGAAGTTCCTTAGCACCCCGACGAAAAATGAGTAGAAAAAGTTTAACGAACAAACTAGAAAATAAAAAGAACTTATCTTTATCAGACAATTCTTTAAAAAAAAGTGATGAAGAAGATAATGAATCTGTAAAACATGAGAATCACGTTAATGATGGAAATTTATTAAATGTTGAAGATGTATATAGCGTTCGTATTAAAAATAAGAAATTGGAGTTTTTGGCTAGCTTGAAAAATGAATCTCCACAATGGGTAGAAGAAACAAATATTAGAAGAACTGGACATTTAAATATTAAAGTCAATGATTTTAAAAGATATGTAAGAAGAAAAAAAAGTTCTAGGGGTAATAGAATAGTTATCAAAAATCTACACAACGTTGGAGATGAATTATATATTTCGGTTATTCATAATATAAATAATAAAGAAATTCATAGTTTATATCCTTCCAAAGTTATTGAATATATTTATCCACAGGAACTCTTAAATTTTTTATTATCAAGACTAAGATACCGTACAGCTCCTAGGacaagttatccatatgataatccagattatgcAccagttgcaacattaggtaccatggtgagcaagggcgaggaggataacatggccatcatCaaggagttcatgcgcttcaaggtgcacatggagggctccgtgaacggccacgagttcgaGatcgagggcgagggcgagggccgcccctacgagggcacccagaccgccaagctgaaggtGaccaagggtggccccctgcccttcgcctgggacatcctgtcccctcagttcatgtacggCtccaaggcctacgtgaagcaccccgccgacatccccgactacttgaagctgtccttcccCgagggcttcaagtgggagcgcgtgatgaacttcgaggacggcggcgtggtgaccgtgacCcaggactcctccctgcaggacggcgagttcatctacaaggtgaagctgcgcggcaccaaCttcccctccgacggccccgtaatgcagaagaagaccatgggctgggaggcctcctccgaGcggatgtaccccgaggacggcgccctgaagggcgagatcaagcagaggctgaagctgaaGgacggcggccactacgacgctgaggtcaagaccacctacaaggccaagaagcccgtgcaGctgcccggcgcctacaacgtcaacatcaagttggacatcacctcccacaacgaggactaCaccatcgtggaacagtacgaacgcgccgagggccgccactccaccggcggcatggacgagctgtacaagCCCGGGTCGAGGGATATGGCAGCTTAATGTTCGTTTTTCTTATTTATATATTTATACCAATTGattgtatttataactgtaaaaatgtgtatgttgtgtgcatatttttttttgtgcatgcacatgcatgtaaatagctaaaattatgaacattttattttttgttcagaaaaaaaaaactttacacacataaaatggctagtatgaatagccatattttatataaattaaatcctatgaatttatgaccatattaaaaatttagatatttatggaacataatatgtttgaaacaataagacaaaattattattattattattatttttactgttataattatgtgtctccttcaatgattcataaatagttggacttgatttttaaaatgtttataatatgattagcatagttaaataaaaaaagttgaaaaattaaaaaaaaacatataaacacaaatgatggtttttccttcaatttcgatatcaatttatagaaacaaaatatatacttgtataattttatttttttatataaatcattacatatataattatacaatattttttctaagagataattatatattaatatatataaaaaaaggtgttttttttttttttttttatttttatttttattttatggtaatattttattttccttattttataaattatattagtttatatgtgattaattttatatattatcaatttatatatttttaaatgcttacttaattatctttttttttttttttttttttttttcccctctttttatattaatttatttttgaaaaaattgatatatatataGTTGAAATATAAATTTCAAAAAAAATGATCACAAAATATACACTTAAATATAGGTACAATAAAAAAAAAAAATAAAAATATAATTACAAGATAATATTTTTTCCTGCTATCAAATTTTTATATATTCTCCTCAAAGAAAAAATATAAATAAATGAAGTAAATTAAAAAAAAAATTTTCTTTTTCTTCTTCTTTTGTAATTCCTTATTTATACATATTTTACTATATTTTCATAAAAATAAATTGTCATATTATATAAATATATATACCAAACCATAATTATATAGCCCTCAACATATTTTTATGATGTTTTTTCTTTTTAAATGTGATACGTAATTAAATATAATAATATATATTAAATATTATATTTTGTAATATTTACTTTCATGAGGTTTAATAAATTATAAAGAGAGAATAAAAAAAAAAAAAAAAAAAATTTATCCATATACAAATTATTAATTTATTTTTATTTTTTATTTCCCTTTGTATATATTATAAAAAAATAATCTACATAATTTTATATGATGATAATTACAATATAATATTTATAATATATATTATTTGTTAAGAGAAAAAAAAAATAAATATATACCTTCTTTTTGAGAATTGAATAAATTGTTTAATATATATATATATATATATAAATATATTATACATTTATGGTGAAAAAAAATGTTGTATTTAATTAGATTTAATATATATATATAAAAAATAGCGTATTTAAAATAATATATATATATATATATTATTATTATTACAAAATGACGGATATTATAAAAGTATATATCTATATATATGTATATATATATAATATTATTTTACTATATATATATAATATATAAATGTAAATGCATATAGTATCTATGTATATTATATATATATATAATATTAATTACATATCTAAGTTTTCTTTTCTTTTCTTTTTTTTTTTTTTTTTTTTTATTTTTTATAGAGAGCCCGTTATATATTATATATTAAAATTTTTTATAATAGCATTTATATACAATATTTTATTAACTAAAAGAAAAAAAAAAAAAAAAAAAAAAAAAAAACGAGGAAATTTATATTTCTTTAACAACATTTTAATTAAAATCATGATAATACAATTTCAAATCATTTTGTAATTATATAAATAAATATATATATATATATATATATATATATGTACTTTTAAATTAGGAATATTCTCATTTATAAATATATCTTATTTTTTAAATTGGTATAAAAAAAAAAAAAAAAATAAGAAACCGTTGATTAAATAATACATATATAATATAAATATATTTTATAAATATATATTTATATATATATATATATATATTTATAACGTATATCATTTTAAAGATAAACTAGTATGGGATGTATAAAATCAAAAGGGAAAGACAGCGCGGGAGCAGATCCCACCAGAacgcgtgatccaacaagaagtgcaaatagtggagcaggagcaggagcaggagcaatattaagtagagctagcatggcttctagaatcctctggcatgagatGtggcatgaaggcctggaagaggcatctcgtttgtactttggggaaaggaacgtgaaaggCatgtttgaggtgctggagcccttgcatgctatgatggaacggggcccccagactctgaaGgaaacatcctttaatcaggcctatggtcgagatttaatggaggcccaagagtggtgcagGaagtacatgaaatcagggaatgtcaaggacctcctccaagcctgggacctctattatcaTgtgttccgacgaatctcaaagGGTGAGGGTCGTGGTTCACTTCTTACTTGCGGTGACGTTGAGGAGAACCCTGGTCCTGTCGACATGACAGCCAGTTTAACTACCAAGTTCTTGAACAATACCTATGAAAACCCATTTATGAATGCATCCGGTGTTCATTGCATGACTACACAAGAATTAGATGAATTAGCAAACTCTAAAGCTGGCGCATTCATTACAAAGAGTGCTACAACCTTAGAAAGAGAAGGTAACCCTGAACCACGTTACATTTCTGTCCCTCTAGGCAGTATCAACTCCATGGGTTTACCAAACGAAGGTATCGACTACTATTTGTCCTATGTATTAAACCGTCAAAAGAATTATCCTGATGCACCTGCTATTTTCTTCTCAGTTGCTGGTATGAGCATTGATGAAAATTTAAATTTGTTGAGGAAAATCCAAGATAGCGAATTCAACGGTATTACCGAGTTAAACTTGTCTTGTCCTAATGTGCCTGGGAAACCACAAGTTGCTTATGACTTTGACTTGACAAAGGAAACCTTGGAAAAGGTTTTTGCCTTTTTCAAAAAACCTCTTGGTGTCAAGTTGCCTCCTTATTTTGATTTTGCCCATTTTGATATCATGGCAAAAATATTGAACGAGTTCCCATTAGCTTATGTCAACTCTATCAATAGTATAGGAAATGGTCTTTTCATTGATGTGGAGAAGGAGAGTGTAGTAGTGAAGCCAAAGAATGGTTTCGGGGGTATTGGAGGTGAATATGTTAAGCCAACCGCGCTCGCCAATGTTCGTGCATTTTACACTCGTTTGAGACCTGAAATCAAAGTTATCGGTACAGGTGGAATTAAGTCCGGTAAGGATGCATTTGAACATCTTCTATGTGGTGCCTCTATGCTACAGATTGGTACAGAATTACAAAAAGAGGGCGTCAAGATTTTTGAACGTATCGAAAAAGAATTAAAAGACATAATGGAAGCTAAGGGTTATACATCCATAGATCAGTTCCGTGGGAAGTTGAACAGCATTTAAAAGCTTATTTAATAATAGATTAAAAATATTATAAAAATAAAAACATAAACACAGAAATTACAAAAAAAATACATATGAATTTTTTTTTTGTAATCTTCCTTATAAATATAGAATAATGAATCATATAAAACATATCATTATTCATTTATTTACATTTAAAATTATTGTTTCAGTATCTTTAATTTATTATGTATATATAAAAATAACTTACAATTTTATTAATAAACAATATATGTTTATTAATTCATGTTTTGTAATTTATGGGATAGCGATTTTTTTTACTGTCTGTATTTTTCTTTTTTAATTATGTTTTAATTGTATTTTATTTTTATTATTGTTCTTTTTATAGTATTATTTTAAAACAAAATGTATTTTCTAAGAACTTATAATAATAATAAATATAAATTTTAATAAAAATTATATTTATCTTTTACAATATGAACATAAAGTACAACATTAATATATAGCTTTTAATATTTTTATTCCTAATCATGTAAATCTTAAATTTTTCTTTTTAAACATATGTTAAATATTTATTTCTCATTATATATAAGAACATATTTATTAAATCTAGAATTCTATAGTGAGTCGTATTACAATTCACTGGCCGTCGTTTTACAACGTCGTGACTGGGAAAACCCTGGCGTTACCCAACTTAATCGCCTTGCAGCACATCCCCCTTTCGCCAGCTGGCGTAATAGCGAAGAGGCCCGCACCGATCGCCCTTCCCAACAGTTGCGCAGCCTGAATGGCGAATGGCGCCTGATGCGGTATTTTCTCCTTACGCATCTGTGCGGTATTTCACACCGCATATGGTGCACTCTCAGTACAATCTGCTCTGATGCCGCATAGTTAAGCCAGCCCCGACACCCGCCAACACCCGCTGACGCGCCCTGACGGGCTTGTCTGCTCCCGGCATCCGCTTACAGACAAGCTGTGACCGTCTCCGGGAGCTGCATGTGTCAGAGGTTTTCACCGTCATCACCGAAACGCGCGAGACGAAAGGGCCTCGTGATACGCCTATTTTTATAGGTTAATGTCATGATAATAATGGTTTCTTAGACGTCAGGTGGCACTTTTCGGGGAAATGTGCGCGGAACCCCTATTTGTTTATTTTTCTAAATACATTCAAATATGTATCCGCTCATGAGACAATAACCCTGATAAATGCTTCAATAATATTGAAAAAGGAAGAGTATGAGTATTCAACATTTCCGTGTCGCCCTTATTCCCTTTTTTGCGGCATTTTGCCTTCCTGTTTTTGCTCACCCAGAAACGCTGGTGAAAGTAAAAGATGCTGAAGATCAGTTGGGTGCACGAGTGGGTTACATCGAACTGGATCTCAACAGCGGTAAGATCCTTGAGAGTTTTCGCCCCGAAGAACGTTTTCCAATGATGAGCACTTTTAAAGTTCTGCTATGTGGCGCGGTATTATCCCGTATTGACGCCGGGCAAGAGCAACTCGGTCGCCGCATACACTATTCTCAGAATGACTTGGTTGAGTACTCACCAGTCACAGAAAAGCATCTTACGGATGGCATGACAGTAAGAGAATTATGCAGTGCTGCCATAACCATGAGTGATAACACTGCGGCCAACTTACTTCTGACAACGATCGGAGGACCGAAGGAGCTAACCGCTTTTTTGCACAACATGGGGGATCATGTAACTCGCCTTGATCGTTGGGAACCGGAGCTGAATGAAGCCATACCAAACGACGAGCGTGACACCACGATGCCTGTAGCAATGCCAACAACGTTGCGCAAACTATTAACTGGCGAACTACTTACTCTAGCTTCCCGGCAACAATTAATAGACTGGATGGAGGCGGATAAAGTTGCAGGACCACTTCTGCGCTCGGCCCTTCCGGCTGGCTGGTTTATTGCTGATAAATCTGGAGCCGGTGAGCGTGGGTCTCGCGGTATCATTGCAGCACTGGGGCCAGATGGTAAGCCCTCCCGTATCGTAGTTATCTACACGACGGGGAGTCAGGCAACTATGGATGAACGAAATAGACAGATCGCTGAGATAGGTGCCTCACTGATTAAGCATTGGTAACTGTCAGACCAAGTTTACTCATATATACTTTAGATTGATTTAAAACTTCATTTTTAATTTAAAAGGATCTAGGTGAAGATCCTTTTTGATAATCTCATGACCAAAATCCCTTAACGTGAGTTTTCGTTCCACTGAGCGTCAGACCCCGTAGAAAAGATCAAAGGATCTTCTTGAGATCCTTTTTTTCTGCGCGTAATCTGCTGCTTGCAAACAAAAAAACCACCGCTACCAGCGGTGGTTTGTTTGCCGGATCAAGAGCTACCAACTCTTTTTCCGAAGGTAACTGGCTTCAGCAGAGCGCAGATACCAAATACTGTCCTTCTAGTGTAGCCGTAGTTAGGCCACCACTTCAAGAACTCTGTAGCACCGCCTACATACCTCGCTCTGCTAATCCTGTTACCAGTGGCTGCTGCCAGTGGCGATAAGTCGTGTCTTACCGGGTTGGACTCAAGACGATAGTTACCGGATAAGGCGCAGCGGTCGGGCTGAACGGGGGGTTCGTGCACACAGCCCAGCTTGGAGCGAACGACCTACACCGAACTGAGATACCTACAGCGTGAGCTATGAGAAAGCGCCACGCTTCCCGAAGGGAGAAAGGCGGACAGGTATCCGGTAAGCGGCAGGGTCGGAACAGGAGAGCGCACGAGGGAGCTTCCAGGGGGAAACGCCTGGTATCTTTATAGTCCTGTCGGGTTTCGCCACCTCTGACTTGAGCGTCGATTTTTGTGATGCTCGTCAGGGGGGCGGAGCCTATCGAAAAACGCCAGCAACGCGGCCTTTTTACGGTTCCTGGCCTTTTGCTGGCCTTTTGCTCACATGTTCTTTCCTGCGTTATCCCCTGATTCTGTGGATAACCGTATTACCGCCTTTGAGTGAGCTGATACCGCTCGCCGCAGCCGAACGACCGAGCGCAGCGAGTCAGTGAGCGAGGAAGCGGAAGAGCGCCCAATACGCAAACCGCCTCTCCCCGCGCGTTGGCCGATTCATTAATGCAGCTGGCACGACAGGTTTCCCGACTGGAAAGCGGGCAGTGAGCGCAACGCAATTAATGTGAGTTAGCTCACTCATTAGGCACCCCAGGCTTTACACTTTATGCTTCCGGCTCGTATGTTGTGTGGAATTGTGAGCGGATAACAATTTCACACAGGAAACAGCTATGACCATGATTACGCCAAGCTATTTAGGTGACACTATAGAATACTC

**>pSLI-SMC1-GFP_GlmS**

XXXX = SMC1 homology region

XXXX = Linker

XXXX = GFP

XXXX = T2A skip peptide

XXXX = G418 resistance

XXXX = GlmS

ggatatggcagcttaatgttcgtttttcttatttatatatttataccaattgattgtatttataactgtaaaaatgtgtatgttgtgtgcatatttttttttgtgcatgcacatgcatgtaaatagctaaaattatgaacattttattttttgttcagaaaaaaaaaactttacacacataaaatggctagtatgaatagccatattttatataaattaaatcctatgaatttatgaccatattaaaaatttagatatttatggaacataatatgtttgaaacaataagacaaaattattattattattattatttttactgttataattatgtgtctccttcaatgattcataaatagttggacttgatttttaaaatgtttataatatgattagcatagttaaataaaaaaagttgaaaaattaaaaaaaaacatataaacacaaatgatggtttttccttcaatttcgatatcaatttatagaaacaaaatatatacttgtataattttatttttttatataaatcattacatatataattatacaatattttttctaagagataattatatattaatatatataaaaaaaggtgttttttttttttttttttatttttatttttattttatggtaatattttattttccttattttataaattatattagtttatatgtgattaattttatatattatcaatttatatatttttaaatgcttacttaattatctttttttttttttttttttttttttcccctctttttatattaatttatttttgaaaaaattgatatatatatatatatataatatatatatatacatgtagtagtattaaacaatgtataatatatataaataatatatttatatatttcatttcaattttaattttttttggttttttttttttttctttttgtcatatttaaaaaaaattatattcatataagttatgcattttttataaacattattcaatatatgtataatataatatatatatatatattaatgtattattccaatgtgcatgataaaagaaaaaaataatatttataaaaaaaaagaaaaataaaacaaaaaaagaaaaaaaaaaaaaaaaaaaaaaaaatacaaaaataaataatataatttataattatatattcttgtcacaataaaaatatatatatatatatatatatttataatatgtatattttaaactagaaaaggaataactaatattttatttattatcattcaagatttatattttataataataaatacctaatagaaatatatcaggatccatgcatggttcgctaaactgcatcgtcgctgtgtcccagaacatgggcatcggcaagaacggggactacccctggccaccgctcaggaacgaatttagatatttccagagaatgaccacaacctcttcagtagaaggtaaacagaatctggtgattatgggtaagaagacctggttctccattcctgagaagaatcgacctttaaagggtagaattaatttagttctcagcagagaactcaaggaacctccacaaggagctcattttctttccagaagtctagatgatgccttaaaacttactgaacaaccagaattagcaaataaagtagacatggtctggatagttggtggcagttctgtttataaggaagccatgaatcacccaggccatcttaaactatttgtgacaaggatcatgcaagactttgaaagtgacacgttttttccagaaattgatttggagaaatataaacttctgccagaatacccaggtgttctctctgatgtccaggaggagaaaggcattaagtacaaatttgaagtatatgagaagaatgattaagcttatttaataatagattaaaaatattataaaaataaaaacataaacacagaaattacaaaaaaaatacatatgaattttttttttgtaatcttccttataaatatagaataatgaatcatataaaacatatcattattcatttatttacatttaaaattattgtttcagtatctttaatttattatgtatatataaaaataacttacaattttattaataaacaatatatgtttattaattcatgttttgtaatttatgggatagcgattttttttactgtctgtatttttcttttttaattatgttttaattgtattttatttttattattgttctttttatagtattattttaaaacaaaatgtattttctaagaacttataataataataaatataaattttaataaaaattatatttatcttttacaatatgaacataaagtacaacattaatatatagcttttaatatttttattcctaatcatgtaaatcttaaatttttctttttaaacatatgttaaatatttatttctcattatatataagaacatatttattaaatctagaattctatagtgagtcgtattacaattcactggccgtcgttttacaacgtcgtgactgggaaaaccctggcgttacccaacttaatcgccttgcagcacatccccctttcgccagctggcgtaatagcgaagaggcccgcaccgatcgcccttcccaacagttgcgcagcctgaatggcgaatggcgcctgatgcggtattttctccttacgcatctgtgcggtatttcacaccgcatatggtgcactctcagtacaatctgctctgatgccgcatagttaagccagccccgacacccgccaacacccgctgacgcgccctgacgggcttgtctgctcccggcatccgcttacagacaagctgtgaccgtctccgggagctgcatgtgtcagaggttttcaccgtcatcaccgaaacgcgcgagacgaaagggcctcgtgatacgcctatttttataggttaatgtcatgataataatggtttcttagacgtcaggtggcacttttcggggaaatgtgcgcggaacccctatttgtttatttttctaaatacattcaaatatgtatccgctcatgagacaataaccctgataaatgcttcaataatattgaaaaaggaagagtatgagtattcaacatttccgtgtcgcccttattcccttttttgcggcattttgccttcctgtttttgctcacccagaaacgctggtgaaagtaaaagatgctgaagatcagttgggtgcacgagtgggttacatcgaactggatctcaacagcggtaagatccttgagagttttcgccccgaagaacgttttccaatgatgagcacttttaaagttctgctatgtggcgcggtattatcccgtattgacgccgggcaagagcaactcggtcgccgcatacactattctcagaatgacttggttgagtactcaccagtcacagaaaagcatcttacggatggcatgacagtaagagaattatgcagtgctgccataaccatgagtgataacactgcggccaacttacttctgacaacgatcggaggaccgaaggagctaaccgcttttttgcacaacatgggggatcatgtaactcgccttgatcgttgggaaccggagctgaatgaagccataccaaacgacgagcgtgacaccacgatgcctgtagcaatgccaacaacgttgcgcaaactattaactggcgaactacttactctagcttcccggcaacaattaatagactggatggaggcggataaagttgcaggaccacttctgcgctcggcccttccggctggctggtttattgctgataaatctggagccggtgagcgtgggtctcgcggtatcattgcagcactggggccagatggtaagccctcccgtatcgtagttatctacacgacggggagtcaggcaactatggatgaacgaaatagacagatcgctgagataggtgcctcactgattaagcattggtaactgtcagaccaagtttactcatatatactttagattgatttaaaacttcatttttaatttaaaaggatctaggtgaagatcctttttgataatctcatgaccaaaatcccttaacgtgagttttcgttccactgagcgtcagaccccgtagaaaagatcaaaggatcttcttgagatcctttttttctgcgcgtaatctgctgcttgcaaacaaaaaaaccaccgctaccagcggtggtttgtttgccggatcaagagctaccaactctttttccgaaggtaactggcttcagcagagcgcagataccaaatactgtccttctagtgtagccgtagttaggccaccacttcaagaactctgtagcaccgcctacatacctcgctctgctaatcctgttaccagtggctgctgccagtggcgataagtcgtgtcttaccgggttggactcaagacgatagttaccggataaggcgcagcggtcgggctgaacggggggttcgtgcacacagcccagcttggagcgaacgacctacaccgaactgagatacctacagcgtgagctatgagaaagcgccacgcttcccgaagggagaaaggcggacaggtatccggtaagcggcagggtcggaacaggagagcgcacgagggagcttccagggggaaacgcctggtatctttatagtcctgtcgggtttcgccacctctgacttgagcgtcgatttttgtgatgctcgtcaggggggcggagcctatcgaaaaacgccagcaacgcggcctttttacggttcctggccttttgctggccttttgctcacatgttctttcctgcgttatcccctgattctgtggataaccgtattaccgcctttgagtgagctgataccgctcgccgcagccgaacgaccgagcgcagcgagtcagtgagcgaggaagcggaagagcgcccaatacgcaaaccgcctctccccgcgcgttggccgattcattaatgcagctggcacgacaggtttcccgactggaaagcgggcagtgagcgcaacgcaattaatgtgagttagctcactcattaggcaccccaggctttacactttatgcttccggctcgtatgttgtgtggaattgtgagcggataacaatttcacacaggaaacagctatgaccatgattacgccaagctatttaggtgacactatagaatactcgcggccgcatacatgaacatgatgaagaagaacaaaataacatacttttatcgaatatatcattcgatcttttatcagatgatttaaaaaatatggaaaatgataaagatataaacaatgaaaaagaaaatatggaaaaagaaattgaaagaaaaaaaaaactgttaaaattaaaaaatgtaaattgtaatgctgaaaaagaatatgaaaaattaacatctaaattaaaatcaattgatgtatcgctttctgaagaaagaaaagaatgtaatttatttgaacgtaattttagaatattacaaaaaaaaagatcttacaaattcttacattgttttaattatattaaaaatattatagataatgtatataataatttaacatataatgctaaacatcatgtcggtggacaagcctttttagacttatgtaattttaatgaatttaataaagatgatgaacctttctattgtggtatcaaatataataatatgcctccaatgaaaagatattttgaaatatctgaattaagtggaggtgaaaaaagtattagtgcattagcattaattttctctattcaaaaatatattaataattcatttattatacttgacgaagtggatgcaaatatggaccctctcaaaatccagtctctaacaagatacctaaattcaattaatagtcaagttattgttatttctttgaaggaaaaattctttagtaaaagtcaatcattaatcggagtgtacaaaaataaacataaaaagtgttcaaaaacaattactttggatataagtaaatatcgacaggatgtaccaaacaatacgcgtgctagaggtgctgctgctggtgctggaggtgcaggtagacgtacgatgagtaaaggagaagaacttttcactggagttgtcccaattcttgttgaattagatggtgatgttaatgggcacaaattttctgtcagtggagagggtgaaggtgatgcaacatacggaaaacttacccttaaatttatttgcactactggaaaactacctgttccatggccaacacttgtcactactttcgcgtatggtcttcaatgctttgcgagatacccagatcatatgaaacagcatgactttttcaagagtgccatgcccgaaggttatgtacaggaaagaactatatttttcaaagatgacgggaactacaagacacgtgctgaagtcaagtttgaaggtgatacccttgttaatagaatcgagttaaaaggtattgattttaaagaagatggaaacattcttggacacaaattggaatacaactataactcacacaatgtatacatcatggcagacaaacaaaagaatggaatcaaagttaacttcaaaattagacacaacattgaagatggaagcgttcaactagcagaccattatcaacaaaatactccaattggcgatggccctgtccttttaccagacaaccattacctgtccacacaatctgccctttcgaaagatcccaacgaaaagagagaccacatggtccttcttgagtttgtaacagctgctgggattacacatggcatggatgagctctacaaagtcgacggagaaggaagaggaagtttattaacatgtggagatgtagaagaaaatccaggaccaatgattgaacaagatggattgcacgcaggttctccggccgcttgggtggagaggctattcggctatgactgggcacaacagacaatcggctgctctgatgccgccgtgttccggctgtcagcgcaggggcgcccggttctttttgtcaagaccgacctgtccggtgccctgaatgaactgcaggacgaggcagcgcggctatcgtggctggccacgacgggcgttccttgcgcagctgtgctcgacgttgtcactgaagcgggaagggactggctgctattgggcgaagtgccggggcaggatctcctgtcatctcaccttgctcctgccgagaaagtatccatcatggctgatgcaatgcggcggctgcatacgcttgatccggctacctgcccattcgaccaccaagcgaaacatcgcatcgagcgagcacgtactcggatggaagccggtcttgtcgatcaggatgatctggacgaagagcatcaggggctcgcgccagccgaactgttcgccaggctcaaggcgcgcatgcccgacggcgaggatctcgtcgtgacccatggcgatgcctgcttgccgaatatcatggtggaaaatggccgcttttctggattcatcgactgtggccggctgggtgtggcggaccgctatcaggacatagcgttggctacccgtgatattgctgaagagcttggcggcgaatgggctgaccgcttcctcgtgctttacggtatcgccgctcccgattcgcagcgcatcgccttctatcgccttcttgacgagttcttctaactcgagtaattatagcgcccgaactaagcgcccggaaaaaggcttagttgacgaggatggaggttatcgaattttcggcggatgcctcccggctgagtgtgcagatcacagccgtaaggatttcttcaaaccaagggggtgactccttgaacaaagagaaatcacatgatctctcgag
